# Quantifying microbial fitness in high-throughput experiments

**DOI:** 10.1101/2024.08.20.608874

**Authors:** Justus Wilhelm Fink, Michael Manhart

## Abstract

Few concepts are as central to evolution as is fitness, and yet the quantification of fitness is often ambiguous. In particular, high-throughput experiments to measure mutant fitness in microbes are increasingly common but vary widely in their definitions of fitness, which makes their results difficult to compare. What are the consequences of these different fitness statistics, and is there a best way to quantify fitness in a given context? Here we systematize the set of possible fitness statistics according to the following three choices: 1) the encoding of relative abundance (e.g., transforming by a log or logit function), 2) the time scale over which to measure the change in relative abundance, and 3) the choice of a reference subpopulation for calculating relative fitness in bulk competition experiments, such as those using DNA-barcoded mutants. We show that these different choices can lead to significantly different interpretations of mutant fitness, affecting the magnitude of fitness effects, the apparent presence of epistasis, and even the fitness ranking across mutants. This can confound predictions for evolutionary dynamics and gene functions. Altogether our results demonstrate the importance of consistent fitness definitions for reproducible results across experiments.

## INTRODUCTION

The concept of fitness is supposed to describe the fate of mutations as they arise in a population [1] and defines other core evolutionary concepts such as tradeoffs [2]. At the same time, fitness is often a confusing term in evolutionary biology [3–5]. It is possible to classify fitness measures by their role in theory [6] and their scale of measurement [7–9], but these arguments offer little guidance on how to measure fitness in practice. Empirical fitness measurements have become widespread in the microbial sciences — to test evolutionary theory [10–15], detect microbial interactions [16, 17], annotate gene functions [18, 19], and understand the spread of antibiotic resistance genes [20, 21] — but the approaches to quantifying fitness in these studies vary widely.

The classic approach to measuring the fitness effects of mutations uses pairwise competition experiments, where the mutant competes with its ancestral wild-type in co-culture [22–24]. This approach is ideal because it mimics the dynamics of spontaneous mutations, and thus is typically used in experimental evolution [10, 12, 25–27]. For example, the Long-Term Evolution Experiment (LTEE) has evolved *Escherichia coli* over tens of thousands of generations and measured the fitness of each evolved lineage in competition with the ancestor [11, 26, 28]. However, pairwise competition experiments require a way to distinguish mutants from the wild-type in cocultures (e.g., fluorescent labeling or selection markers), which is not always feasible to engineer. A second approach is thus to measure properties of genotypes in monocultures, like the growth rate, and estimate relative fitness between genotypes from these measurements. Monoculture growth curves are straightforward to measure, even for large numbers of mutants [29–31], but empirical tests have shown that the monoculture growth rate is insufficient to predict the relative fitness between genotypes [32–34]. Summary statistics of whole growth curves, like the area under the curve (AUC), perform better [35–37]. However, this approach breaks down for strains that engage in cross-feeding or toxin production, and attempts to improve monoculture-based fitness estimates require additional experiments [34, 37] and do not go beyond qualitative fitness rankings [37].

A third and more recent approach is to measure relative fitness in bulk competition experiments using high-throughput sequencing, often of DNA barcodes [13, 38]. For example, transposon-insertion mutagenesis generates libraries of gene knockouts that can be tracked as they grow in a single batch culture [39]. Estimating the fitness of each mutant allows us to identify genes that are particularly important in the growth environment [18, 19, 40], in the given genetic background [14, 15, 41, 42], and in the context of an ecological community [16, 17, 42, 43]. Bulk fitness estimation also plays a major role in identifying beneficial lineages in experimental evolution [13, 44] (not unlike the tracking of viral pathogens in public health [45–47]) and especially testing their fitness in new environments [48–52]. Many of these studies make different choices about the design of experiments and the calculation of fitness, without explaining why a certain fitness measure was used or how to compare between data sets.

The inconsistent choices for quantifying microbial fitness make it is difficult to compare fitness across experiments, especially because different groups tend to use different definitions. For example, the LTEE reports relative fitness per-generation [11, 26], whereas other evolution experiments exclusively report relative fitness per-cycle of batch growth [12, 53]. New techniques, like barcode sequencing, often spark their own fitness metrics [18, 40, 54–56]. A different choice of fitness metric might lead to a different ranking of mutants in a genetic screen (and the importance we assign to those genes) or change statistical properties of the distribution of mutant fitness effects, which is essential to predict the probability of mutant fixation [57], the speed of adaptation [58], and the effect size in statistical tests of gene function [18, 19]. This raises three key questions: 1) How do these fitness statistics differ? 2) Can we say which choices are optimal, in the light of first principles or some practical considerations? 3) Does this tell us how we should design these experiments? Previous work has addressed aspects of these questions [6–9, 59] but the arguments are scattered across disciplines and based on generic models of population growth [6, 7, 9, 60], rather than incorporating the specific dynamics of microbes in laboratory experiments. Here we address these questions from a unified framework, using realistic microbial population dynamics with empirical traits to systematically test different fitness metrics.

## RESULTS

### Predictive power of fitness depends on the encoding of relative abundance

We distinguish between three related but distinct notions of fitness [6], all based on predicting dynamics of a population [61–64]. The first type of fitness is *absolute fitness*, which is a property of a single genotype by itself and serves to predict the change in the genotype’s absolute abundance *N* (*t*) over a future time window Δ*t* (Fig. S1A). This is important for questions about extinction and evolutionary rescue [65]. The second type of fitness is *relative fitness*, which is a property of two genotypes as it describes how the relative abundance *x*(*t*) of one genotype changes compared to the other over a time Δ*t* (Fig. S1B). This is important to determine the fixation probability of new mutations [57, 66, 67]. In general these dynamics are stochastic [66, 68, 69], but throughout this paper we focus on their average behavior across replicate cultures (as sketched in Fig. S1C).

A practical challenge of working with relative fitness is that it must be measured between all pairs of genotypes in coculture competitions. Therefore it is common to infer relative fitness of two genotypes based on some individual properties of the genotypes [70–73]. We denote this third notion of fitness as the *fitness potential*. Like absolute fitness, fitness potential is a property of an individual genotype, but unlike the absolute fitness, it has no meaning by itself. In analogy with the potential energy of physical systems, fitness potential is intended only to be compared between two genotypes (e.g., by taking the difference or ratio) to estimate the relative fitness between those genotypes [36, 37]. The collection of fitness potential values across a large set of genotypes forms a fitness landscape [74, 75]. Note that the term “fitness potential” has been used with a related but somewhat different meaning in the population genetics literature [76–78].

We emphasize that fitness as defined here gives information about short-time dynamics but not necessarily the long-term outcome (compare Smith et al. [6]). For example, this excludes the ratio of growth rates [7, 73] or the resource concentration *R* ^∗^ in chemostat equilibrium [79] since these quantities cannot tell you how fast the absolute or relative abundance is changing. We also note that in all these definitions, the fitness of a genotype is specific to one environment, including all abiotic and biotic factors. How fitness in one environment is related to fitness in other environments is important [14, 19, 42, 43, 49, 50, 73, 80, 81] but beyond the scope of this article, as we focus on specific laboratory measurements of fitness rather than abstract concepts.

In this article we will focus on relative fitness as this is the most common object of high-throughput laboratory experiments in microbes [18, 26, 40, 55, 56] and also has the most straightforward connection to key evolutionary outcomes such as fixation [57, 66, 67]. We define a genotype’s relative fitness as any number that is sufficient to predict the genotype’s relative abundance *x*(*t*) over a short time period (Fig. S1). The simplest approach is to take a linear expansion of *x*(*t*) and use its slope at time *t* to predict the change over a time Δ*t* (e.g., Fig. 1A, top panel):

**FIG. 1.**
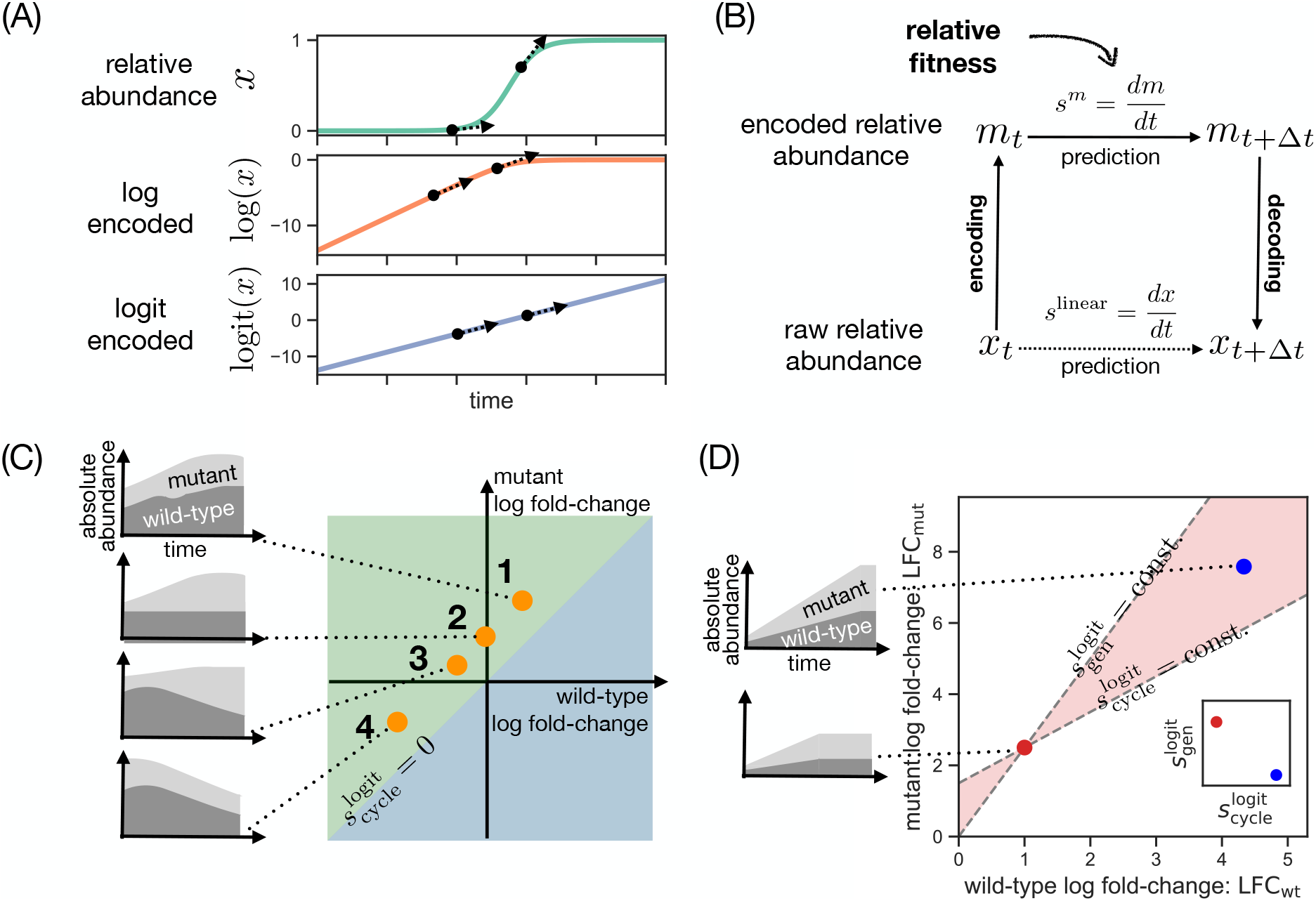
Overview of the choice of encoding and the choice of time scale for quantifying relative fitness. (A) Example trajectory of relative abundance *x* (top panel) for a mutant invading and eventually replacing a wild-type population. The same trajectory is plotted under the encoding log *x* (middle panel) and the encoding logit *x* = log(*x/*(1 − *x*)) (bottom panel). (B) Flowchart to predict the future relative abundance of a mutant given a relative fitness value *s*^*m*^ = *dm/dt* for some encoding *m*. The current relative abundance *x*_*t*_ is transformed into the new variable *m*_*t*_ = *m*(*x*_*t*_), then projected into the future through a linear extrapolation using *s*^*m*^ (upper horizontal arrow) and finally converted back into a frequency *x*_*t*+!*t*_ using the decoding function *m*^−1^. (C) Four scenarios for a mutant with the same relative fitness per-cycle but with different underlying population dynamics. For each scenario, we show an example trajectory of absolute abundance (stacked) for the wild-type (dark grey) and mutant population (light grey). Each scenario is mapped as a single-dot onto the fold-change diagram (center plot) and colored areas indicated positive (green area) and negative relative fitness per-cycle (blue area; compare Eq. (9)). Fitness per-cycle has isoclines that are parallel to the identity in the LFC diagram. (D) Scenario for rank discrepancy between relative fitness per-cycle 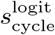 and relative fitness per-generation 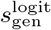. For a given competition (red dot), rank discrepancy occurs in a bow tie-shaped area of the fold-change space (shaded red). Any competition in the right half of this area (blue dot) will have higher mutant fitness 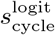 but lower mutant fitness 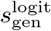 (right inset). As small plots on the left, we show possible population dynamics that generate this fold-change variation. Here the growth rates of the mutant and wild-type are identical in the two scenarios, but the growth period is different (e.g., from additional resources).

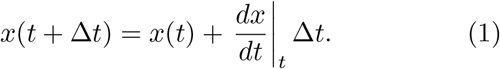

The slope

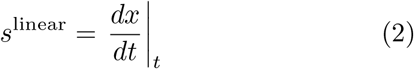

qualifies as a fitness statistic but often will incorrectly estimate the actual change in relative abundance because *x*(*t*) is nonlinear (Fig. 1A, top panel).

Alternatively, we can *encode* the relative abundance *x* in a new variable *m*(*x*) (any smooth, strictly-increasing function) whose dynamics may be easier to linearly approximate (Fig. 1B; compare Denniston and Crow [82]):

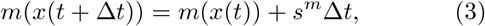

where

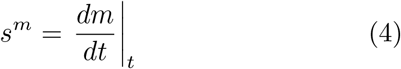

is the relative fitness under the encoding *m*. In Sec. S1 we show examples of the encoded relative fitness for an explicit model of population dyanmics, and in Sec. S2 we show how this concept generalizes to absolute fitness as well.

The ideal encoding transforms the relative abundance *x*(*t*) into a linear trajectory that can be fully predicted from the slope *s*^*m*^. While the log-encoding *m*(*x*) = log *x* is common in many TnSeq studies [17–19, 43, 80, 81] (and is implicit in plots of relative abundance on a logarithmic scale [44]), the log-encoded fitness *s*^log^ tends to overestimate the change at high relative abundance (Fig. 1A, middle panel). Instead, the best encoding in most cases is the logit function (Fig. 1A, bottom panel):

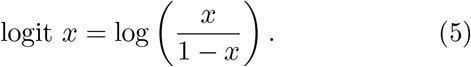

This is because the relative abundance *x*(*t*) typically follows logistic dynamics (Sec. S1), at least as a first-order approximation (Sec. S3, Fig. S2), and the logit function is linearly related to the inverse function of logistic *x*(*t*). The logit-encoded relative fitness of a mutant is equivalent to the standard definition of its selection coefficient in population genetics [83]. For this reason we focus on the logit-encoded relative fitness for the remainder of this work. Note that the logit encoding and logistic dynamics apply to relative abundance and are not to be confused with the logistic growth model used with microbial growth curves, where it describes the absolute abundance [84].

### Estimating relative fitness requires a choice of time scale

Since the relative abundance of a genotype can only be measured at discrete times, we must estimate the instantaneous relative fitness of Eq. (4) using a finite difference between consecutive time points (Fig. S1C, Sec. S4):

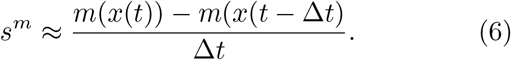

For the logit-encoding this becomes

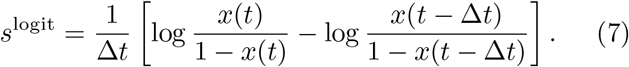

In the case of a single mutant competing against a wild-type strain, we can rewrite Eq. (7) in terms of the log fold-changes of the mutant and wild-type strains:

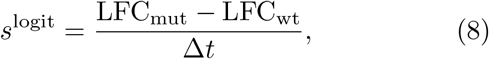

where the log fold-change of each strain is LFC = log *N* (*t*)*/N* (*t* − Δ*t*) and *N* is the biomass of the strain. The LFC is sometimes also referred to as a Malthusian parameter [9, 26, 65].

Therefore a key element of estimating relative fitness from discrete time data is the choice of time interval Δ*t*. There are two common choices in empirical measurements of microbial fitness. The first is to use a time scale extrinsic to the population. In particular, microbial populations in laboratory experiments [26, 85] but also in some natural environments [86−88] grow in discrete growth cycles dictated by external pulses of nutrients (e.g., batch culture). In this case, it is convenient to quantify relative fitness *per-cycle*, by choosing time units in Eq. (8) such that Δ*t* = 1:

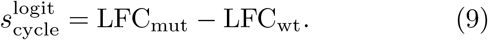

The second common choice for Δ*t* is a time scale intrinsic to the population, typically its generation time. The relative fitness per-cycle may depend on the number of generations in the growth cycle, and it can be valuable to normalize for this dependence, especially when comparing across environments [26]. However, defining the number of generations for a population with multiple genotypes growing at different rates is ambiguous. In the case of a single mutant competing with a wild-type, it is common to consider only the number of generations experienced by the wild-type strain, which is estimated by its log fold-change LFC_wt_. It is convenient to express the generations in base *e* to match the natural logarithm in the logit-encoding, but this could equivalently be done by converting all logarithms in Eq. (8) to base 2. Thus the relative fitness of a mutant *per-generation* is defined by choosing Δ*t* = LFC_wt_ in Eq. (8):

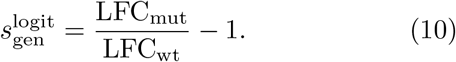

This quantity is equivalent to the fitness statistic 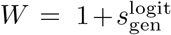 used in the LTEE [26] and other studies [89–93]. Some authors assume a fixed number of generations per growth cycle [12, 15], but since this may not be true, we use the term “per-generation” strictly for fitness statistics where the wild-type generations are explicitly measured.

### Relative fitness per-cycle and per-generation can rank mutants differently when wild-type growth depends on mutant

Although relative fitness of a mutant per growth-cycle (Eq. (9)) and per-generation (Eq. (10)) differ only by choice of time scale, they are not just equivalent quantities in different units. First, while fitness per-cycle is well-defined regardless of whether the population undergoes net growth or death (Fig. 1C), fitness per-generation is not defined when the wild-type undergoes no change or net death (denominator of Eq. (10) is not positive) [5, 94]. This can occur for microbes under high drug concentrations, as well as for microbial populations under harsh environmental conditions as found in sediments or wastewater [95, 96].

However, even when the wild-type LFC is positive, the relative fitness per-cycle and per-generation are not necessarily proportional to each other across a set of mutants competed against the same wild-type. The reason is that while the growth cycle time scale is extrinsic to a population and therefore the same across all pairs of genotypes, the generation time scale depends on the growth of the wild-type, which may be affected by the specific mutant it is competing against. This discrepancy can be quantitative as well as qualitative, producing different rankings of mutants. For example, Fig. 1D shows two mutant genotypes (red and blue) that have opposite rankings under these fitness statistics: the blue mutant has higher relative fitness per-cycle 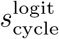, but the red mutant has higher relative fitness per-generation 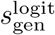. The disagreement in ranking requires positive covariation in the LFCs (red bow tie-shaped area in Fig. 1D), such that the mutant with higher LFC also induces a higher LFC in the wild-type (see illustration in Fig. and exact conditions in Sec.S5)

### Basic features of microbial population dynamics are sufficient to cause rank discrepancy between fitness statistics

Since the discrepancy between fitness per-cycle 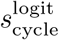 and per-generation 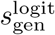 occurs when the wild-type LFC varies across pairwise competitions with different mutants, we need to determine what biological mechanisms cause this. One way to explore this is to use a minimal model of microbial population dynamics parameterized by empirical data on a set of mutants. While a minimal model does not capture all possible mechanisms, it provides a null expectation for how common these discrepancies should be in real populations.

We use a model that approximates microbial population dynamics as three growth phases — a lag phase, an exponential growth phase, and a stationary phase (Fig. 2A) — where strains compete for a single limiting resource with no other interactions [97, 98], since resource competition is likely to be the main interaction between a wild-type and a mutant with only small genetic differences (e.g., a point mutation or gene knockout). This model captures the minimal dynamics of microbial growth under laboratory conditions, and therefore serves as a baseline for possible outcomes. We parameterize the model using growth curves of single-gene knockout mutants in *Saccharomyces cerevisiae* [29, 99] (Figs. and S5, Sec. S6). We simulate a pairwise competition for each knockout against the wild-type (Fig. 2A) and calculate relative fitness per-cycle 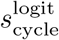 and per-generation 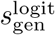 for each knockout (Methods). While the relative fitness per-cycle and per-generation in the model are highly correlated across mutants overall (Fig. S6), the model predicts major differences in individual mutants when we rank them by fitness (Fig. 2B). Indeed, when we plot all simulated mutant competitions on the LFC diagram, we find that there are many pairs of points that have positive covariation in the mutant and wild-type LFC and thus give rise to rank differences (compare Figs. 2C and 1D). Since these rankings often serve to select important mutants from a functional genomics screen or an evolution experiment, discrepancies between fitness definitions may lead analyses down different paths.

**FIG. 2.**
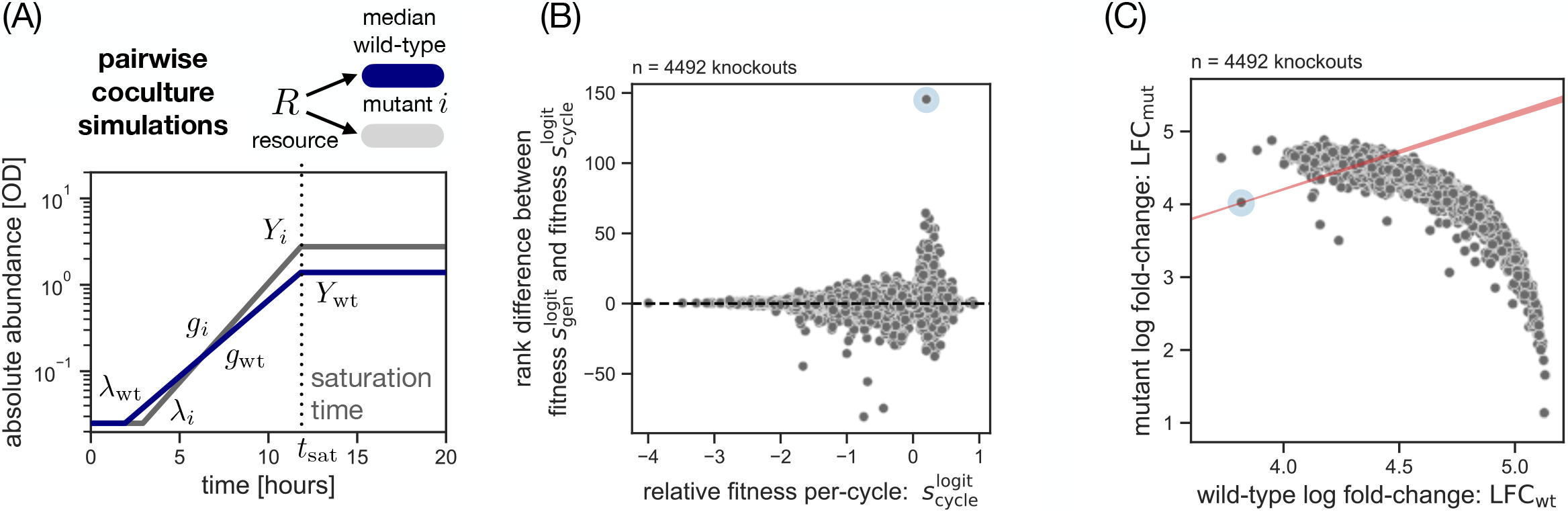
Comparison of mutant fitness rankings with different time scales using empirical trait variation. (A) Overview of pairwise coculture simulations. For each mutant strain (grey), we simulate a competition growth cycle against a reference wild-type strain (blue) using the estimated traits and laboratory parameters for the initial condition (*N*_0_ = 0.05 OD, *R*_0_ = 111 mM glucose, *x* = 0.5; Methods and see Fig. for the underlying trait distribution). (B) Rank discrepancy between relative fitness per-cycle 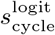 and per-generation 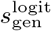. Higher rank is defined as higher fitness, and the rank difference is defined as the rank in 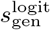 minus the rank in 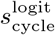. The blue halo marks the mutant with the greatest rank difference. (C) Covariation between wild-type and mutant fold-change across all simulated competitions for all mutant strains (grey dots). The blue halo marks the same mutant as in panel B, with its red bow tie-shaped area defining the space of LFCs that would have a rank discrepancy with it (compare Fig. 1D), which includes many other mutant competitions.

This model therefore shows that the basic features of microbial population dynamics and resource competition with realistic parameters are sufficient to cause substantial disagreement between fitness per-cycle and per-generation, underscoring the importance of this analysis choice. Furthermore, the model reveals a basic mechanism that causes this discrepancy: mutants must have diminished biomass yield plus variation in one additional trait (lag time or growth rate), and a high mutant relative abundance (e.g., 50%) at the start of the competition (Fig. S7). These conditions are necessary because they allow mutants both to have variation in fitness and to alter the wild-type LFC in pairwise competition. Even though the wild-type growth rate is unaffected by mutants, the wild-type LFC will change in the presence of mutants that have lower yield because the population on average consumes more resources per cell division (Sec. S7). By this mechanism, it is possible to concoct a set of growth traits where relative fitness per-generation and per-cycle deliver completely opposite rankings (Fig. S8).

### Fitness statistics can disagree on epistasis and long-term fitness trends

Since the relative fitness of single mutants differs depending on whether fitness is quantified per-cycle or per-generation, how does that choice impact the apparent epistatic interactions between multiple mutations and the long-term evolutionary trend in fitness? Using the same minimal model of population dynamics as in Fig. 2A, we simulate a set of six mutants, each one having an increase or decrease in each of the three quantitative traits in the model (lag time, growth rate, and yield), and calculate the relative fitness per-cycle and per-generation for all pairs of those mutations (assuming the mutation effects on the traits themselves are additive; Methods). We quantify epistasis as the fitness difference (in 1:1 competition with the wild-type; Methods) between the double mutant and the sum of the single mutants (Fig. 3A) [100].

**FIG. 3.**
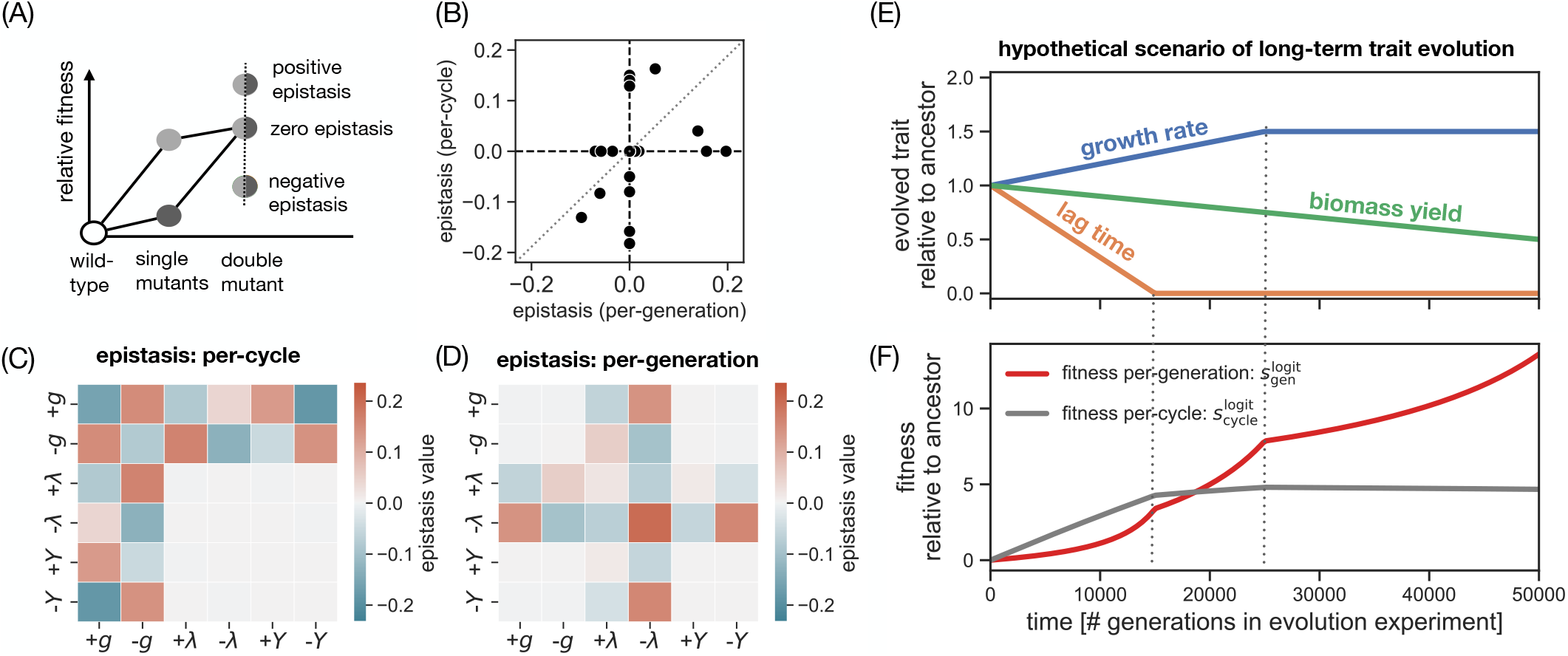
Discrepancy between fitness per-generation and per-cycle in epistasis detection and long-term trend. (A) Schematic detection of epistasis for two single mutants (light and dark circle). Epistasis occurs if the corresponding double mutant (two-component circles) deviates from the additive fitness (parallelogram shape in solid lines; Methods). (B) Correlation between epistasis in relative fitness per-cycle *s*_cycle_ and per-generation *s*_gen_. Each dot corresponds to a hypothetical double mutant that combines pairs of mutations systematically perturbing each of the three growth traits (Fig. 2A) in our minimal model of population dynamics (Methods). (C) Heatmap for epistasis in the per-cycle fitness *s*_cycle_ for all double mutants in our simulated data set, same data as in panel B. The rows and columns correspond to the six single mutations in our model of population dynamics (Methods): an increase or decrease in lag time (+*ω*, − *ω*), an increase or decrease in maximum growth rate (+*g*, − *g*), or an increase or decrease in biomass yield (+*Y*, −*Y*). (D) Heatmap for epistasis in the per-generation fitness *s*_gen_, same data as in panel B. All epistasis plots are based on 1:1 competition growth cycles with the wild-type (Figs. –). See Fig. for epistasis patterns at low mutant relative abundance. (E) Hypothetical scenario for trait evolution in an evolution experiment. An evolving population decreases lag time to zero (orange line), increases growth rate until saturation (blue line), and gradually decreases biomass yield (green line). This trend is similar to initial observations from the LTEE [101]. (F) Corresponding long-term trend in relative fitness based on the trait evolution in panel E. We estimate relative fitness per-cycle 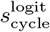 (grey line) and per-generation 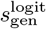 (red line) every 250 generations. Dotted vertical lines mark the end of trait evolution in lag time and growth rate in panel E. For the actual fitness trend in the LTEE, see Fig.

Figure 3B shows that almost all the double mutants differ in their apparent epistasis per-cycle vs. per-generation. Epistasis per-cycle occurs when one of the mutations affects growth rate (Fig. 3C), while epistasis per-generation occurs when one of the mutations affects lag time (Fig. 3D). This is because the effect of a change in growth rate accrues per-generation (and hence is non-epistatic when measured over that time scale; Sec. S8), while the effect of a change in lag time accrues once per-cycle (and hence is non-epistatic in the per-cycle statistic). See Figs. S9–S14 for growth curves in all cases. As an example, the double mutant in Fig. S12, row D has two lag time-reducing mutations that are perfectly additive in the per-cycle statistic. However, the lag time mutations also reduce the saturation time of the growth cycle such that the fitness of the double mutant is spread over a fewer number of wild-type generations LFC_wt_, creating super-additive epistasis in the per-generation statistic. Note that this depends on measuring fitness for a mutant at relatively high abundance in the pairwise competition with the wild-type; when the mutant is rare, the wild-type fold-change LFC_wt_ is effectively constant across mutants and epistasis measured per-cycle is proportional to its value measured per-generation (Fig. S15). Differences in apparent epistasis can also lead to differences in apparent long-term trends of fitness per-cycle and per-generation in an evolution experiment. For example, if growth traits of a population in the minimal model evolve according to Fig. 3E, then fitness per-cycle will saturate at an early time while fitness per-generation will continue to increase (Fig. 3F). This is because the long-term decrease in yield reduces the wild-type LFC in 1:1 competitions between an evolved mutant and the wild-type, which causes fitness per-generation to increase (Eq. (10)).

### Different fitness time scales agree on LTEE trend but not on mutant ranking

As an empirical test for discrepancies between relative fitness per-generation and per-cycle, we test how the choice of fitness time scale affects conclusions from the LTEE (Sec. S9). We reanalyze competition data from the LTEE [11] using the relative fitness per-generation 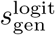 (equivalent to the original definition used in those studies, 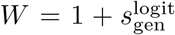) and relative fitness per-cycle 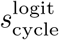. Note these competitions are between the ancestral wild-type and each of the 12 evolved populations, rather than single mutants, across time points. We confirm that the long-term fitness trend in the LTEE is the same for 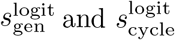 (Fig. S16). When we look more closely at the fitness ranking between lineages, we find that the two statistics disagree on the fitness ranking at almost every time point (Fig. 4A) and the fittest population overall (Fig. 4B; see Fig. S17A for rankings). This discrepancy arises from positive covariation across pairs of LFC values in the LTEE competition data set (Fig. S17B).

**FIG. 4.**
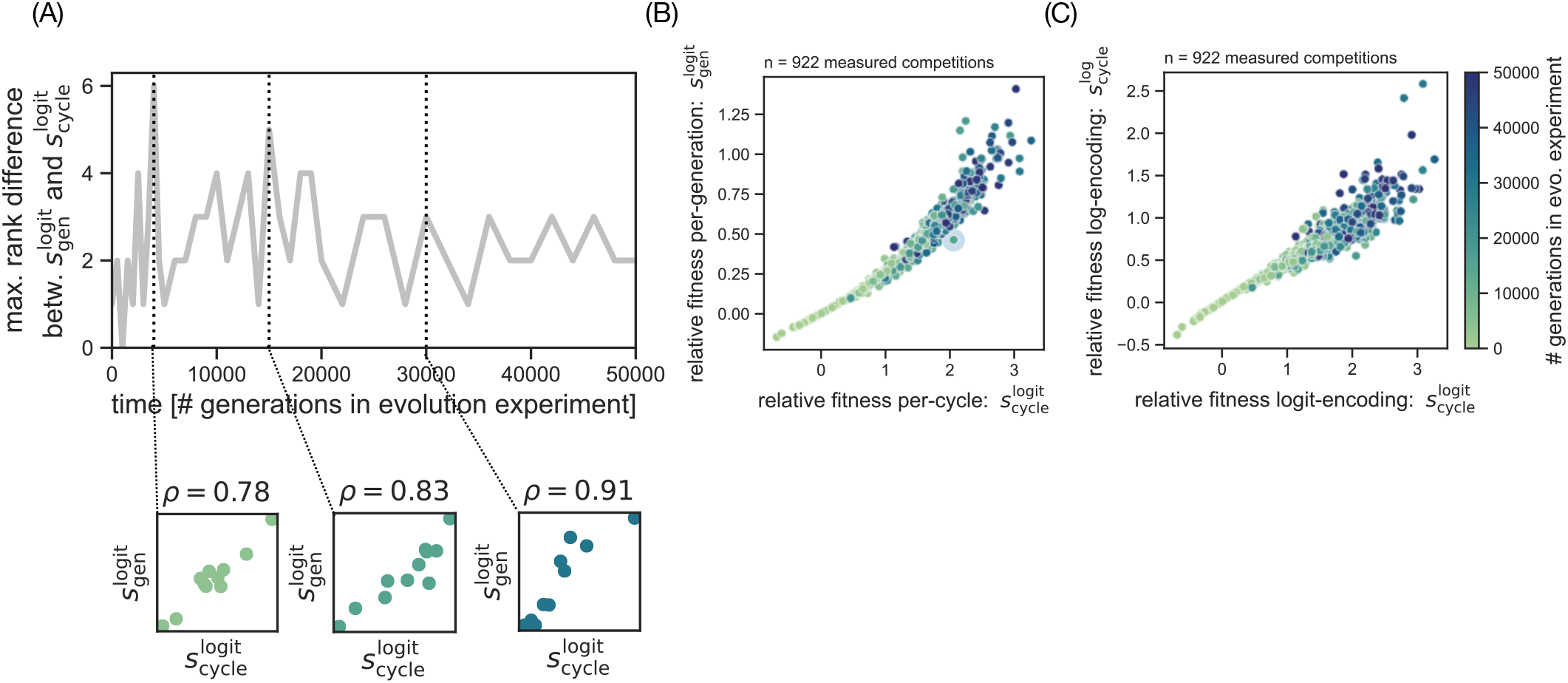
Rank discrepancies between fitness statistics in LTEE competition data. (A) Rank difference between relative fitness per-cycle and per-generation as a function of time in the LTEE. For each of the 12 lines in the LTEE, we compute the relative fitness per-cycle 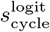 and per-generation 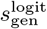 as a function of time, by averaging the data from the LTEE competition data set [11] across replicate measurements (Sec. S9). For some lines, the time-series is truncated due to measurement difficulties [11]. Based on the quantitative fitness values 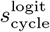 and 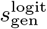, we compute fitness rankings between the replicate lines at any given time point (higher rank means higher fitness; compare Fig. 2B). The rank difference is defined as the rank in 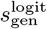 minus the rank in 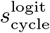, and we combine the rank difference between all pairs of evolving lines to compute the maximum rank difference at each time-point. Insets: At three chosen time-points (*t* = 4000, *t* = 15000, *t* = 30000 generations), we show the correlation between relative fitness per-cycle 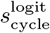 and relative fitness per-generation 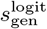 (each dot corresponds to one of the 12 lines in the LTEE). We quantify the correlation between 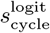 and 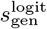 with the Spearman rank correlation *ε* (panel title). Colors indicate the time-point and correspond to the color bar in panel B. (B) Covariation between relative fitness per-cycle 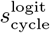 and per-generation 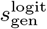 for evolved populations of the LTEE. Each line of the LTEE contributes roughly two competitions per time-point due to replicate measurements (Sec. S9) [11]. The color of the points indicates evolutionary time point; see panel (C) for the color bar. We highlight the evolved population with the greatest rank difference (blue halo; same as in Fig. S17). (C) Same as panel B but comparing relative fitness under the logit-encoding 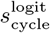 and under the log-encoding 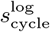.

While we have focused on the logit encoding so far since logistic dynamics are usually the best null model of mutant frequency dynamics (Fig. 1A), the log encoding is common in many TnSeq studies [17–19, 43, 80, 81]. The LTEE competition data is a valuable opportunity to empirically test the consequences of this choice as well. Figure 4C shows that the relative fitness per-cycle under the log-encoding 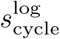 differs from the logit-based fitness 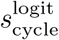 for evolved LTEE populations, leading to ranking discrepancies especially for the fittest populations (Fig. S18A). The discrepancy in fitness between encodings has a different mechanism compared to the discrepancy from fitness time scale: it arises from variation in the initial relative abundance in mutant-to-wild-type competitions in the LTEE data set. Although these pairwise competitions are intended to start with equal abundances of the evolved population and of the wild-type (1:1), Fig. S18B shows that there is a systematic trend toward lower initial frequencies of the evolved populations over time. This trend is a side effect of the inoculation method in LTEE competition experiments, where a fixed volume of the evolved population is transferred to the competition growth cycle but the cell density for the evolved population in pre-culture decreases [101, 102].

### Fitness in bulk competitions is distorted by higher-order effects depending on reference subpopulation

So far we have focused on relative fitness in pairwise competition experiments, but this is often not practical for large numbers of mutants due to the required strain labeling (e.g., with fluorescent proteins or selection markers). One alternative is to measure a fitness potential (as defined at the beginning of the Results section), such as growth rate or area under the growth curve (AUC) [32, 34, 36, 37], from monoculture experiments, under the assumption that true relative fitness is proportional to the difference in this quantity between genotypes. Using our minimal model of population dynamics parameterized by single-gene knockouts in yeast (Fig. 2A), we find that common fitness potentials perform poorly overall (Fig. S19; see refs. [32, 34, 36, 37] for empirical tests), with some success for the AUC but strongly depending on the choice of time window (Fig. S20, Sec. S10).

Another approach to measuring relative fitness for a large number of mutants is to use bulk competition experiments, where many mutant genotypes compete simultaneously in a single culture and each genotype is tracked through high-throughput sequencing [39, 44]. This raises the question of how well relative fitness of a mutant in bulk competition corresponds to its relative fitness in a pairwise competition with the wild-type, since the growth in bulk might be influenced by the presence of other mutant genotypes, a phenomenon known as a higher-order interaction [81, 98, 103].

An important choice in bulk competition experiments is the relative abundance of the library of all mutant genotypes compared to the wild-type. The invasion of a spontaneous mutation into an existing population is best captured by pairwise competitions with low initial relative abundance of the mutant (Case I, Fig. 5A), and one way to recreate this scenario in a bulk competition is to use a low relative abundance of the mutant library overall (Case II, Fig. 5A) [40, 48–52]. A practical problem with Case II is that individual genotypes in the library will have low absolute abundances, which leads to stochasticity in the population dynamics and the sequencing process [13, 104, 105]. Therefore, it is common to compete the mutant library by itself (Case III, Fig. 5A) [13, 15, 17–19, 80].

**FIG. 5.**
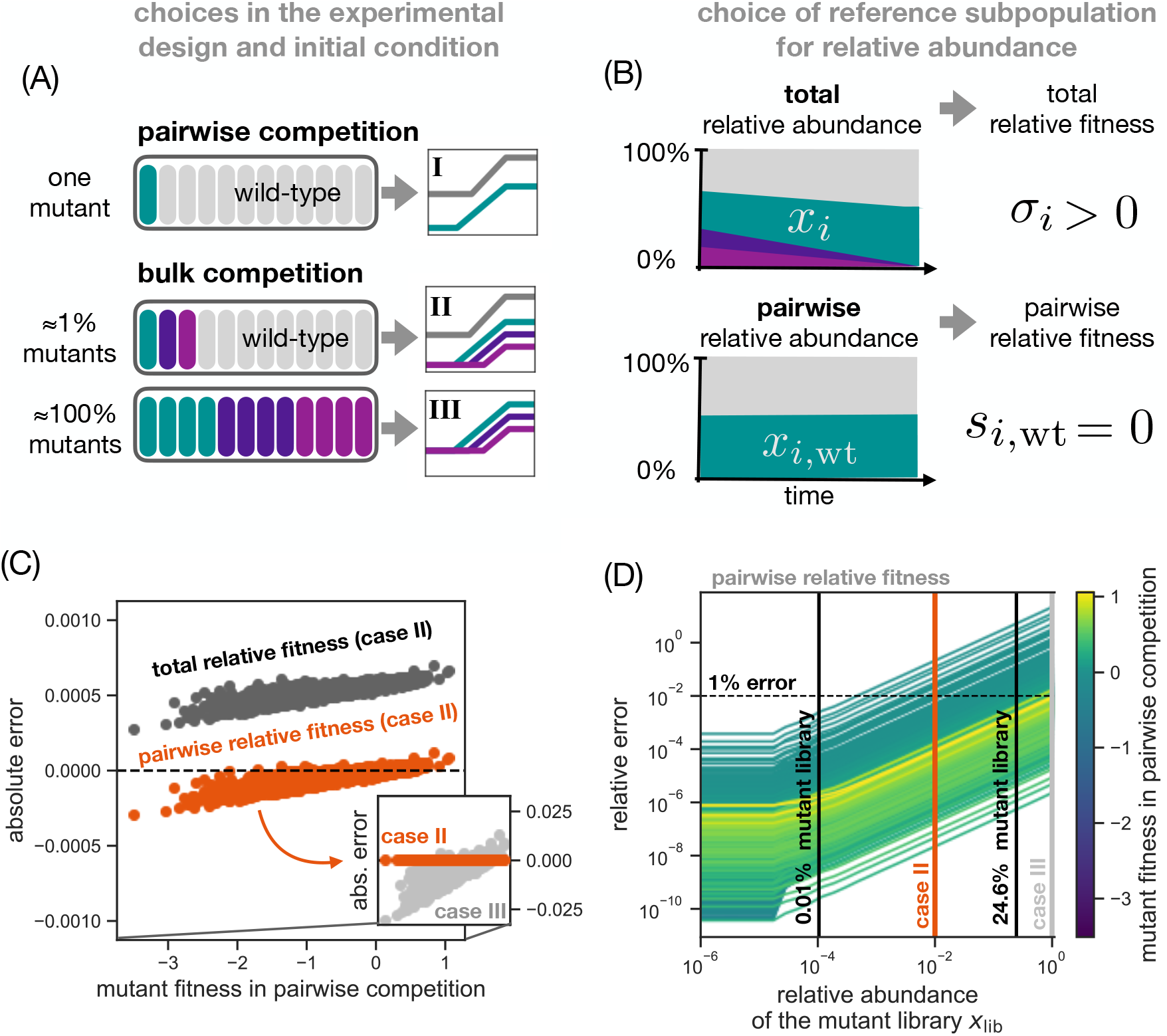
The choice of library abundance and reference group in bulk competition experiments. (A) Overview of a pairwise competition experiment with one mutant (top row) and scenarios for bulk competition experiments with many different mutants (middle and bottom rows). The two bulk competition scenarios differ in their initial fraction of the mutant library (colored ovals) in the inoculum (open box). For each scenario, we show a schematic growth cycle (log absolute abundance) in the inset on the right. (B) Schematic relative abundance trajectories for a mutant compared to two alternative subpopulations. We distinguish between the total relative abundance *x*_*i*_ with respect to the population as a whole (height of green band in the top box) and the pairwise relative abundance *x*_*i*,wt_ with respect to the wild-type (height of green band in the bottom box; Eq. (20)). We indicate the sign of total relative fitness (Eq. (24)) and pairwise relative fitness (Eq. (25)) on the right. (C) The absolute error between bulk and pairwise competition experiments. The total relative fitness (dark grey dots; Eq. (24)) and the pairwise relative fitness (orange dots; Eq. (25)) for mutants in the minimal population dynamics model (Fig. 2A, Methods) parameterized by the yeast knockout data (Fig. S4) in bulk competition growth cycle with low mutant library abundance (panel A, case II; Methods). The absolute error is defined as the bulk fitness statistic minus the relative fitness in pairwise competition (Eq. (S92) in Sec. S15). In the inset, the absolute error for pairwise relative fitness (Eq. (25)) for a bulk competition growth cycle with 99.9% library abundance (light grey dots; case III). This still includes a wild-type reference at small percentage to be able to compute a pairwise relative fitness. The x-axis and the orange dots in the inset are identical to the main plot. (D) The relative error in bulk competition experiments as a function of mutant library abundance in the inoculum. Each line corresponds to a knockout mutant in our data set, and represents the relative error between the pairwise relative fitness in bulk competition and the relative fitness in pairwise competition (Eq. (S102) in Sec. S16). The black vertical lines show the recommended mutant library abundance for our data set based on Eq. (11) (*x*_lib_ ≈ 24.6%) and based on the more conservative Eq. (S118) (*x*_lib_ ≈ 0.02%, Sec. S16). Note these bounds are calculated *a priori* and are not the result of a fitting procedure.

A second important choice is whether to quantify fitness of a mutant relative to the whole population (“total relative fitness”) [18, 19, 80] or relative to another specific genotype, like the wild-type (“pairwise relative fitness”; see Fig. 5B) [40, 48–52]. In the case of a mutant library growing by itself (Case III), it is still possible to estimate a pairwise fitness by using a few known neutral mutants as the reference population [15, 17] and calculating fitness relative to this group (Methods; Secs. and S12).

To test the consequences of these choices, we use the minimal population dynamics model (Fig. 2A) parameterized by yeast single-gene knockouts and compare the estimates of total and pairwise relative fitness in bulk competitions to fitness measured in pairwise competition, which we use as the ground truth (Methods). Figure 5C shows that in bulk competitions, the error in total relative fitness (compared to fitness in pairwise competitions) is systematically offset from the error in pairwise relative fitness. For example, a mutant that grows identically to the wild-type (neutral phenotype) does not change in abundance relative to the wild-type, but it may have a net increase in its total relative abundance due to the poor growth of other mutants (compare top and bottom panel in Fig. 5B). This offset affects the total relative fitness of all mutants in a uniform way (Fig. 5C), such that total and pairwise relative fitness completely agree in the ranking of genotypes (Fig. S21). Note that this offset between total and relative fitness is general and does not depend on our minimal population dynamics model (Sec. S13).

While the pairwise relative fitness is the preferred method for estimating fitness in bulk competitions, the presence of higher-order interactions means it still deviates from the fitness in pairwise competitions. Figure 5C shows the absolute error for pairwise fitness has a positive trend with fitness and increases with the relative abundance of the mutant (Case III, inset in Fig. 5C). For the specific population dynamics in our simulation, we decompose the higher-order interaction into two terms (Secs. and S15): a fitness-independent term and a fitness-dependent term that acts as an amplifier for fitness values (Fig. S22) and drives the positive error trend in Fig. 5C. Intuitively, the population in bulk competition consumes resources more slowly and gives more time for growth rate differences to accrue. The amplification of mutant pairwise fitness in bulk is important because it widens the apparent distribution of fitness effects (leading to, for example, higher estimates for the speed of adaptation) than if fitness was measured in pairwise competition.

In order to control the error from higher-order interactions, we derive an explicit error bound. Figure 5D shows the relative error from higher-order interactions for each mutant across a range of mutant library abundances. Using the specific population dynamics of our model, we derive the following estimate: Assuming that the mutants in the library only have variation in growth rate, the mutant library abundance should satisfy

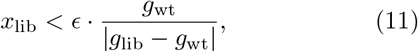

where *ω* is the desired threshold on the relative error (Fig. S23), *g*_wt_ is the wild-type growth rate, and *g*_lib_ is the growth rate of the library as a whole (Sec. S16). In the case of our single-gene knockout library, Eq. (11) predicts a maximum library abundance of 24.6% based on the wild-type and library growth rate (*g*_wt_ = 0.406, *g*_lib_ = 0.389) for a relative error *ω* = 1%. Figure 5D shows that this maximum library abundance keeps the relative error below 1% for high-fitness mutants (bright yellow), because they are dominated by growth rate effects, but fails for mutants close to neutrality because they have a tradeoff between growth rate and lag time, which the estimate in Eq. (11) neglects. It is possible to derive a more precise bound that keeps the relative error below 1% for all mutant genotypes (in this case *x*_lib_ = 0.01%, see vertical line in Fig. 5D), but this requires prior knowledge of the trait covariation in the mutant library (Sec. S16).

### Previous benchmark experiments show deviation of bulk fitness from pairwise competition

Our models predict that higher-order effects amplify pairwise fitness measured in bulk compared to the fitness in a pairwise competition experiment. To test this prediction, we reanalyze two published benchmark experiments of bulk fitness measurements. The first data set by van Opijnen et al. [106] measured pairwise (1:1) and bulk competitions (*x*_lib_ ≈ 99.9%) of 46 single-gene knockouts in *Streptococcus pneumoniae* using logit-encoded fitness per-generation and neutral strains as the reference group. Figure 6A shows that the fitness in bulk deviates from the diagonal, and the absolute error has a positive correlation with the underlying true fitness from pairwise competition (Fig. S24A; Pearson *r* = 0.40,*p* = 0.006), similar to the pattern observed in our simulations (compare Fig. S24A with Fig. 5C case III, and Fig. S24B with Fig. S23C case III).

**FIG. 6.**
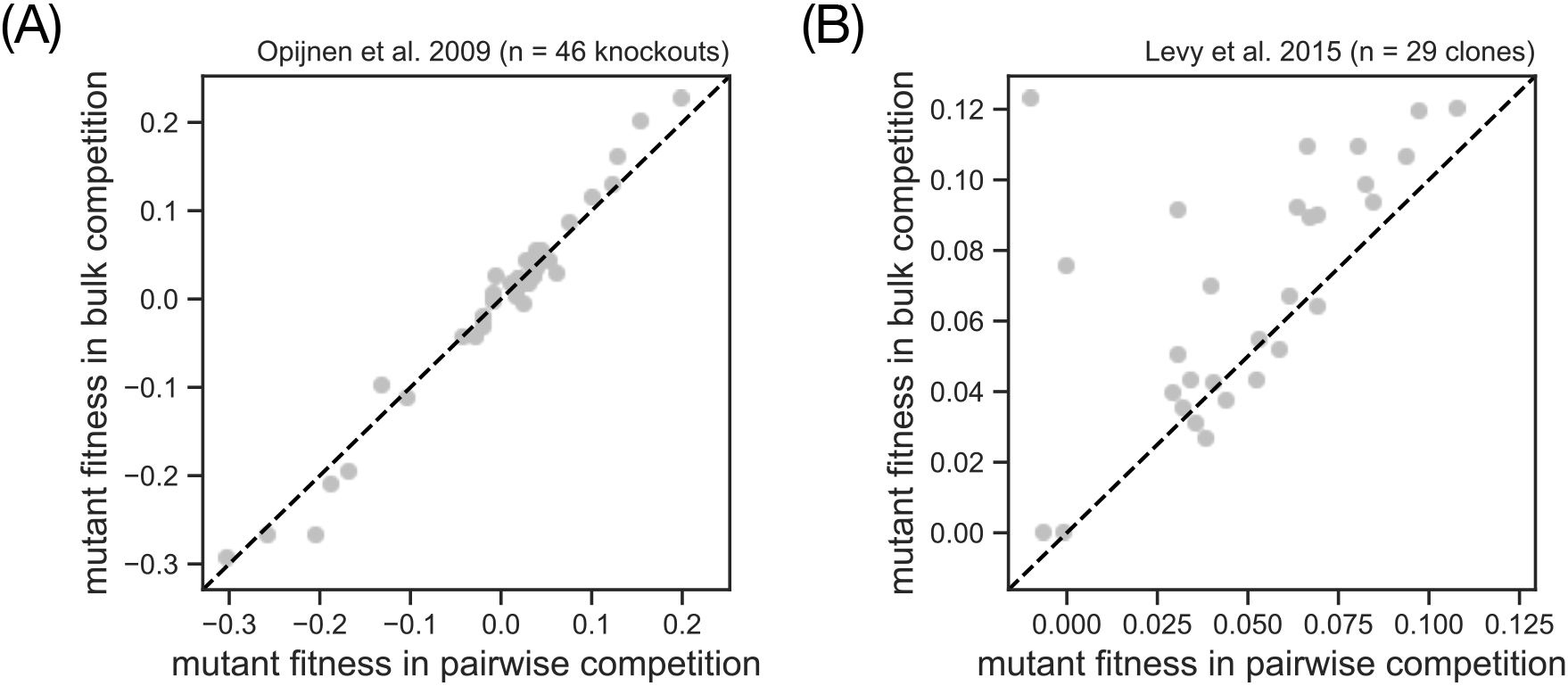
Empirical benchmarks of bulk fitness against pairwise competition experiments. (A) Fitness measurements as reported by Opijnen et al. [106] for knockouts of *S. pneumoniae*. Pairwise fitness was measured in 1:1 competitions of knockouts strains with the wild-type over a single growth cycle using the per-generation statistic 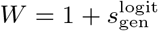 [26]. Bulk fitness was measured from a single growth cycle using 100% initial abundance of the mutant library and a set of neutral strains as the reference to calculate a per-generation fitness, replacing LFC_wt_ with the library LFC in Eq. (S31) (Sec. S5). Since we were unable to obtain the original data, we extracted these fitness data from Fig. 3c in the original publication and subtracted 1 to plot as 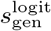. See Fig. S24 for the absolute and relative error of bulk fitness based on these data. (B) Fitness measurements as reported by Levy et al. [13] for evolved genotypes of of *S. cerevisiae* after 80 generations of the evolution experiment. Pairwise fitness was measured in 1:1 competitions of knockouts trains with a fluorescently-labeled wild-type over seven growth cycle using relative fitness per-cycle under the log-encoding 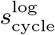, divided by a fixed number generations inferred from the dilution factor. Bulk fitness was measured from relative abundance data throughout the evolution experiment and using a multi-step algorithm, Bayesian inference, and a set of neutral strains as the reference, leading to a pairwise fitness estimate equivalent to 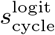 (compare SI Eq. (50) in Levy et al. [13]). We extracted the fitness data from the table presented in SI Sec. 8.1 of the original publication. See Fig. for the absolute and relative error of bulk fitness based on these data.

The second data set by Levy et al. [13] measured pair-wise competitions (1:1) and relative abundances for 33 lab-evolved genotypes of *S. cerevisiae* with beneficial fitness. The bulk fitness is inferred under the log-encoding and per-cycle, with neutral strains as the reference group. Figure 6B shows that the fitness of strains in the bulk evolution environment is typically higher than in pair-wise competition, but the absolute error has no positive correlation (Fig. S25A; Pearson *r* = −0.26,*p* = 0.140) and thus does not support the pattern we see our simulations (compare Fig. S25A with Fig. 5 case III, and Fig. S25B with Fig. S23C case III).

## DISCUSSION

### Significance of fitness quantification choices for microbial ecology and evolution

In this work, we have introduced a conceptual framework to derive common statistics of relative fitness from three essential choices:

1. The choice of the state variable for relative abundance (encoding *m*).
2. The choice of the time scale for the change in relative abundance (Δ*t*).
3. In bulk competition experiments, the choice of sub-population that acts as the reference for relative abundance of a chosen mutant genotype (Fig. 5B).

The combination of these choices leads to a range of fitness statistics, including those commonly used in population genetics [83], experimental evolution [11, 12, 26], and gene knockout screens [17–19, 42, 43, 80, 81, 106].

However, as we compare these statistics in simulated competition experiments, we find that the choice of time scale can lead to different mutant rankings (Figs. 2B and 4A,B), confound apparent epistasis between mutations (Fig. 3B), and alter the apparent long-term trend in fitness (Fig. 3F), while the choice of the reference subpopulation can lead to a systematic shift in inferred fitness values (Fig. 5C). The quantitative differences between relative fitness statistics affect our ability to make evolutionary predictions. Fixation probabilities and other population genetics quantities depend on ratios of parameters defining the rates of selection, mutation, and genetic drift processes [66, 67]. These rates must be measured in the same units. For example, in batch cultures with serial transfer, genetic drift is set by the dilution bottleneck and thus operates per cycle (e.g., 1*/N* per cycle, where *N* is the bottleneck population size). Thus quantifying a mutation’s relative fitness *s* (equivalent to the rate of selection) per generation and comparing to genetic drift per cycle (*Ns*) would incorrectly predict the fixation probability, especially in cases where per-generation and per-cycle fitness rank mutants differently. The choice of statistic may also affect the outcome of a significance test in the context of gene essentiality tests (e.g., using log or logit with the test statistic in Wetmore et al. [18]). Since these measurements often serve as a first screen to narrow down the investigation to the top set of genes [17, 40], a difference in mutant ranking means the investigation might miss relevant genes because of the choice of fitness statistic.

### Best practices for quantifying mutant fitness in high-throughput experiments

Based on these insights, we recommend the following choices for quantifying relative fitness of a mutant:

1. Use the logit encoding of relative abundance, because under the null model of logistic dynamics this linearizes the trajectory of relative abundance. Only use a different encoding if alternate, non-logistic dynamics are known.
2. For batch growth or any experiment with non-steady state growth, use a fixed extrinsic time scale (e.g., a single growth cycle), rather than an intrinsic time scale (e.g., the number of generations), which introduces additional variation between competitions. Only quantify fitness per generation for experiments that are known to have steady-state growth (e.g., in a chemostat).
3. In bulk competition experiments, always use the pairwise fitness relative to the wild-type (by including a barcoded wild-type or by grouping neutral mutants into a virtual wild-type) because this more closely matches the relative fitness in pairwise competition.
4. To control the error from higher-order interactions, we recommend minimizing relative abundance of the mutant library as practically feasible. For a given error tolerance, Eq. (11) gives an estimate for the maximum library abundance.

Our recommendations agree with previous criticisms of relative fitness per-generation [5, 8, 9, 94] and total relative fitness [59, 107] but differ from the standard practice in high-throughput genetic screens [18, 19, 80] and many evolution experiments [33, 91], including the LTEE [11, 26]. The choice of the reference group is a well-recognized issue in high-throughput evolutionary studies, in which one first estimates a total relative fitness and then subtracts a correction based on the fitness of the wild-type [12, 50, 52] or a mean population fitness [13, 41, 42, 51, 107]. In contrast, we recommend to choose the reference group at the level of relative abundance (see Methods for a step-by-step demonstration). While researchers must ultimately choose how to quantify fitness based on the specific goals and conditions of their experiment, we believe that a more consistent adherence to the recommendations above would be valuable to the field as a whole so that fitness data from one study can be quantitatively compared to fitness from another.

### How general is the disagreement between fitness statistics in pairwise competition?

The discrepancy in mutant ranking for relative fitness per-cycle 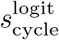 and per-generation 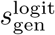 requires positive covariation between the wild-type and mutant log fold-changes (LFCs) across a set of pairwise competition experiments. This condition is not dependent on any particular mechanism or model, although our minimal model of population dynamics (Fig. 2A) demonstrates one way this covariation can occur: when mutants have diminished yields and are present at sufficiently-high abundances to affect wild-type LFCs (Fig. S7).

Our model focused on the effect of resource competition on fitness discrepancies since that interaction is almost always present between microbial genotypes, providing a baseline expectation. Moreover, resource competition is likely to be the only interaction in high-throughput fitness measurements of mutants with only small genetic differences (e.g., point mutations or gene knockouts) relative to a wild-type. We expect that in cases with additional interactions between mutants and the wild-type, there will be further variation in wild-type LFCs across competitions and thus even more discrepancies betwen fitness statistics. For example, mutants that stimulate wild-type growth via the secretion of nutrients should create positive LFC covariation across mutants and thus be ranked differently per-cycle and per-generation. Indeed, our reanalysis of LTEE competition data shows that fitness per-cycle 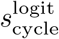 and pergeneration 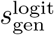 (equivalent to the statistic 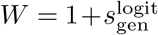 used in the LTEE) disagree on the fitness ranking at almost every time point (Fig. 4A) and in the first 50,000 generations overall (Fig. 4B).

### How general is the deviation of bulk fitness statistics from pairwise competition experiments?

The offset in total relative fitness (between grey and orange points in Fig. 5C), equivalent to the mean library fitness (Eq. (S88) in Sec. S13), is independent from the population dynamics (Fig. 5B, Sec. S13). In contrast, the error between pairwise fitness in bulk competition and in pairwise competition (orange points in Fig. 5C) is specific to the population dynamics studied here, although it points to more general mechanisms. Specifically, we expect pairwise fitness in bulk competition to deviate from pairwise competition as soon as the mutant library is able to influence the chemical environment of the competition, for example, when a large fraction of mutants fails to complete respiration of the main carbon source and instead turns to fermentation [108–110], with the release of pathway intermediates like acetate that indirectly lower the pH in the growth medium [111, 112].

### The choice of fitness statistic in the light of other sources of discrepancy

Besides the conceptual choices of fitness quantification discussed in this article, experimental limitations can create discrepancies between replicate fitness measurements of the same mutant [36, 50, 113]. For example, relative abundance measurements entail sampling uncertainties (when we sample liquid for colony counting or DNA sequencing) [44, 59, 60], copy number variation [18, 44], as well as PCR jackpots and sequencing read errors [13, 104, 105]. Furthermore, there will inevitably be some variation in initial condition between replicates and fluctuations during the fitness assay [36, 105, 113].

## METHODS

### Inferring growth traits for the single-gene knockout collection in yeast

We use a previously published data set [29] for the single-gene knockout collection in *Saccharomyces cerevisiae* [99], where the authors track growth of each genotype for 47 hours in monoculture, using microwell plates with defined growth medium. We download this growth curve data set from the PROPHECY database [114], choosing specifically the data set measured in Synthetic Defined medium. We last accessed the PROPHECY website (http://prophecy.lundberg.gu.se/) on March 30, 2020, but as of November 15, 2025 the website is no longer accessible. We have included a reformatted version of the data with our code repository (https://github.com/justuswfink/24FitnessQuantification). From the raw time series of optical density, the authors subtracted a background correction based on blank wells, applied another correction for nonlinearity at high optical densities, and smoothed the growth curve to remove electrical noise [29].

We start with the curves in the published data set (9951 curves) and apply further trimming and smoothing steps (Sec. S6). From this data we calculate the time series of instantaneous per-capita growth rate *N*^−1^*dN/dt* (where *N* is the optical density) and identify time windows where the rate is approximately constant, which we interpret as distinct growth phases (Sec. S6). We only include curves that have a single phase of constant exponential growth followed by a stationary phase of approximately zero growth (9424 curves).

We quantify the biomass yield, maximum growth rate, and the lag time from each remaining curve as follows. First, we estimate the initial abundance *N*_initial_ (average optical density over first three time points) and the final abundance *N*_final_ (average optical density over stationary phase) from the growth curve. Then we calculate the biomass yield as *Y* = (*N*_final_ − *N*_initial_)*/R*(0), where *R*(0) = 111 mM (20 g/L) is the initial concentration of glucose (assuming glucose is the single limiting resource). To estimate the maximum growth rate, we average the instantaneous growth rate over the exponential phase. Finally, we estimate the lag time from the intersection of the log initial abundance (log *N*_0_) with the slope of the maximum growth rate during the exponential growth phase [84]. We excluded curves with negative initial OD, curves with negative inferred lag times, and curves with a low quality of fit between the measured time series and a simulated curve based on the inferred trait values (*R*^2^ *<* 0.95; see below for the model). Our final database includes trait estimates for 9195 curves, which represents 92.4% of the original data set [29].

Each single-gene deletion strain was measured in two technical replicate growth curves, using a second plate with identical layout in the same plate reader. Some genotypes have only one estimate of the growth traits because the other replicate did not pass our filters (273 genotypes), but for most genotypes we retain two replicate estimates in our final data set (4163 genotypes); a few genotypes even have three (2 genotypes) or four replicate estimates (54 genotypes) because these genotypes were included multiple times by the original authors [29]. From these replicates we finally calculate the average yield, growth rate, and lag time for each genotype. The data set also contained many replicate growth curves of the wild-type strain, 374 of which passed our filters. Since wild-type traits inferred from these replicates had large variation (potentially due to measuring the wild-type across many different plates and days), we define the wild-type trait as the median, not the mean, of these replicates. Although the wild-type measurements included in this data set have large trait variation (orange dots in Fig. S4B,C), the fact that the variation across knockouts is significantly greater than for the wild-type replicates in terms of growth rate (Levene’s test *p* = 1.3 × 10^−9^) and lag time (Levene’s test *p* = 0.025), and that the replicate measurements of gene knockouts are correlated (Fig. S5), suggests that these trait measurements do capture true genetic variation. In contrast, the variation in biomass yield across knockouts is not significantly different from the variation across wild-type replicates (Levene’s test *p* = 0.44), suggesting that the variation in biomass yield is driven by non-genetic factors like the abundance or batch of resources used for each growth measurement.

### Simulating population dynamics in competition experiments

Throughout this work, we model the competition between a set of genotypes during a single batch culture growth cycle using the following model of population dynamics [97, 98]:

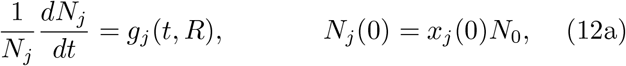

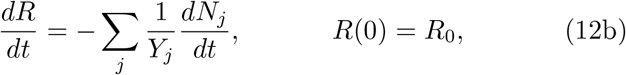

where for each genotype *j, N*_*j*_ is the absolute abundance, *x*_*j*_(0) the initial relative abundance, *g*_*j*_(*t, R*) the growth rate, and *Y*_*j*_ the biomass yield, where the genotypes include a wild-type as well as one or more mutants. The initial absolute abundance of all genotypes together is *N*_0_, and the concentration of the single limiting resource is *R* with initial value *R*_0_. Note that Eq. (12b) assumes that cells consume resources only for biomass growth and not for maintenance of existing biomass.

To capture the sigmoidal shape of typical growth curves with a lag phase, exponential growth phase, and saturation phase (e.g., Fig. S4A), we model the dependence of growth rate on time and resource concentration as

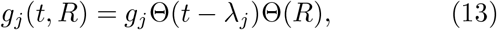

where Θ is the step function:

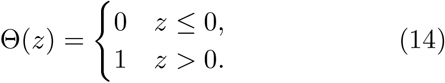

In this model, a genotype has an initial lag phase of time *ε*_*j*_ where no growth occurs, followed by a phase of constant exponential growth at rate *g*_*j*_, and ending when the resource concentration *R* reaches zero. The time to resource depletion, which we also call the saturation time, is defined by the implicit equation *R*(*t*_sat_) = 0. The simple form of the growth response (Eq. (13)) means that the absolute abundance of genotype *j* at saturation is

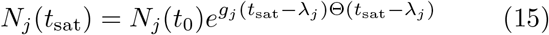

and its log fold-change is

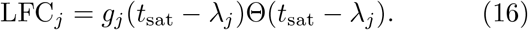

To calculate relative fitness per-cycle (Eq. (9) and per-generation (Eq. (10)), we numerically determine the saturation time *t*_sat_ by integrating the biomass and resource dynamics (Eq. (12)) as described in previous work [97, 98]. We note that it also possible to derive an approximate expression for the saturation time *t*_sat_ which we use for our analytical calculations (Sec. S7).

### Testing epistasis between mutations in population dynamics model

Consider a wild-type strain with growth rate *g*_wt_ = 1, lag time *ε*_wt_ = 2, and biomass yield *Y*_wt_ = 1. We define six basic mutation that represent the six directions of the trait space in our model: a mutation that increases growth rate (*g*_mut_ = 1.2; symbol +*g*), a mutation that decreases growth rate (*g*_mut_ = 0.8; symbol − *g*), a mutation that increases lag time (*ε*_mut_ = 3; symbol +*ε*), a mutation that decreases lag time (*ε*_mut_ = 1; symbol − *ε*), a mutation that increases yield (*Y*_mut_ = 4; symbol +*Y*), and a mutation that decreases yield (*Y*_mut_ = 0.25; symbol −*Y*). We combine these mutations into double mutants as follows: mutations are additive at the level of traits (e.g., the double mutant (+*g*, −*g*) is neutral) and mutations in different traits are independent. For a given pair of mutations, epistasis is defined as [100]

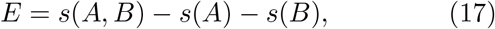

where *s*(*A*) is the relative fitness of the single mutant with mutation *A, s*(*B*) is the relative fitness of the single mutant with mutation *B*, and *s*(*A, B*) is the relative fitness of the double mutant with both *A* and *B*. We estimate the fitness of each mutant by simulating one growth cycle of competition with the wild-type (at initially equal abundance) using an initial biomass of *N*_0_ = 0.01 and initial resource concentration *R*_0_ = 0.5. These conditions are chosen such that mutations that increase biomass yield reach a maximum density of up to *N* (*t*_sat_) = 2, on the same scale as the optical density measured for the empirical knockouts of *S. cerevisiaea* (Fig. S4C).

### Quantifying total and pairwise relative fitness in bulk competition experiments

We define total and pairwise relative fitness,two alternative measures of fitness in bulk competition experiments, as follows. Consider a population with multiple genotypes such that each genotype *i* has absolute abundance *N*_*i*_(*t*) at time *t*. The *total relative abundance* of the genotype is

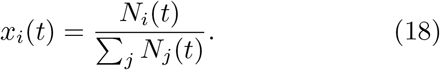

The “total” here refers to the fact that this is the abundance of genotype *i* relative to all other genotypes in the population. Similarly, we can think of the definition of relative fitness discussed in the main text (Eq. (4)) as the *total relative fitness* of this genotype under an encoding *m*:

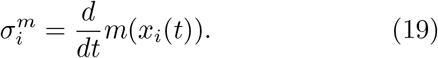

However, sometimes we want to track the dynamics of a genotype relative to another specific genotype; for example, to follow the fate of a mutant against the wild-type in a bulk competition experiment. For a pair of genotypes *i* ≠ *j*, we thus define the *pairwise relative abundance*

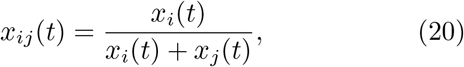

which is the relative abundance of genotype *i* in a sub-population of only genotypes *i* and *j*. To predict the change in the pairwise relative abundance, we can use the slope

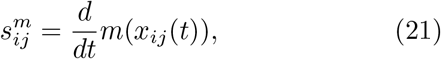

which defines the *pairwise relative fitness* of genotype *I* with respect to genotype *j* (see Sec. for a generalize definition that covers the case *i* = *j*). In the special case of a population with only two strains, the pair-wise relative fitness and total relative fitness are identical 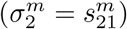 but they may differ with more than two genotypes (Sec. S13).

It is also possible to define the total and pairwise relative fitness statistics as finite differences over a time interval, rather than as instantaneous derivatives. For example, we can define them over a growth cycle starting at *t* = 0 and ending at *t* = *t*_sat_:

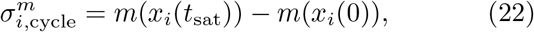

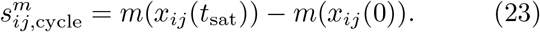

For these fitness statistics in the bulk competitions, we use the logit encoding

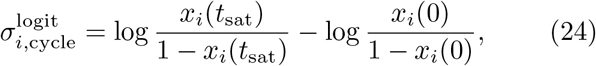

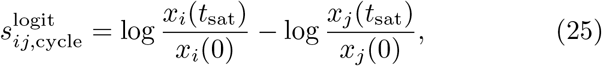

where we have rewritten 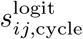 in the form that it is presented in bulk competition experiments [15, 51, 52]. The logit encoding has mathematical advantages for coarse-graining the relative fitness of genotype groups (Sec. S11) but using the log encoding is another common choice in the literature [17, 41, 50]. However, the relative abundances of the individual mutants in our simulated bulk competition experiments are low enough that these two encodings are approximately equivalent (logit *x* ≈ log *x* for *x* ≪ 1).

Unlike the comparison between per-cycle and per-generation relative fitness where we focused on rank differences (Fig. 2B), here we can evaluate absolute differences in fitness estimates because the total and pairwise fitness in bulk fitness are measured in the same units as the relative fitness in pairwise competition. For all mutant genotypes, we calculate the absolute error between these bulk fitness estimates (Eqs. (24) and (25)) and the relative fitness in pairwise competitions (Eq. (9)), which we take as the ground truth (Fig. 5C). Note that the pairwise competition could depend on the initial relative abundance of the mutant; we have chosen a very low relative abundance (10^−6^) that mimics a mutant arising *de novo* and where this dependence is very weak.

Clearly, minimizing the abundance of the mutant library minimizes the strength of higher-order interactions (also reducing ranking disagreement, Fig. S21B,C). As low mutant abundances create practical problems for experiments, it is valuable to identify a maximum abundance for the mutant library that keeps the error from bulk fitness estimates below a desired threshold (Eq. (11)). It is convenient to express this threshold in relative rather than absolute terms (compare Fig. S23 to Fig. 5C).

### A guide to calculate pairwise relative fitness under the logit encoding from bulk competition data

Here *r*_*i*_(*t*) represents the best estimate of read numbers for lineage *i* after correcting for experimental artifacts, like jackpot events in the sequencing (e.g., see discussion in Levy et al. [13]).

1. Calculate the total relative abundance *x*_*i*_ of each lineage *i* by dividing with the total number of reads:

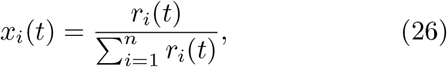

where the sum is over all *n* lineages in the experiment.
2. Define a group of lineages *i* = 1, …, *k* as the reference (e.g., a group of neutral mutants) and sum their total relative abundance

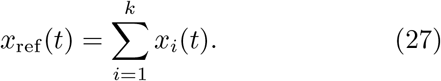
3. Calculate the pairwise relative abundance *x*_*i*,ref_ of each lineage by

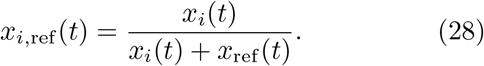
4. Calculate the pairwise fitness per-cycle for each strain *i* under the logit encoding

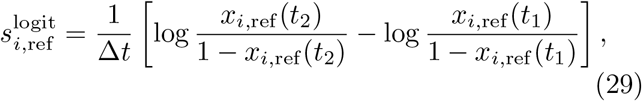

where Δ*t* is the number of growth cycles between time points *t*_1_ and *t*_2_.

## ACKNOWLEDGMENTS

JWF wishes to thank Luis-Miguel Chevin, Olivier Tenaillon, Henrique Teotónio, and the participants of the 2021 ENS autumn course on Experimental Evolution for valuable discussions early on. The authors are grateful to Jeremy Schreier, Benjamin Raach, and Gatwa Tshinsele-Van Bellingen for a critical reading of the manuscript. JWF and MM were supported by an Ambizione grant from the Swiss National Science Foundation (PZ00P3 180147).

## Supplementary Information

## S1. DIFFERENT TYPES OF FITNESS UNDER EXAMPLE MODELS OF POPULATION DYNAMICS

In this section we explicitly calculate relative fitness of a mutant under a few example models of population dynamics, using the different encodings of relative abundance as described in the main text (main text Fig. 1A). Consider a competition coculture between a wild-type genotype with absolute abundance *N*_wt_(*t*) and a mutant genotype with absolute abundance *N*_mut_(*t*). We can describe their dynamics according to the ordinary differential equations (ODEs)

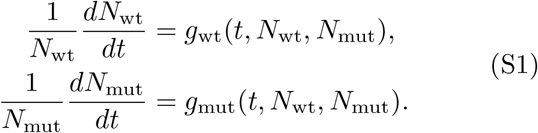

Note that the per-capita growth rates *g*_wt_ and *g*_mut_ of each genotype can depend on both genotypes to reflect competition or other interactions. The relative abundance of the mutant genotype at time *t* is

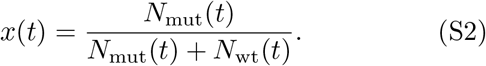

The dynamics of the mutant relative abundance are therefore described by

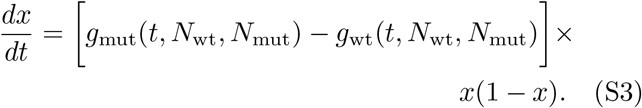

As defined in the main text (Eq. (4)), the relative fitness of a mutant is *s*^*m*^ = *dm/dt* for an encoding *m*(*x*) of the relative abundance *x*. Under the trivial linear encoding (*m*(*x*) = *x*), the relative fitness is therefore just the right-hand side of the relative abundance ODE (Eq. (S3)):

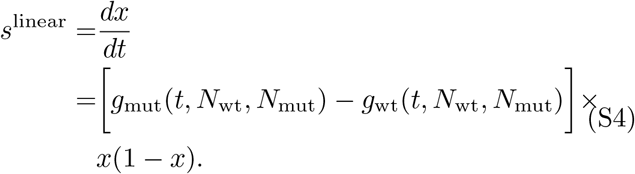

For the log encoding *m*(*x*) = log *x*, we use the identity *d* log *x/dt* = *x*^−1^*dx/dt* to obtain the log-encoded relative fitness:

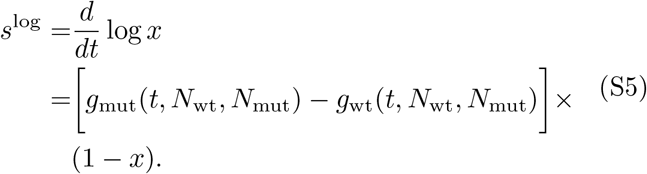

Finally, for the logit encoding *m*(*x*) = logit *x* = log(*x/*(1 − *x*)), we use the identity *d* logit *x/dt* = *x*^−1^ (1 − *x*)^−1^*dx/dt* to obtain the logit-encoded relative fitness:

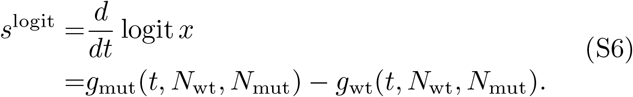

By comparing relative fitness values under the linear encoding (Eq. (S4)) and under the logit encoding (Eq. (S6)), we see how the logit encoding has removed the explicit dependence on the mutant relative abundance (factors of *x* and 1 − *x*), although there can be implicit dependence on the mutant relative abundance within the per-capita growth rates of each strain (*g*_wt_ and *g*_mut_) due to density-dependent growth rates.

If the per-capita growth rates *g*_wt_ and *g*_mut_ (Eq. (S1)) are constants, then these constant growth rates also act as fitness potentials since they each depend only on a single genotype but their difference determines relative fitness (under the logit encoding, Eq. (S6)) between the genotypes. The growth rate is not a fitness potential under more complex dynamics, however. For example, consider a competition model with explicit density dependence:

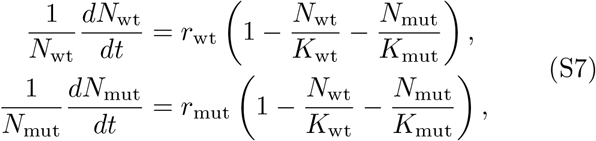

where the growth rates decrease as the genotype abundances reach their carrying capacities *K*_wt_ and *K*_mut_, and the maximum growth rates at low abundances are *r*_wt_ and *r*_mut_. In this case, the relative fitness under the logit encoding is (from Eq. (S6))

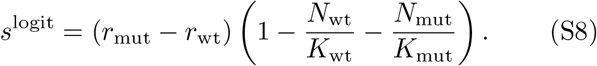

In this case, there is no fitness potential because it is not possible to separate Eq. (S8) into a difference between terms that only depend on each genotype separately.

## S2. DEFINITION OF ABSOLUTE FITNESS FOR A GENOTYPE

Here we give an explicit definition of a genotype’s absolute fitness, analogous to the definition of relative fitness in the main text (Eqs. (1)–(4)). Conceptually, absolute fitness is any number that is sufficient to predict a genotype’s absolute abundance *N* over a short time window. Let an encoding *m*(*N*) be any smooth, strictly-increasing function of the absolute abundance *N*. We can then predict the absolute abundance over a time window Δ*t* using a linear expansion of the encoded abundance (analogous to main text Eq. (3) for relative fitness):

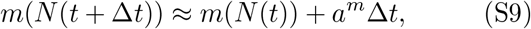

where

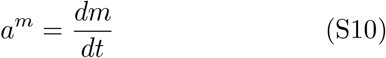

is defined as the absolute fitness of the genotype under the encoding *m* (analogous to main text Eq. (4) for relative fitness; see also main text Fig. 1B). For example, the absolute fitness of a genotype under the log encoding *m*(*N*) = log *N* is the per-capita growth rate:

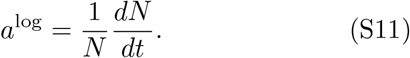

In general, the ideal encoding of absolute abundance is the inverse function of the absolute abundance trajectory *N* (*t*) (up to a shift and rescaling), so that the first-order expansion in Eq. (S9) is exact and the absolute fitness *a*^*m*^ is sufficient to determine changes in absolute abundance up to any future time. The log encoding *m*(*N*) = log *N* is therefore ideal when absolute abundance grows or decays exponentially at a constant rate, while the logit encoding *m*(*N*) = logit *N* is ideal for a population that grows with logistic density dependence (Eq. (S7) in case of a single genotype).

Absolute fitness and relative fitness are related, since the relative abundance of a genotype is determined by normalizing its absolute abundance by the absolute abundance of all genotypes in the population. Specifically, the relative fitness of a genotype is determined by the absolute fitnesses for all genotypes in the population. For example, in the case of two genotypes, the relative fitness under the logit encoding (Eq. (S6)) is the di!erence between the genotypes’ absolute fitness under the log encoding (Eq. (S11)). In the case of constant per-capita growth rates, these log-encoded absolute fitnesses also act as fitness potentials.

## S3. THE ROLE OF LOGISTIC POPULATION DYNAMICS IN LOGIT-ENCODED RELATIVE FITNESS

Here we show how the logit encoding of relative abundance is related to the logistic model of population dynamics. For a relative abundance *x*, logistic dynamics are

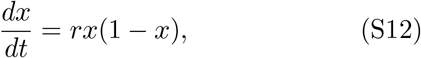

where *r* is the exponential rate at which relative abundance increases from low values. This form emerges from the general dynamics of relative abundance (Eq. (S3)) when the difference in per-capita growth rates *g*_mut_ and *g*_wt_ is constant. Once can also interpret the logistic model as a lowest-order approximation for more complex dynamics. That is, consider a general equation for relative abundance:

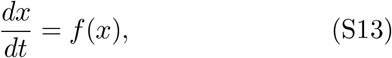

for an arbitrary function *f*(*x*). Since this function must obey the boundary conditions *f*(0) = 0 and *f*(1) = 0 (the relative abundance must stop changing when it either goes extinct or fixes), a polynomial expansion of *f*(*x*) must have roots at these values:

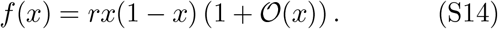

Thus the logistic model in Eq. (S12) is a lowest-order approximation even when the true dynamics are more complex.

The logistic di!erential equation in Eq. (S12) has the solution

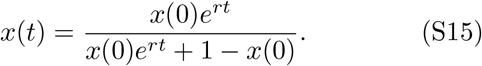

The logit encoding of the logistic relative abundance has linear dependence on time:

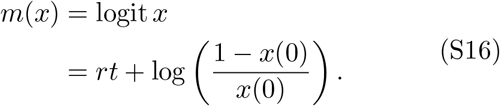

This is another way to see why the relative fitness under the logit encoding (the time derivative of Eq. (S16)) is constant under logistic dynamics. Mathematically, this occurs because the logit function is the inverse of the logistic dynamics (Eq. (S15)), up to a shift and rescaling. Thus if the relative abundance dynamics are different from logistic (Eq. (S12)), the logit encoding no longer exactly linearizes the trajectory of relative abundance and thus is no longer the optimal encoding for relative fitness (see Fig. S2 and the mathematical notes by Mallet [1] for more examples).

More generally, the ideal encoding for a given relative abundance dynamics *x*(*t*) is any linear function of the inverse *m*(*x*(*t*)) = *at*(*x*) + *b* of those dynamics. Note that the ambiguous scale *a* includes log and logit encodings with a different base of the logarithm (discussed by Chevin [2]). The inversion of *x*(*t*) is equivalent to removing frequency (relative abundance) dependence from the relative fitness statistic *s*^*m*^ [3].

## S4. RELATIVE FITNESS PREDICTIONS IN DISCRETE TIME: ADDITIVE VS. MULTIPLICATIVE FORM

In our framework, relative fitness is a statistic that predicts relative abundance in an additive equation (main text Eq. (3)) but sometimes the dynamics of individual genotypes are modeled using a multiplicative form of fitness. In this section, we show how these multiplicative fitness statistics are related to the additive fitness statistics, in particular for the logit encoding.

We consider a population of a wild-type and a mutant genotype, where we track the mutant’s relative abundance over multiple rounds of competition (e.g., growth cycles)

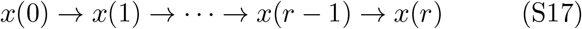

and the variable *x* changes each round according to some underlying population dynamics.

In a modeling approach typical to many studies in population genetics [4], these population dynamics are captured in the genotype-specific growth factors

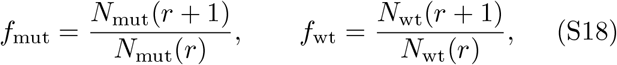

which determine the update equation for the mutant relative abundance

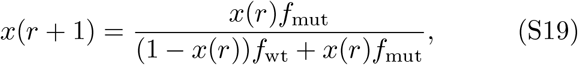

which is the discrete-time analogue to a differential equation (Eq. (S1)). We divide Eq. (S19) by 1 − *x*(*r* + 1) to obtain the form

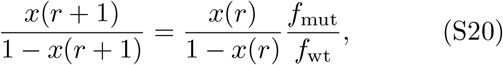

which allows us to recognize the ratio of growth factors *f*_mut_*/f*_wt_ as a relative fitness statistic, since it is sufficient to predict the relative abundance of the mutant genotype (under the encoding *m*(*x*) = *x/*(1 − *x*)). But the statistic *f*_mut_*/f*_wt_ acts as a multiplying factor in Eq. (S20), whereas the general form of relative fitness *s*^*m*^ (main text Eq. (3)) acts as an additive factor. What is the relationship between the multiplicative and the additive form of discrete-time relative fitness statistics?

To describe discrete rounds of population dynamics in our framework, we treat the relative abundance *x*(*r*) as samples from a continuous time series, separated by a time-gap Δ*t* = *t*(*r* + 1) − *t*(*r*). For a chosen encoding *m*, The relative abundance of the mutant genotype in the future round is predicted by

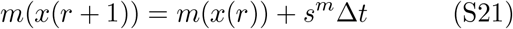

since this is how we defined relative fitness *s*^*m*^ in main text Eq. (3). This relative fitness acts as additive factor, but we apply the exponential function on both sides of Eq. (S21) to obtain the updated equation

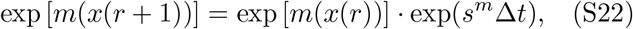

where the additive fitness *s*^logit^ together with the time scale Δ*t* acts as a multiplicative factor. Specifically for the logit encoding *m*(*x*) = logit *x* we have

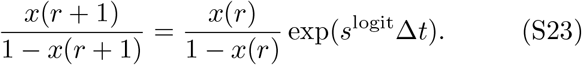

We compare Eq. (S23) to Eq. (S20) and solve for the relative fitness of the mutant genotype

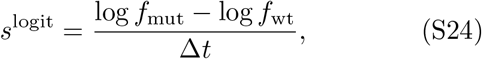

as a function of the growth factors *f*_mut_, *f*_wt_. Using the definition of the growth factors (Eq. (S18)), we see that Eq. (S24) is simply the discrete-time relative fitness for the logit encoding in terms of the mutant and wild-type LFC (main text Eq. (8)). As a general point, we note that the growth-factor *f*_mut_ qualifies as an absolute fitness (since it is sufficient to predict the absolute abundance *N* (*r* +1)), but does not constitute a relative fitness statistic (since we also need to know *f*_wt_, see Eq. (S20) or Eq. (S24)).

More generally, for a given encoding function *m*(*x*) we define the mutant’s multiplicative relative fitness over a discrete growth cycle as

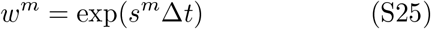

where *s*^*m*^ is the additive relative fitness for this growth-cycle under the chosen encoding (main text Eq. (4)) and Δ*t* is the duration of the growth-cycle in time units that match *s*^*m*^. Equation (S25) shows that the additive fitness *s*^*m*^ has time units, but the multiplicative fitness does not and is formally a dimensionless quantity. These dimensionless units are preserved in the approximate formula

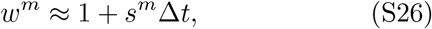

which is the first-order expansion of Eq. (S25) in the limit of weak selection (|*s*^*m*^Δ*t*| ≪ 1). For example, if we choose to measure relative fitness with the logit encoding on the time scale per-cycle (Δ*t* = 1 cycle), we get

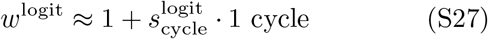

which is the same multiplicative fitness as if we choose to measure the relative fitness on the time-scale per-generation (Δ*t* = LFC^wt^):

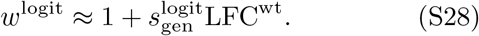

In both cases, the time-units for the discrete-time additive fitness cancel in the product term.

So how would a mutant ranking in the multiplicative fitness *w*^logit^ rank a set of mutant genotypes compared to the fitness statistics 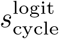 or 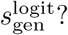 Equation (S27) shows that the multiplicative fitness statistic *w*^logit^ agrees with the relative fitness per-cycle *s*^logit^, but can differ from the ranking in the relative fitness per-generation 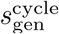 as both terms of the product in Eq. (S28) depend on the mutant.

Finally, we address the question how the relative fitness 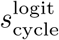 agrees in the ranking with the multiplicative fitness *w*^logit^, but disagrees with the fitness statistic defined in the Long-Term Evolution Experiment [5] as

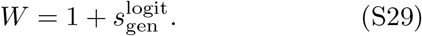

By comparing Eq. (S29) to the multiplicative fitness in Eq. (S26), we see that the LTEE fitness statistic *W* does not derive from the general form of the multiplicative fitness (Eq. (S25)) as all statistics derived from this approximation have a multiplying factor Δ*t* that cancels the units of time in *s*^*m*^. The fact that the LTEE fitness statistic *W* (Eq. (S29)) misses the term LFC_wt_ compared to the logit-based multiplicative fitness *w*^logit^ (Eq. (S28)) means that the objects have different units of time and different rankings.

## S5. DERIVATION OF MISMATCH CONDITIONS FOR RELATIVE FITNESS PER-CYCLE AND PER-GENERATION

In this section, we derive the conditions for a ranking mismatch between the relative fitness per-cycle and per-generation (for the logit-encoded relative abundance) across a set of competition experiments. Consider a batch culture where a competing wild-type and mutant genotype have log fold-changes LFC_wt_ and LFC_mut_ over a single growth cycle. The LFCs are convenient variables to describe these dynamics since we can express the mutant’s relative fitness per-cycle as (main text Eq. (9))

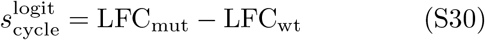

and the mutant’s relative fitness per-generation as (main text Eq. (10))

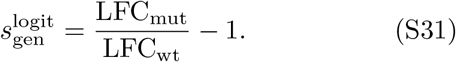

Here we assume both LFCs are nonzero (so that measuring fitness per-generation is meaningful; see discussion in main text).

To determine how these two fitness statistics lead to different rankings, we consider two competition experiments *A* and *B*, which may represent two different mutants competing against the same wild-type or the same mutant tested in two different environments. A mismatch in ranking occurs when the fitness per-cycle in competition is higher in *B*, while the fitness per-generation is higher in *A*:

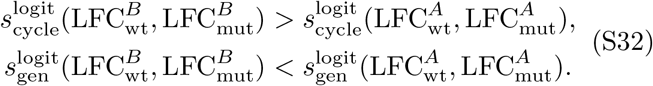

We insert the expressions for relative fitness per-cycle (Eq. (S30)) and per-generation (Eq. (S31)) into Eq. (S32) to rewrite the condition for ranking mismatch in terms of the LFCs:

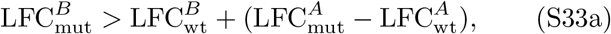

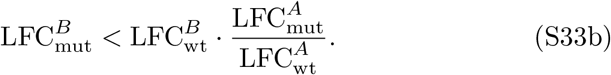

For a given competition *A*, Eq. (S33) defines an area in the space of LFCs where competition *B* can lie such that competitions’ fitness is ranked differently per-cycle versus per-generation (main text Fig. 1D shows an example as the red-shaded area). Biologically, these constraints describe a situation where the mutant and wild-type LFCs are both higher or both lower in competition *B* than in competition *A* (i.e., so that they grey point is up and to the right of the red point in Fig. 1D, or down and to the left), but the LFC differences must be sufficiently balanced between the mutant and wild-type (i.e., so that the point lies within the red area in Fig. 1D).

Typically, however, the LFCs of the wild-type and the mutant are not independent, since these LFCs are jointly constrained by the fact both strains compete for the same finite resources. For example, assume that a single limiting resource with concentration *R* is consumed in proportion to the growth of each genotype’s biomass according to

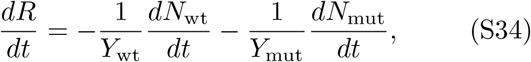

where *Y*_wt_ and *Y*_mut_ are the wild-type and mutant biomass yields (stoichiometry of biomass to resource). We can integrate Eq. (S34) to obtain

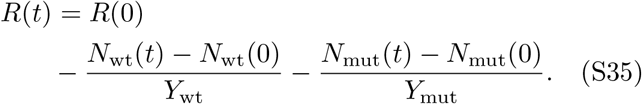

The growth cycle stops when no resource remains (*R*(*t*) = 0), such that

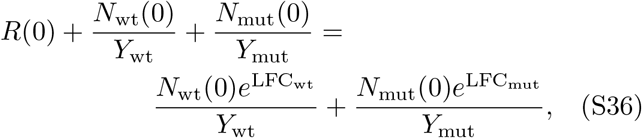

where we have expressed the genotype abundances at the end of the growth cycle in terms of their LFCs. Equation (S36) thus entails a constraint between the wild-type and mutant LFCs. For a set of a mutant competitions with the same initial resource concentration *R*(0), initial abundances *N*_wt_(0) and *N*_mut_(0), and yields *Y*_wt_ and *Y*_mut_, the mutant and wild-type LFCs are constrained by Eq. (S36) to fall along a one-dimensional curve (black line in Fig. S3A). Geometrically, we see that this constraint on LFCs is incompatible with the requirements for a ranking mismatch between fitness per-cycle and per-generation for a pair of mutant competitions (compare black line and red shaded areas in Fig. S3A). However, if some mutant competitions deviate from this constraint, for example by having different yields *Y*_mut_ or initial conditions *N*_wt_(0) and *N*_mut_(0), then ranking mismatches may be possible (Fig. S3B).

## S6. ANALYSIS OF GROWTH CURVES TO IDENTIFY GROWTH PHASES

In this section, we describe in more detail how we identify growth phases from the original data set of growth curves and use this to choose a subset of curves that matches the simplified growth dynamics of our population dynamics model (Fig. S4A). As mentioned in the main text (Methods), we downloaded this original data from the PROPHECY database (http://prophecy.lundberg.gu.se/), downloading specifically the data set for growth in Synthetic Defined medium as first analysed and reported by Warringer et al. [6]. We last accessed the PROPHECY website (http://prophecy.lundberg.gu.se/) on March 30, 2020, but as of November 15, 2025 the website is no longer accessible. We have included a reformatted version of the data with our code repository (https://github.com/justuswfink/24FitnessQuantification). The original growth curve data is already corrected for background and instrument non-linearities [6] (summarized in Methods), but we decided to apply additional corrections as follows: To begin with, we concatenate the original 51 data files (for different plate reader runs) into a single, consecutive dataframe and manually handle a duplication in one of the files (Experiment NO. 18). This file has no measurements for the first time point (*t* = 0) due to technical error in the original data export [6] and for curves from this experimental run, we set the initial time point to <monospace>NAN</monospace> value in Python, meaning that these points will be ignored for any subsequent calculations of averages. More generally, we decided to trim the first four time points of all growth curves (equivalent to 1h20min from 47h total) and remove OD measurement below a noise threshold (OD = 0.001) as this improves the quality of the fit later on.

After pre-processing, we estimate a smooth time series for the instantaneous growth rate in each growth curve, using a previously published script <monospace>gaussianprocess.py</monospace> by Swain et al. [7] that implements the Gaussian Process approach to smoothing (downloaded from https://swainlab.bio.ed.ac.uk/software.html). We apply this script to the logarithmic absolute abundance log OD, and reconstruct a smoothed trajectory *f* (*t*) log OD(*t*) as well as the first derivative *df /dt* and the second derivative *d*^2^*f/dt*^2^ [7]. Effectively, the script estimates three hyperparameters that capture the shape of each curve and we find that the estimation works best if we constrain the parameter ranges as outline in Table S1.

**TABLE S1.**
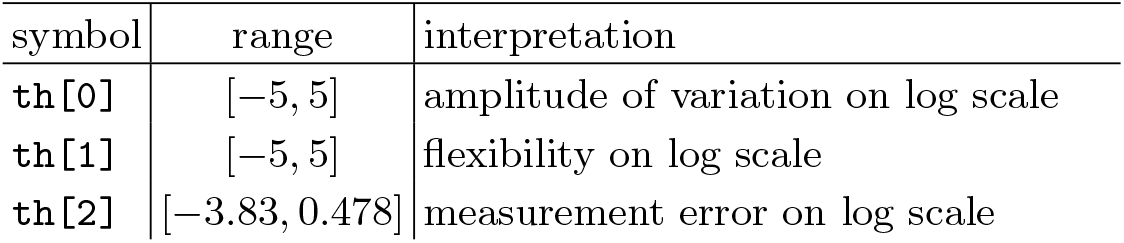
Parameter settings for Gaussian Process optimisation.

| symbol | range | interpretation |
| --- | --- | --- |
| th[0] | $[-5, 5]$ | amplitude of variation on log scale |
| th[1] | $[-5, 5]$ | flexibility on log scale |
| th[2] | $[-3.83, 0.478]$ | measurement error on log scale |

With a smoothed time series at hand, we now identify “plateaus” of constant growth rate using the functions available in Scipy [8]. For each growth curve, we start by identifying so-called “plateau seeds,” which are small intervals where the second derivative is below a chosen threshold *d*^2^*f/dt*^2^ *<* 5 10^−6^. For each plateau seed *k* in the growth curve, we calculate the average growth rate *ĝ*_*k*_ in the time-window of the plateau. Due to some noise in the second derivative, we find many plateau seeds that are adjacent and need to be merged. To do so, we iterate over the plateau seeds in the growth curve and merge the current candidate *k* with the previous plateau *k* − 1, except one of the following conditions is true:

- Both plateaus seeds have a duration that is too long (equal or greater than 100 minutes).
- The transition time between the plateau seeds is too large (equal or greater than 200 minutes).
- The first plateau has a significantly different growth rate that the second one, sucht that the following equation is satisfied

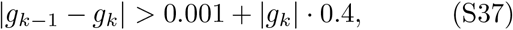

where *g*_*k*_, g_k−1_ are the average growth rate in each plateau (per minute).

Empirically, we find that the duration of these merged plateaus is shorter than what one would expect from visually inspecting the growth curve. Therefore we extend the remaining plateaus in each growth curve as follows: For each plateau, we estimate the lower and upper growth rate *df /dt* in the time window and take this as a growth rate corridor. We extend the plateau to the left, until *df /dt* leaves that growth rate corridor, and similarly extend to the right. By definition, the resulting plateau is equal or larger to the original time-window and we recal the average growth rate over the time window.

From this analysis, we obtain a list of growth phases for each curve that allows us to choose a subset of curves that match our model of population dynamics (Methods). We only choose curves that have two plateaus, where the first plateau has significant growth (exponential phase), and the second plateau has no growth (stationary phase). Here we define significant growth as the average growth rate in the plateau time window is larger or equal to 0.0011 per minute. This forms the set of growth curves that we use to estimate growth traits (9424 curves).

## S7. THE SATURATION TIME IN OUR MODEL OF POPULATION DYNAMICS

In this section, we restate an explicit expression for the saturation time *t*_sat_ in pairwise competition that was derived in earlier work and allows us to see how mutants can influence resource depletion and the wild-type LFC (main text Eq. (16)). Using the same model of population dynamics (Methods, main text Eq (12)), previous work [9, 10] derived an approximate formula for *t*_sat_ that shows how it depends on the underlying parameters:

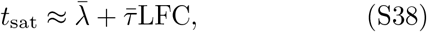

where

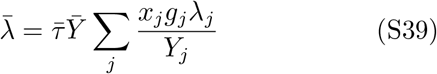

is the effective lag time of the population,

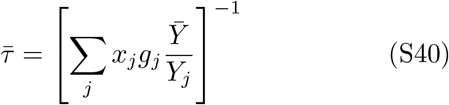

is the effective *e*-fold growth time (reciprocal exponential growth rate) of the population,

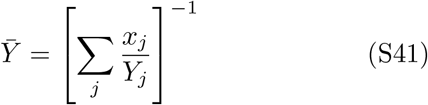

is the effective yield of the population, and

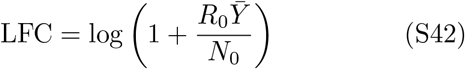

is the log fold-change of the total biomass, which depends on the initial absolute abundance *N*_0_ of all genotypes and the initial concentration of resources *R*_0_.

Equation (S38) shows how each competing genotype influences the saturation time *t*_sat_ and thus the LFCs of all other genotypes via main text Eq. (16). For example, adding a mutant with slow growth rate increases the effective doubling time (Eq. (S40)), while a mutant with long lag time will increase the effective lag time (Eq. (S39)). Genotypes also influence the saturation time (Eq. (S38)) through the effective yield 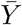, which is the harmonic average of yields for all genotypes (Eq. (S41)). The harmonic average means that adding mutants with low biomass yield *Y*_*j*_ can significantly shorten the saturation time, but mutants that are more efficient (high *Y*_*j*_) have little influence on the duration of the growth cycle. Note that genotypes must be at sufficiently high relative abundance to significantly influence the effective population traits, since each the contribution of each genotype *j* is weighted by its relative abundance *x*_*j*_.

For bulk competition experiments, the saturation time of the culture can be expressed as an interpolation between the traits of the wild-type and the mutant library. We rewrite Eq. (S39) and Eq. (S40) as function of mutant library traits

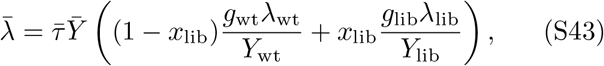

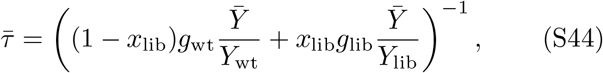

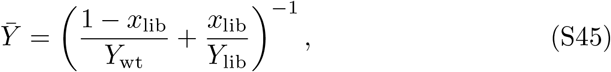

where *g*_wt_ is the wild-type growth rate, *λ*_wt_ is the wildtype lag time,

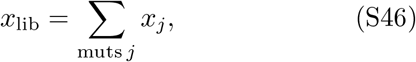

is the mutant library abundance at the beginning of the bulk competition,

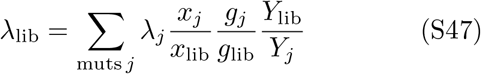

is the mutant library lag time,

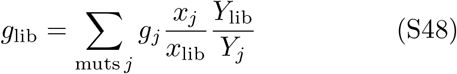

is the mutant library growth rate, and

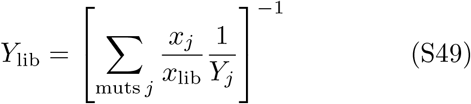

is the mutant library yield. Note that the effective traits in Eq. (S47)–(S49) are independent under a rescaling of the mutant library abundance *x*_lib_ as long as the relative proportion of the individual lineages *x*_*j*_*/x*_lib_ stays constant.

To simplify calculations, we sometimes assume that the mutants have variation in lag time and growth rate, but no variation in biomass yield (*Y*_*i*_ = *Y*_*j*_ for all *i* and *j*). Under this assumptions, we can rewrite Eq. (S43) and Eq. (S44) as

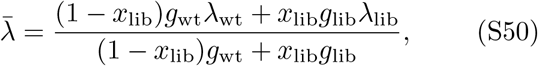

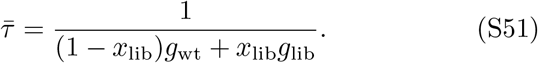

## S8. EPISTASIS DETECTION FOR DOUBLE MUTANTS USING RELATIVE FITNESS PER-CYCLE AND PER-GENERATION

In this section, we discuss the patterns of epistasis observed in relative fitness per-cycle and relative fitness per-generation for the systematic scan of double mutants in our model of population dynamics (Methods). For relative fitness per-cycle, main text Fig. 3C shows that epistasis occurs if and only if one of the mutations involves a change in growth rate. This is because the selection on growth rate in the per-cycle statistic depends on the duration of the growth cycle [9]

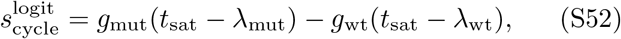

where *t*_sat_ is the saturation time, *t*_sat_ − *λ*_mut_ is the time of growth for the mutant strain and *t*_sat_ − *λ*_wt_ is the time of growth for the wild-type strain. At high relative abundance of the mutant, the saturation time *t*_sat_ depends on all three mutant traits (Eq. (S38)), leading to quadratic mutant trait terms in Eq. (S52) and non-zero epistasis in main text Eq. (17). For example, a double mutant in growth rate (+*g*, +*g*) has lower than expected relative fitness per-cycle (negative epistasis in Eq. (17)) because the second growth rate mutation shortens the growth window *t*_sat_ − *λ*_mut_ in Eq. (S52) (see growth curves in Fig. S9; row A). Intuitively, there is no epistasis between lag time and yield because changes to lag time play out early in the growth cycle and changes in yield only change the final time point of growth (see Fig. S11). Note that here we have chosen examples where both mutants exit the lag phase before saturation – for deleterious mutations with extremely long lag time, a double mutant (+*λ*, +*λ*) eventually shows epistasis in relative fitness per-cycle since the wild-type growth advantage is bounded from above due to finite resources.

Using relative fitness per-generation *s*_gen_, main text Fig. 3D shows that epistasis occurs if and only one of the mutations involves lag time, a pattern that is distinct from the per-cycle statistic (compare Fig. 3C and D). This is because the relative fitness per-generation uses the wild-type LFC as a normalizing time scale (main text Eq. (10)) and nonlinearities in this time scale with respect to the mutant traits can modify or create new epistasis. For example, epistasis is modified in the per-generation statistic for a double mutant in growth rate and lag time (e.g., (+*g*, − *λ*); Fig. S9, row D). There is positive epistasis in relative fitness per-cycle, since the shorter lag time leads to a longer mutant growth time which amplifies selection on growth rate (compare Eq. (S52)), but there is even higher positive epistasis in the per-generation statistic, because the double mutant growth is spread over less wild-type generations compared to the single mutant growth. Sometimes, the nonlinearity in the time-scale LFC_wt_ works to remove epistasis. For example, a double-mutant of growth rate (+*g*, +*g*) (Fig. S9, row A) is epistatic in the per-cycle statistic but non-epistatic in the per-generation statistic

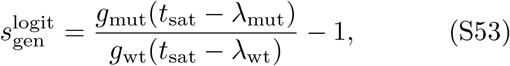

where the mutant growth time *t*_sat_ − *λ*_mut_ and the wild-type *t*_sat_ − *λ*_wt_ are both reduced but happen to cancel. Finally, the mutant-dependent time-scale LFC_wt_ of the per-generation statistic also introduces new epistasis between lag time and yield (Fig. S11), as well as for mutants with two lag time mutations (Fig. S11, row C and Fig. S12, row D). These mutations have no epistasis in relative fitness per-cycle but the normalization by the wild-type LFC in main text Eq. (10) introduces an additional dependence on *t*_sat_ that renders fitness a quadratic function of lag time.

We repeat the fitness estimates using a 1% mutant relative abundance in the simulated competition experiments (Fig. S15). At this low mutant abundance, Fig. S15B shows the epistasis in relative fitness per-cycle *s*_cycle_ is limited to combinations of mutants that perturb growth rate and lag time. Note that in this low abundance limit, the relative fitness per-generation *s*_gen_ agrees on the presence and absence of epistasis (Fig. S15C), simply because the wild-type LFC is approximately constant across mutants.

## S9. ANALYSIS OF FITNESS TRAJECTORIES FROM THE LONG-TERM EVOLUTION EXPERIMENT

A previous analysis of the Long-Term Evolution Experiment (LTEE) performed by Wiser et al. [11] found that evolved populations of *Escherichia coli* increased in relative fitness over 50,000 generations without converging to a maximum fitness. Here we reanalyze the same data by directly comparing the relative fitness (under the logit encoding) per-cycle 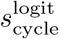 and per-generation 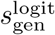 to see if the choice of fitness statistic changes the conclusion. The experimental protocol of the LTEE has been described elsewhere [5, 12], but we briefly summarize the main aspects: Starting with a single ancestral strain of *E. coli*, 12 replicate populations were inoculated in 1988 and are perpetually grown in batch cultures with serial transfers, such that 1% of the population biomass is transferred to fresh growth medium each day. Samples from each replicate population are stored every 500 generations, leading to a record of evolved populations over time [12].

Previous work performed competition experiments between the ancestral population and each evolved population (every 500–2000 generations) by combining them in equal proportions and growing them over a single batch culture growth cycle, with measurements of their initial and final absolute abundances taken by colony counting [11]. This data has been prepared in a convenient format by Good et al. [13] and is available for download at https://github.com/benjaminhgood/LTEE-metagenomic/blob/master/additional_data/Concatenated.LTEE.data.all.csv. The total dataset lists *n* = 928 competition experiments. We remove *n* = 6 entries because either the initial or final abundances are missing.

For a few of the 12 populations, the time series is truncated: population Ara+6 has competition measurements up to generation 4000, population Ara-2 has competition measurements up to generation 30,000, and population Ara+2 has competition measurements up to generation 32,000 [11]. Note that the evolved population tested in these competitions is not a single genotype, but a sample of many genotypes that were present in that evolving population at that time. From these values of absolute abundance, we compute the log fold-change (LFC) of the evolved and ancestral populations in each competition and then calculate the evolved population’s relative fitness per-cycle (main text Eq. (9)) and per-generation (main text Eq. (10)). The original data set has two to four replicate measurements for each evolved sample, corresponding to a repeat of the competition experiment at a different day [11]. Initially, we collect all competition experiments into a single data set (*n* = 928 competitions) and find that relative fitness per-generation and per-cycle differ in the ranking of these competitions (main text Fig. 4B, Fig. S17A). For example, one competition is ranked 261 positions lower in relative fitness per-generation than per-cycle (where higher rank indicates higher fitness). The scatter occurs because the biomass yield evolves downward over time [14, 15]. We can understand the ranking mismatch from the underlying LFCs in the competition experiment (Fig. S17B), that show considerable scatter, with some mutant-wild-type pairs in a positive covariation (compare Fig. S17B and main text Fig. 1D). The scatter occurs because the biomass yield evolves downward over time [14, 15], shifting the wild-type LFCs downward in 1:1 competitions with the evolved populations (horizontal trend across time points in Fig. S17B).

We can also construct a time series of relative fitness for each population and compare fitness rankings at a individual time points. For each of the 12 populations in the LTEE, we define the relative fitness per-generation 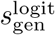 at time *t* by averaging the value 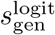 across all competition experiments with the frozen sample from time *t*. Simlarly, we define a time series of relative fitness per-cycle 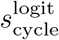 for each population. In summary, we can pool all time series into a single data set (with the truncation described above) and test how the two statistics rank the 12 populations at any point in the experiment. We find that the mismatch between relative fitness per-cycle and per-generation is consistently low at all time points (main text Fig. 4A), with a few exceptions (for example, at generation 4,000 the two statistics disagree on the top six populations).

A key result from previous analysis by Wiser et al. [11] on this data set is that the evolving populations increase indefinitely in relative fitness, rather than leveling off at some maximum fitness value. The empirical analysis by Wiser et al. [11] tested the long-term trend by fitting the statistic 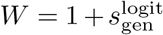 to a hyperbolic model of the time series:

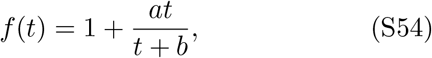

where *t* is the evolutionary time point at which the evolved population is measured against its ancestor, and *f* (*t*) is the fitness statistic measured from that competition (here fitted to the measured values of *W*). The important feature of the hyperbolic model is that it assumes that relative fitness saturates at a maximum fitness over long times (lim_*t*⟶∞_ *f* (*t*) = 1 + *a*). To contrast this model, they also tested a power law

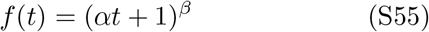

under which fitness increases without bound over long times (lim_*t*⟶∞_ *f* (*t*) = ∞). To repeat this analysis with the fitness statistics 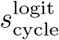 and 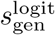 used in this article, we must adjust the models to account for the fact that *W* takes 1 as its neutral value (occurring at *t* = 0 by definition) while 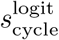 and 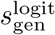 are zero under neutrality. Thus we use

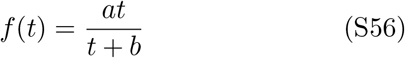

as the hyperbolic model and

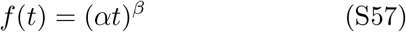

as the power law model.

As a control against the original analysis of Wiser et al. [11], we first perform our own fit of the time series of the relative fitness per-generation 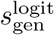 to the hyperbolic (Eq. (S56)) and the power law (Eq. (S57)) models. Wiser et al. compared their two models using the Bayesian Information Criterion but since the models have the same number of parameters this is mathematically equivalent to comparing the values of *R*^2^. Figure S16A shows that in our analysis, the power law model has a higher quality of fit (*R*^2^ = 0.701) than the hyperbolic model does (*R*^2^ = 0.682) for the fitness statistic 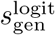, consistent with the original result by Wiser et al. for the statistic 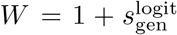. Since the equations for the hyperbolic model only differ in the constant offset (compare Eqs. (S54) and (S56)), our fit of 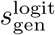 to Eq. (S56) is mathematically equivalent to the fit of *W* to Eq. (S54). Therefore, our fit should give the exact same results as the original publication [11] in the case of the hyperbolic model (although we could not find the fitted values *a, b* documented in the original publication so we were unable to check them). This is not true for the power law, because the modified power law used by Wiser et al. (Eq. (S55)) includes the offset within the parentheses, rather than as an added constant outside (compare Eq. (S55) to Eq. (S57)). This means that the fit of *W* to the power law with an initial value of 1 (Eq. (S55), fitted values *α* = 0.00515, *β* = 0.0950; matching [11]) is different from the fit of 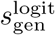 to the power law with an initial value of zero (Eq. (S57), fitted values *α* = 0.000007, *β* = 0.299891).

We next perform a new analysis by calculating the relative fitness per-cycle 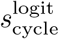 from the same time series data (12 time series pooled together, one for each line) and fitting this fitness statistic to the same hyperbolic (Eq. (S56)) and power law (Eq. (S57)) models. As shown in Fig. S16B, we find that the power law model outperforms the hyperbolic model for the per-cycle fitness, consistent with the model performance for the per-generation fitness. This suggests that Wiser et al.’s original conclusion about fitness increasing without bound [11] is robust to the choice of fitness statistic.

As we previously mentioned, the fitness measurements of all replicate populations are not uniform across time (Fig. S16A): there are fewer fitness measurements at late time points (generation 34,000 and higher) because three populations were eventually excluded from the fitness measurements [11]. To further corroborate our results, we thus repeat the model fits using a single time series of the evolved fitness, rather than fitting the models to a all 12 fitness time series simultaneously. We calculate the average fitness per-generation 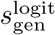 and per-cycle 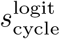 in the evolution experiment as the average across all 12 populations at each time point and fit this population-averaged time series to the hyperbolic (Eq. (S56)) and power law (Eq. (S57)) models (Wiser et al. refer to this as a fit to the “grand mean”). Figure S16C,D shows that the power law still has a better quality of fit than the hyperbolic model does, and we thus conclude that the increasing fitness trend reported by Wiser et al. [11] is also robust to the uneven distribution of measurements over time.

We note that the quality of fit *R* reported in the main text of Wiser et al. [11] differs from the quality of fit we show on our plots. While both studies performed the fits of the models on all 12 populations simultaneously, Wiser et al. [11] evaluated the quality of fit by calculating *R* for the fitted model (blue and pink lines in Fig. S16A) against the fitness time series averaged over replicates (grey points in Fig. S16C). That is, they fit the model to the data without averaging over population but calculate quality of fit using the population-averaged data. Since we believe this was inconsistent, we have followed a more standard approach of calculating the quality of fit on the same input data used for the fit (i.e., correlating the blue and pink lines in Fig. S16A with grey points in the same plot). As a consequence, the correlations between the fitted models and data reported in the original publication (hyperbolic *R* = 0.969, power law *R* = 0.986 [11]) are systematically higher than the values we find in our re-analysis (hyperbolic *R* = 0.826, power law: *R* = 0.837; take square root of the values in Fig. S16A). Although the long-term fitness dynamics in the case of the LTEE are robust to the choice of fitness statistic, we note that it is possible to construct scenarios of microbial growth trait evolution where relative fitness saturates in one statistic but not in the other (main text Fig. 3E-F). For theory work that on the long-term trend and other models than the power law or hyperbolic model tested here, see refs. [16–20].

## S10. TESTING AUC AND OTHER FITNESS POTENTIALS USING SIMULATED COMPETITION EXPERIMENTS

In this section we explain our tests of estimated fitness potentials against true relative fitness in simulated competition experiments. We first focus on the area under the growth curve (AUC), which is defined as

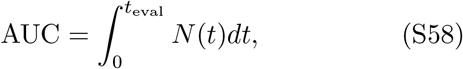

where *N* (*t*) is a growth curve of absolute abundance (or a proxy such as optical density) and *t*_eval_ is a cutoff time for evaluating the area. Many previous studies [21–25] and growth curve analysis packages [26] have used this definition. The idea of AUC is that, unlike estimated fitness potentials that only account for individual traits of a genotype’s growth (e.g., growth rate or lag time; see below), the AUC literally integrates the whole growth dynamics into a single number. For example, both fast growth rate and short lag time are manifested in greater AUC. Note that while one can attempt to use the AUC as an approximate fitness potential (as we investigate here), it is not a measure of absolute fitness (Sec. S2) since it is insufficient to predict changes in absolute abundance (i.e., the area under the growth curve does not determine the change in absolute abundance from beginning to end).

To compute the AUC for the set of single-gene deletion genotypes in our data set, we first simulate a growth curve for each genotype under the population dynamics model in main text Eq. (12) with the traits estimated from the yeast single-gene knockout data (Methods). We use simulated growth curves, with trait values inferred from the measured growth curves, rather than the actual measured growth curves since we are comparing the AUCs to relative fitness also from simulations of competitions; we do not have actual competition data for these genotypes, so it would be an apples-to-oranges comparison if we used AUCs from the actual growth curve data. Furthermore, in empirical growth curves, the AUC is also influenced by technical variation in the initial biomass *N*_0_ and the initial concentration of resources *R*_0_, so using simulations removes these effects.

Figure S20A shows the distribution of saturation times *t*_sat_ (as defined for the population dynamics model in Methods) numerically calculated for all simulated growth curves of the deletion mutants. We calculate the AUC for each growth curve using Eq. (S58) with an evaluation time of *t*_eval_ = 16 hours, since that includes the stationary phase in the vast majority of our simulated growth curves (Fig. S20A). From the mutant’s AUC we calculate the AUC-based estimator of the mutant’s relative fitness

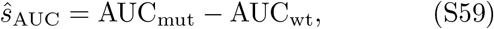

where AUC_wt_ is the AUC for a simulated growth curve of the wild-type (using the median wild-type traits in our database; see Methods). We then simulate a single growth cycle of the mutant competing against the wild-type (Eq. (12) with equal initial abundance of the mutant and wild-type) to calculate the mutant’s “true” (under the model assumptions) relative fitness per-cycle with the logit encoding 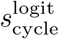. We simulate both the coculture and the monoculture growth dynamics with the same initial biomass *N*_0_ = 0.05 OD and resource concentration *R*_0_ = 111 mM [6]. Repeating this analysis for all mutant genotypes in the data set, we calculate the Spearman rank correlation between the AUC-predicted relative fitness from monoculture *ŝ*_AUC_ and the true relative fitness in competition 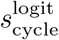 (Fig. S19, column C).

The success of the AUC depends on the evaluation time *t*_eval_, which sets the time window from which information is captured from the growth curve by the integral (Eq. (S58)). Figure S20B–D shows the correlation between the relative fitness in coculture 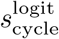 and the AUC estimator *ŝ*_AUC_ (Eq. (S59)) for three values of *t*_eval_: the mean saturation time of all genotypes in monoculture (*t*_eval_ ≈ 13 hours), a significant longer value (*t*_eval_ ≈ 24 hours), and an intermediate value (*t*_eval_ = 16 hours; used for Fig. S19). For short evaluation times (*t*_eval_ ≈ 13 hours), the AUC underestimates the fitness of mutants with long lag but fast growth, which leads to a nonlinear relationship between the AUC estimator and the true relative fitness (compare Fig. S20B and Fig. S20C). For long evaluation times (*t*_eval_ = 24 hours), there is greater scatter between the AUC predictor and true relative fitness for the highest fitness mutants (compare spread in Fig. S20C and Fig. S20D). Intuitively, these mutants have short lag time or fast growth rate and saturate early in monoculture, so their AUC values are effectively set by the biomass yield, which has no predictive value on the competition outcome. In summary, Fig. S20B–D shows a tradeoff between accurately ranking highly-deleterious mutants (which need long *t*_eval_) and ranking highly-beneficial mutants (which need short *t*_eval_).

Besides AUC it is possible to use other features of the monoculture growth curve as approximate fitness potentials. For example, one can use the monoculture growth rates alone as fitness potentials:

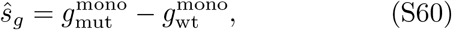

or the monoculture lag times:

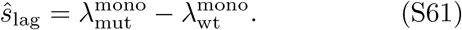

Another possible fitness potential is the absolute abundance at saturation, which in our model of population dynamics (Eq. (12)) is proportional to the biomass yield:

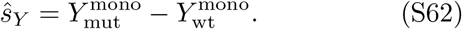

Finally, one can also use the difference in the monoculture log fold-changes (LFCs):

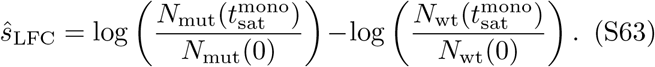

This looks similar to the definition of relative fitness per-cycle (main text Eq. (9)) but is distinct because it uses the LFCs in monoculture rather than the true LFCs in coculture (which may be different).

We test each of these fitness potentials against the relative fitness in pairwise competition, using different input data sets of trait variation and the “GNU Parallel” command to speed up the simulation process [27]. Figure S19 rows B and C show that growth rate and lag time can act as perfect fitness potentials if that trait is the only trait with variation across mutants. This is because the relative fitness is proportional to differences in each of these traits when they are the only source of variation (see Sec. S14). Section S14 also shows that differences in biomass yield (*Y*_mut_ − *Y*_wt_) have no effect on fitness by themselves, which is why the biomass yield of the strains in monoculture is a poor fitness potential to estimate relative fitness. This large but neutral variation in the biomass yield across mutants in our data sets means that the LFC is also a poor fitness potential.

All of these trait-based fitness potentials are outperformed by the AUC, which provides the best approximation of the mutant fitness ranking in coculture under realistic trait variation (Fig. S19, row A). More broadly, it is important to treat absolute and relative fitness, as well as fitness potentials, as distinct concepts, serving different purposes [28, 29]. As we show here in simulation (Fig. S19), and others have shown in experiments [22, 23, 30, 31], measuring fitness potentials is not enough to demonstrate that a mutant genotype will out-compete the wild-type.

## S11. COARSE-GRAINING PAIRWISE RELATIVE FITNESS IN MULTI-GENOTYPE POPULATIONS

In this section, we point out the specific advantages of the logit-encoding for coarse-graining pairwise relative fitness in bulk competition experiments. While the pair-wise relative fitness is defined for any encoding *m*, the logit encoding endows it with some convenient mathematical properties not shared by other encodings (e.g., the log encoding). The logit encoding of the pairwise relative abundance has the property

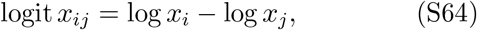

meaning that it is antisymmetric under exchange of the indices *i* and *j* (logit *x*_*ij*_ = −logit *x*_*ji*_) and additive across pairs of indices (logit *x*_*ij*_ = logit *x*_*ik*_ + logit *x*_*kj*_). Since the logit-encoded pairwise relative fitness is just the time derivative of the logit function (Eq. (21)), it carries equivalent properties of antisymmetry and additivity:

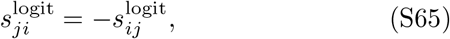

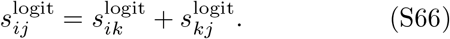

We also note that the logit encoding of the pairwise relative abundance has the property:

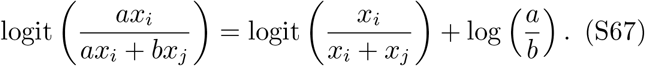

Rescaling the relative abundances of either genotype thus does not change the pairwise relative fitness (since it only shifts the logit by a constant, which does not affect its derivative). This means that pairwise relative fitness is an “intensive” property of a genotype, analogous to intensive properties in statistical mechanics (such as temperature) that do not scale with system size. For example, if we split a mutant genotype into two subgroups (e.g., differentiated by a neutral marker), the pairwise relative fitness of each mutant subgroup with respect to the wild-type will be the same as the pairwise relative fitness of the mutant genotype as a whole compared to the wild-type. In contrast, the logit-encoded total relative fitness does not satisfy this property since logit(*ax*_*i*_) = logit *x*_*i*_+ constant.

When the encoding *m* is the logit function, the pairwise relative fitness per-cycle still satisfies the above properties (antisymmetry and additivity with respect to indices *i, j*, and invariance under relative abundance rescaling) since those are properties of the underlying logit encoding. This is also apparent from interpreting the per-cycle fitness statistic as an integral of the instantaneous statistic:

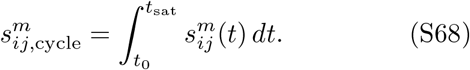

## S12. THE RELATIVE FITNESS BETWEEN COARSE-GRAINED GROUPS OF GENOTYPES

In this section, we generalize the concept of pairwise relative fitness to pairs of genotype groups rather than pairs of individual genotypes. In a multi-genotype population with subsets of genotypes *A* and ℬ, define

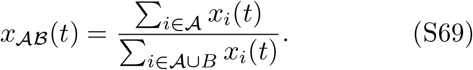

as the relative abundance of genotypes compared to genotypes at time *t*, where *A B* is the union of the two index sets. Analogous with main text Eq. (21), we define the fitness of group *A* relative to group ℬ as

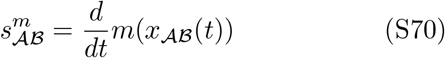

for an encoding *m*. Under the logit encoding, it turns out that the fitness between these two groups can be conveniently expressed as a weighted average of the pairwise fitness between the member genotypes in each group:

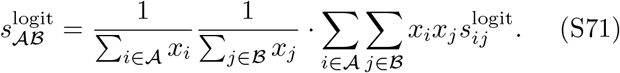

To prove Eq. (S71), we first note that

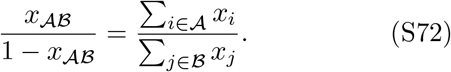

Thus we can rewrite the logit-encoded relative pairwise fitness between *A* and ℬ as

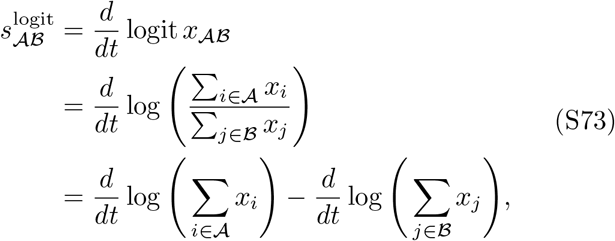

where we have invoked Eq. (S72) on the second line. We can expand each term on the right-hand side of Eq. (S73) as

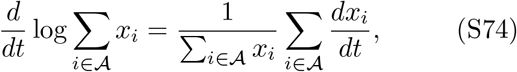

and then insert Eq. (S74) into Eq. (S73) to obtain

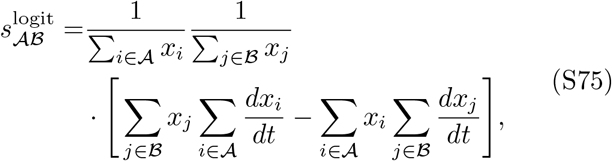

where we have collected the normalization factors (sums over relative abundances in *A* and ℬ) as a single pre-factor. We then rewrite each product of sums in Eq. (S75) as

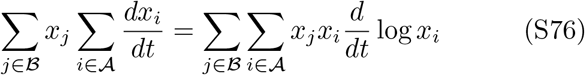

to finally obtain

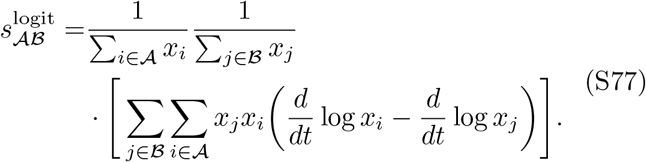

Identifying the term in the inner parentheses as the pair-wise selection coefficient 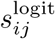 (main text Eq. (21)) then results in Eq. (S71).

Equation (S71) establishes that the relative fitness between a pair of genotype groups is a weighted sum of relative fitnesses between individual pairs of genotypes in those groups, but this holds for relative fitness defined at an instant in time (since it is based on derivatives). For relative fitness defined over a finite time interval (e.g., a growth cycle in batch culture), an analogous but approximate result holds. We first write the relative fitness over a growth cycle time interval as an integral over the instantaneous relative fitness (inserting Eq. (S71) into Eq. (S68)):

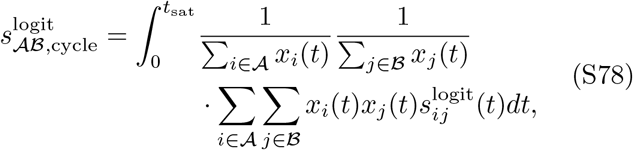

where *t*_sat_ is the end time of the growth cycle (Methods, Sec. S7). The integral in Eq. (S78) is difficult to calculate as the relative abundance trajectories *x*_*i*_(*t*), *x*_*j*_(*t*) depend on the relative fitness of the genotypes 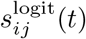 in a non-trivial way. Instead, we make the approximation that the relative abundances do not change significantly over time of the growth cycle (*x*_*i*_(*t*) ≈ *x*_*i*_(0)) and can thus pass the integral through the sums in Eq. (S78) to show that the per-cycle relative fitness between a pair of genotype groups is approximately also a weighted sum of per-cycle relative fitnesses:

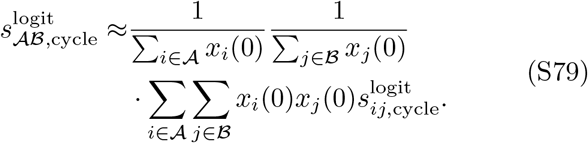

Conceptually, assuming the relative abundances are approximately constant over the growth cycle is equivalent to assuming selection is weak; one can also show this mathematically by expressing the relative abundances *x*_*i*_(*t*) in terms of the pairwise relative fitnesses 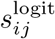 Eq. (S78) and keeping only terms to leading order in 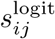.

We finally note that the total relative fitness (instantaneous Eq. (19) and per-cycle Eq. (22) in the main text) is a special case of the relative fitness between groups (Eqs. (S71) and (S79)) where *A* is the single genotype *i* and ℬ is all other genotypes besides *i*:

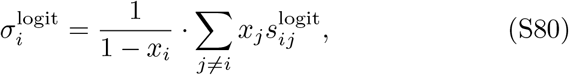

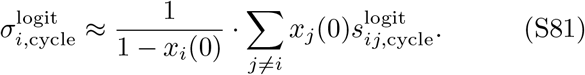

## S13. FITNESS ERROR FROM THE FRAME OF REFERENCE IN BULK COMPETITION EXPERIMENTS

Here we calculate the error that arises from measuring the total relative fitness of each mutant in a bulk competition experiment of a mutant library, rather than the pairwise relative fitness between each mutant and the wild-type. We call this difference the *error from the frame of reference*, the frame of reference being either the whole population in total fitness or the wild-type in pairwise fitness. Note that this is an error between two different fitness quantifications of the same bulk competition experiment; Sec. S15 addresses the error (arising from higher-order interactions) between fitness quantifications in bulk versus pairwise competition experiments. Here we only consider relative fitness under the logit encoding and measured per growth cycle, so we drop these labels to simplify notation.

Consider a bulk competition experiment of a wild-type and a library of large number of mutants over a single batch growth cycle of time *t*_sat_ (Methods). The total relative fitness of mutant *i* is (main text Eq. (22))

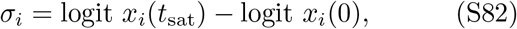

while its pairwise relative fitness compared to the wild-type is (main text Eq. (23))

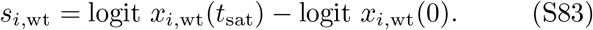

Using the coarse-graining rules from Sec. S12 (namely Eqs. (S79) and (S81)), we can express the total relative fitness of mutant *i* as a weighted sum of the pairwise relative fitnesses between the mutant and the wild-type and between the mutant and the rest of the mutant library:

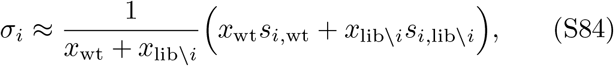

where the notation lib *i* refers to the mutant library excluding the mutant *i*. The approximation here is due to our assumption in Eq. (S79) that selection is weak in enough that the relative abundances of genotypes do not change too much over the growth cycle. Since we can rewrite the pairwise relative fitness between mutant *i* and the rest of the library as a difference between *i* and the wild-type and the rest of the library and the wild-type (using Eq. (S66))

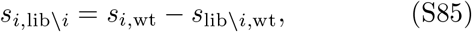

we insert this into Eq. (S84) to obtain

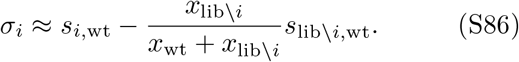

The difference between the total and pairwise relative fitness is therefore

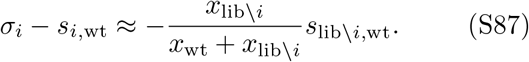

Since mutant libraries in these experiments typically contain hundreds or thousands of mutants, the contribution of a single mutant *i* is small and thus we can assume that the properties of the mutant library excluding mutant *i* (lib *\ i*) are approximately the same as the library as a whole (so that *x*_lib*\i*_ ≈ *x*_lib_, *s*_lib*\i*,wt_ ≈ *s*_lib,wt_). This allows us to further simplify the error as

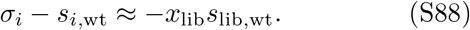

Equation (S88) shows why this offset between the total and pairwise relative fitnesses is approximately independent of the focal mutant *i* (hence the constant shift for all mutant points in main text Fig. 5C). The sign of the error from the frame of reference depends on the mean fitness of the mutant library: a mutant library that is overall deleterious relative to the wild-type (*s*_lib,wt_ *<* 0) causes the total relative fitness for a mutant *σ*_*i*_ to over-estimate the mutant’s pairwise relative fitness *s*_*i*,wt_. Intuitively, this is because the total relative fitness is comparing the mutant to a mixed population of wild-type and other mutants, which are on average worse competitors than the wild-type, which thus makes the mutant appear to be better than if it is just compared to the wild-type alone (compare top and bottom panel in main text Fig. 5B). Equation (S88) also shows that the error from the frame of reference can be reduced if the mutant library is neutral relative to the wild-type (*s*_lib,wt_ = 0) or the mutant library has small relative abundance in the culture biomass (*x*_lib_ 1). In bulk competition experiments with barcoded mutant libraries, these assumptions are often not met since the mutant libraries tend to be overall deleterious (as in our simulated bulk competition for the yeast single-gene knockouts, see Fig. S6) and are inoculated at a high relative abundance [32–36] Since this makes the error in Eq. (S88) significant, we instead recommend including barcoded wild-type strains as references in the bulk competition, so that pairwise fitness can be quantified relative to that (Eq. (S83)) rather than using the total relative fitness. By using a mix of bar-coded wild-type cells and non-barcoded wild-type cells it is further possible to optimize this protocol and save on sequencing investment [37].

Finally, we want to point out a difference between the best practice we recommend here (Discussion) and a wide-spread practice in estimating fitness estimates in bulk competition experiments. In practice, transposon-seq experiments that grow the mutant library by itself start with an estimate of total relative fitness, and then subtract the median total relative fitness of the knockouts [33–35, 38] or a mean total relative fitness [32, 39, 40]. However, these corrections are not explicitly founded in the choice of a reference group (like a set of neutral genotypes or a wild-type), making the correction appear ad-hoc [33, 35, 38], or the reference is a strain that is not part of the culture, like in the fitness estimates for barcoded lineages that are evaluated against the initial ancestor without that ancestor actually being present [32, 39, 40]. To make things more confusing, even those studies that do include a wild-type then describe their method as an estimate of total relative fitness of the mutant under the log encoding, subtracted by the total relative fitness of the wild-type under the log-encoding [37, 41, 42]. The result is a pairwise relative fitness under the logit-encoding (Eq. (25)) but presenting it this way obscures that choice of encoding and the relationship to the classic, logit-based selection coefficient used in pairwise competition experiments (Eq. (7)). We hope that our framework can provide more clarity: The choice of the reference group happens at the level of relative abundance, by calculating a pairwise relative abundance (Eq. (20) or Eq. (S69)), and this removes the need for any correction on the fitness values themselves.

## S14. PAIRWISE RELATIVE FITNESS USING AN EXPLICIT MODEL OF POPULATION DYNAMICS

In the model of population dynamics (main text Methods, Eq. (12)), we can calculate the pairwise relative fitness of genotypes based on the approximation of the saturation time (Sec. S7; [9, 10]). The pairwise relative fitness of genotype *i* relative to genotype *j* (per-cycle and under the logit encoding) is

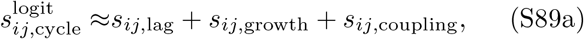

where

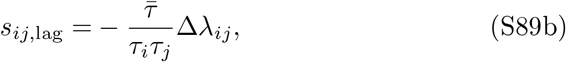

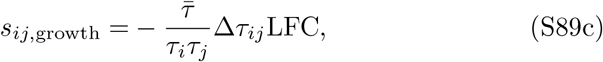

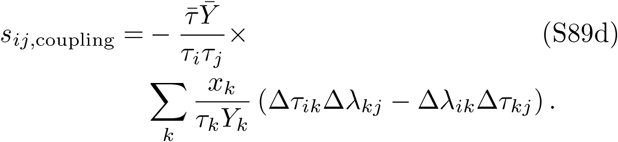

The *e*-fold growth time for genotype *j* is *τ*_*j*_ = 1*/g*_*j*_, and the terms Δ*λ*_*ij*_ = *λ*_*i*_ − *λ*_*j*_ and Δ*τ*_*ij*_ = *τ*_*i*_ − *τ*_*j*_ are the differences between the two genotypes lag times and growth times. Since the terms in Eq. (S89d) depend on the covariation between growth and lag, we interpret these terms as couplings between the growth and lag phases; they are zero if only two genotypes are present.

## S15. FITNESS ERROR FROM HIGHER-ORDER INTERACTIONS BETWEEN PAIRWISE AND BULK COMPETITION EXPERIMENTS

In this section, we calculate the error in relative fitness of a mutant arising from higher-order interactions in bulk competition experiments with large mutant libraries, compared to the “true” relative fitness in pair-wise competitions between just the focal mutant and wild-type alone. Here we only consider relative fitness under the logit encoding and measured per growth cycle, so we drop these labels to simplify notation. Let the pairwise relative fitness of mutant *i* compared to the wild-type (main text Eq. (23)) be

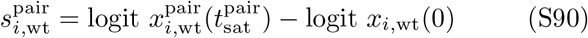

in the pairwise competition with the wild-type alone, and

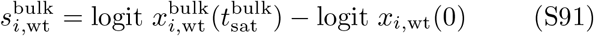

in the bulk competition with all other mutants in the library. The superscripts pair and bulk indicate that the dynamics of *x*_*i*,wt_(*t*) and the saturation time *t*_sat_ may be different in the two competitions. The difference between these two measurements of relative fitness is

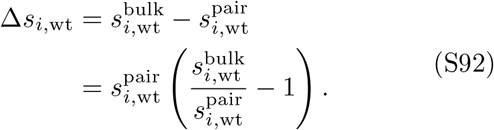

Since the difference between the bulk and pairwise competitions is the presence of the other mutants in the library, we interpret this difference as the *fitness error from higher-order interactions* among the mutants.

We now calculate how this error depends on the underlying growth traits of the genotypes using the population dynamics model (main text Methods, Eq. (12)); and the explicit expression for relative fitness from Sec. S14. Based on the approximate pairwise relative fitness in this model (Eq. (S89)), the relative fitness in the pairwise competition is the sum of two terms

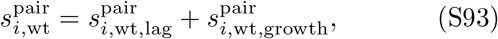

while the relative fitness in the bulk competition is the sum of three terms

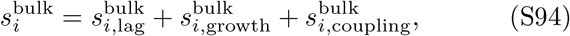

where the third term represents the coupling between growth and lag phases present only in populations with more than two genotypes (Eq. (S89d)). We define the higher-order effects on the selection for lag time as

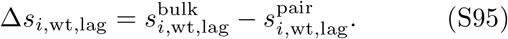

Using Eq. (S89b), we can express this in terms of the underlying traits as

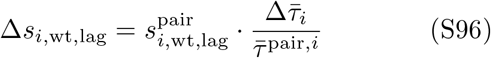

where 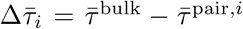 is the difference in effective *e*-fold growth times (Eq. (S40)) in the bulk competition and in the pairwise competition of mutant *i* and the wild-type.

We similarly define the higher-order effects in the selection on growth rate as

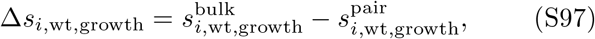

and calculate 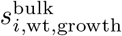 and 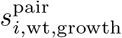 from Eq. (S89c) to get the expression

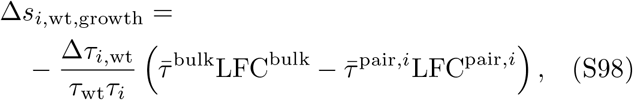

where LFC^bulk^ is the log fold-change (Eq. (S42)) of the total biomass in the bulk competition and LFC^pair,*i*^ is the log fold-change of the total biomass in the pairwise competition between mutant *i* and the wild-type. We note that total biomass growth in the bulk competition LFC^bulk^ depends on the mutant library through the effective biomass yield (Eq. (S41)), but this dependence is weak because the yield enters only logarithmically into Eq. (S42). We thus assume that the bulk and pair-wise competition have equal biomass growth (LFC^bulk^ LFC^pair,*i*^) for all mutants *i* such that the higher-order effect on the growth rate selection (Eq. (S98)) is proportional to the increase in the mean doubling time

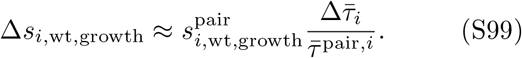

Since this has the same form as the higher-order effect on lag time selection in Eq. (S96), we can combine Eq. (S99) and Eq. (S96)) into

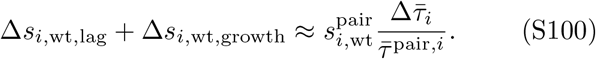

Because the effective growth time in the pairwise competition 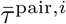 is approximately just the effective growth time of the wild-type alone *τ*_wt_ (assuming the mutant is competed against it at low initial relative abundance), this means that

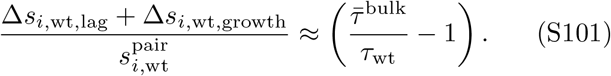

That is, the relative error of lag and growth selection from higher-order interactions is approximately independent of the individual mutant, but rather depends on properties of the wild-type (*τ*_wt_) and the whole mutant library (through 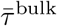). This is why we observe an approximately constant slope across all mutants in Fig. (dark orange points) when comparing the error against the pairwise relative fitness. The slight deviation from the constant slope is due to our approximation that the LFCs are the same between the bulk and pairwise competitions; this is a good approximation for the yeast deletion library but not exactly true, and hence causes a slightly different scaling between the growth Δ*s*_growth_ and lag Δ*s*_lag_ error terms compared to Eq. (S101).

Finally, we note that the growth-lag coupling terms 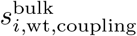 do not have a simple scaling with pairwise relative fitness since they depend quadratically on trait differences; this is shown as the bright orange points in Fig. S22. In the main text, we therefore refer to Δ*s*_*i*,wt,lag_ + Δ*s*_*i*,wt,growth_ as the fitness-dependent error and the 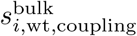 as the fitness-independent error from higher-order interactions.

## S16. CHOOSING THE MUTANT LIBRARY ABUNDANCE IN BULK COMPETITION EXPERIMENTS

Section showed that measuring relative fitness of a mutant in a bulk competition (with a library of other mutants also present) entails an error due to higher-order interactions among the mutants, compared to measuring relative fitness in a pairwise competition consisting of just the mutant and wild-type. Here we show how this error depends on the relative abundance of the mutant library in the bulk competition, so that we can estimate the range of library abundances that keep the error below a desired threshold.

### Calculating the relative error on fitness

Let the absolute error in relative fitness from higher-order interactions for a mutant *i* be Δ*s*_*i*,wt_ (Eq. (S92)). Let the relative fitness of this mutant in a pairwise competition be 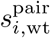 (Eq. (S90)). Since the error Δ*s*_*i*,wt_ depends on the relative abundance *x*_lib_ of the whole mutant library, our goal is to determine what range of *x*_lib_ keeps the error in relative fitness (compared to the “true” relative fitness in the pairwise competition) below a chosen threshold *ϵ*:

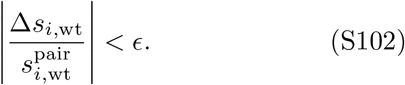

In Sec. we calculated the dependence of Δ*s*_*i*,wt_ on the underlying growth traits. However, since here we are mainly concerned with the dependence on the library abundance *x*_lib_, we present an alternative calculation that better captures that dependence.

We start by pointing out that we can express the relative fitness in pairwise (Eq. (S90)) and bulk competitions (Eq. (S91)) in terms of the saturation times for these competitions, using the explicit solution to the population dynamics model in main text Eq. (15):

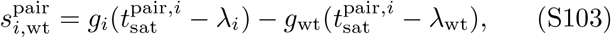

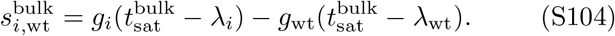

We can thus express the error from higher-order interactions (Eq. (S92)) as

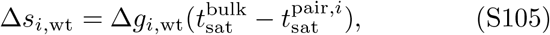

where Δ*g*_*i*,wt_ = *g*_*i*_ − *g*_wt_. Equation (S105) shows that the mutant library affects the fitness of individual mutants 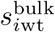 by changing the saturation time. Mathematically, this is equivalent to the results of Sec. S15, which showed how the difference in effective *e*-fold growth times 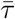 between bulk and pairwise competitions primarily mediated the higher-order effects, but expressed in terms of the saturation time *t*_sat_ (which is not identical to 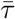 but related through Eq. (S38)). The error in Eq. (S105) is also proportional to the growth rate advantage of the mutant compared to the wild-type. In particular, mutant genotypes that only differ in lag time are not affected by the mutant library since their advantage is accrued once at the beginning of the growth cycle and therefore does not scale with the total time of competition.

According to Eq. (S38), the saturation time *t*_sat_ depends on the effective lag time 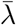 (Eq. (S39)), effective *e*-fold growth time 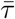 (Eq. (S40)), and the log fold-change LFC (Eq. (S42)) for the competition. To simplify the calculation, we introduce a few assumptions. We assume all mutants have the same yields as the wild-type (*Y*_*i*_ = *Y*_wt_) such that the LFCs in the bulk and pairwise competitions are identical (LFC^bulk^ = LFC^pair^). We also assume that the relative abundance of the mutant in the pairwise competition is small enough that the saturation time in that case is set entirely by the wild-type traits:

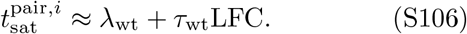

Note that this means that the difference in saturation times is independent of the specific mutant *i*; the only dependence of the mutant *i* on the overall error from higher-order interactions is through the difference in growth rates in Eq. (S105).

We can thus write the fitness error as

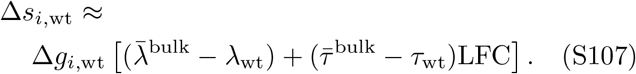

Equation (S107) depends on the mutant library abundance *x*_lib_ through the effective traits 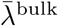 and 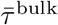 of the bulk competition (Sec. S7). Inserting Eqs. (S50) and (S51) into Eq. (S107), we obtain

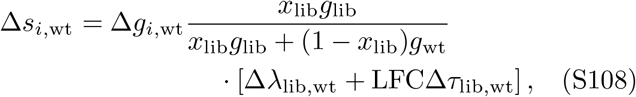

where Δ*λ*_lib,wt_ = *λ*_lib_ − *λ*_wt_ is the difference in lag times, and Δ*τ*_lib,wt_ = *τ*_lib_ − *τ*_wt_ the difference in *e*-fold growth times (reciprocal growth rates), between the library (defined according to Eq. (S47)–(S48)) and the wild-type. All the dependence on the library relative abundance *x*_lib_ is now contained in the fraction outside the square brackets in Eq. (S108).

Since the relative fitness of the mutant library as a whole compared to the wild-type in bulk competition is (using Eq. (S89))

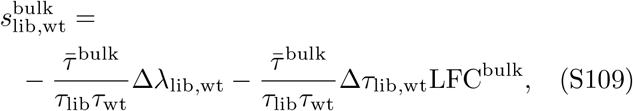

we can rewrite Eq. (S108) as

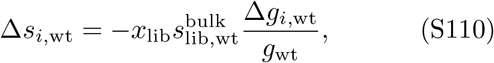

This shows that a mutant library that is neutral relative to the wild-type in the bulk competition 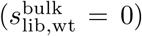 removes the error from higher-order interactions, even when that neutrality is based on a tradeoff between growth rates and lag times (Δ*λ*_lib,wt_ = −Δ*τ*_lib,wt_LFC_bulk_). We note that the error of higher-order interactions (Eq. (S110)) appears similar to the error from the frame of reference (Eq. (S88), but the former includes the relative mutant growth rate Δ*g*_*i*,wt_*/g*_wt_ as an additional pre-factor.

### Calculating the upper bound on relative abundance of the mutant library

To calculate the maximum relative library abundance that keeps the relative error below the threshold *ϵ*, we input the absolute error in Eq. (S108) into the relative error bound of Eq. (S102) and rearrange to isolate the dependence on the library abundance *x*_lib_:

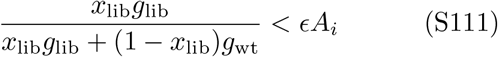

where

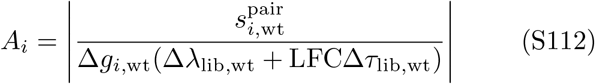

depends on the focal mutant *i* but does not depend on the library abundance *x*_lib_. The left-hand side of Eq. (S111) varies between zero and one as a function of *x*_lib_. This means that if *ϵA*_*i*_ *>* 1, any value of *x*_lib_ will satisfy Eq. (S111), meaning that the relative error on mutant *i* from higher-order interactions will always be less than *ϵ*. For example, this holds when the mutant *i* has the same growth rate as the wild-type (Δ*g*_*i*,wt_ = 0, which causes *A*_*i*_ → ∞), even if it varies in lag time and/or other mutants have variation in growth rates.

We thus next consider the case where *ϵA*_*i*_ *<* 1. We multiply both sides of Eq. (S111) by the denominator on the left-hand side (always positive) to obtain

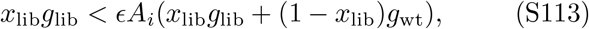

and then collect all the terms involving *x*_lib_ on the left hand side:

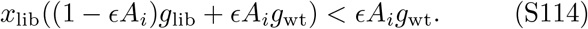

Since *ϵA*_*i*_ *<* 1, the factor multiplying *x*_lib_ on the left-hand side of Eq. (S114) must be positive and thus we can divide both sides to obtain an upper bound on the library abundance *x*_lib_ such that the relative error on fitness is less than *ϵ*:

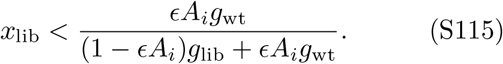

Since *ϵ* will typically be small, we can simplify the right-hand side of Eq. (S115) by approximating it to first-order in *ϵ*:

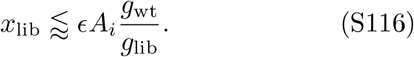

This approximation holds as long as *ϵA*_*i*_(*g*_lib_ − *g*_wt_)*/g*_lib_ ≪1.

The maximum mutant library abundance determined by Eq. (S115) or (S116) is specific to a single mutant genotype *i*, meaning that the relative fitness errors for other mutants may exceed *ϵ* even if the inequality for mutant *i* is satisfied. To keep the relative fitness errors for all mutants less than *ϵ*, we need to choose a library abundance 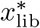 such that

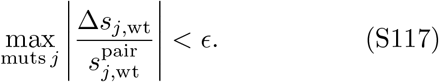

This means that Eq. (S116) must be satisfied for all mutants *j*:

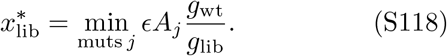

Thus we must determine the minimum value of *A*_*j*_ over all mutants *j*. Using the definition in Eq. (S112), *A*_*j*_ is minimized for the mutant with minimum vale of

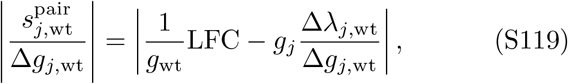

where we have calculated the pairwise relative fitness in Eq. (S103) using 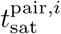 from Eq. (S106). Mutants that minimize Eq. (S119) are those with tradeoffs between lag times and growth rates such that their overall pair-wise relative fitness with respect to the wild-type is zero. This makes sense, since these mutants will have a relative fitness close to zero in the pairwise competition 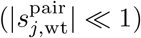, but nonzero fitness relative to the wild-type in the bulk competition as the relative selection on the lag time and growth rate shifts (e.g., because 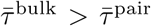 in Eq. (S89)).

### Special case of variation in growth rates only

To determine an even simpler estimate on the maximum mutant library abundance, we consider the special case where genotypes vary only in growth rates and not lag times. Using the definition in Eq. (S112), *A*_*i*_ = *g*_lib_*/*(*g*_wt_ − *g*_lib_) for all mutants *i*, and thus there is a single bound on library abundance for all mutants (using Eq. (S116)):

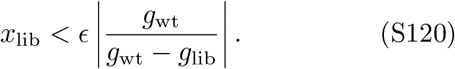

This is the same as Eq. (11) in the main text.

We test Eq. (11) using our simulated competition data for the yeast single-gene deletion library (Methods) with variation in all three traits: lag time, growth rate, and biomass yield (Fig. S4B,C). We compute the growth rates of the mutant library (*g*_lib_ = 0.389 h^−1^ using Eq. (S40)) and the wild-type (*g*_wt_ = 0.406 h^−1^). Using a relative error threshold of *ϵ*= 0.01, the maximum mutant library relative abundance according to main text Eq. (11) is *x*_lib_ = 24.6%. Figure 5D in the main text shows that this mutant library abundance indeed is able to keep the relative error below the threshold for mutants with high relative fitness, but, as expected, this mutant library abundance fails for mutants close to neutrality.

We furthermore compute the maximum library abundance with Eq. (S118), based on the precise trait variation in our data set. The mutant library has an effective lag time *λ*_lib_ = 1.95 h (compared to wild-type lag time *λ*_wt_ = 1.92 h) and an effective *e*-fold growth time *τ*_lib_ = 2.57 h (compared to the wild-type *τ*_wt_ = 2.46 h), leading to an overall negative relative fitness 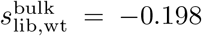. based on a LFC value of LFC^bulk^ = log 100 (a typical fold-change in our data set) and the limit of low relative abundance of the library. We insert these trait values into Eq. (S118) and obtain a maximum mutant library abundance 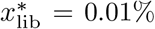. Figure 5D in the main text shows that this much smaller mutant library abundance is able to keep the relative error below the desired threshold for all mutant genotypes. Note, however, this is still not fully exact since we have ignored the underlying variation in biomass yield in our trait data set.

---

**FIG. S1.**
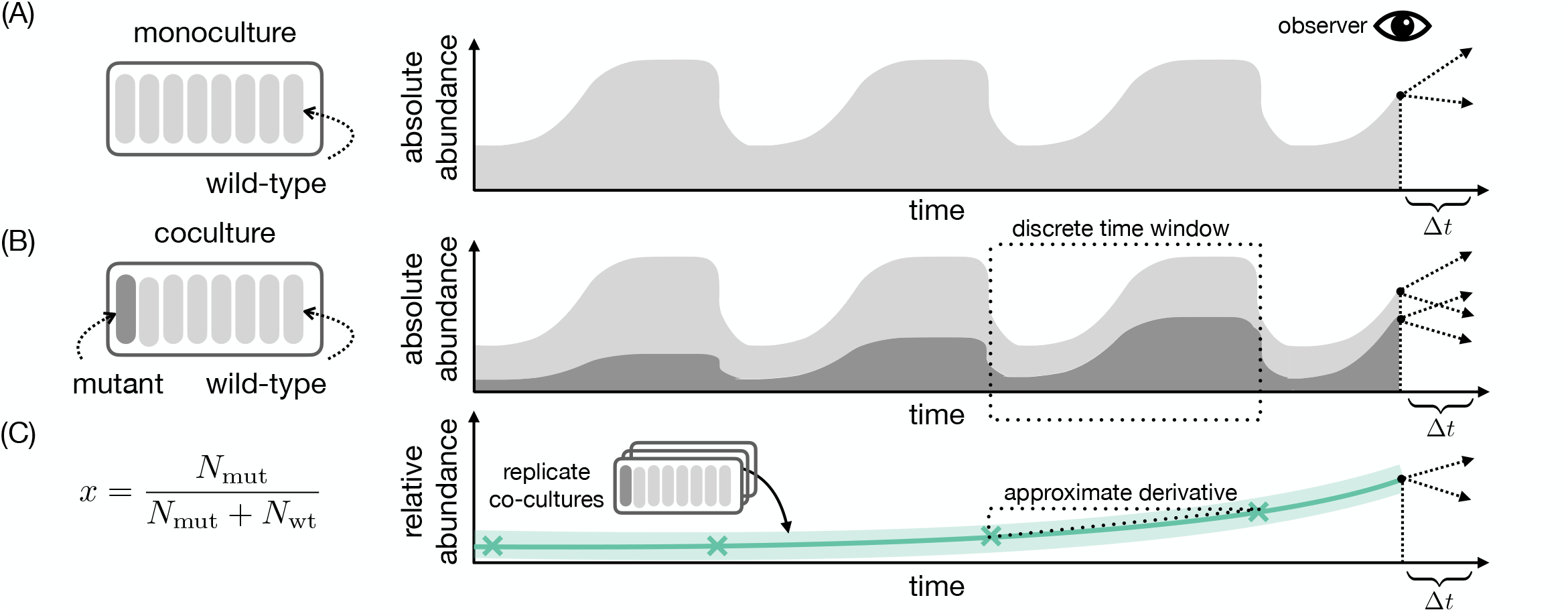
Predicting the absolute and relative abundance of microbial populations. (A) Example time series of absolute abundance for a single microbial population (light grey). The observer (eye symbol) of the time series at time *t* can ask about the future trend in absolute abundance (arrows). (B) Same as panel (A) but for two microbial strains, such as a wild-type and a mutant. The absolute abundance of the wild-type strain (light grey) is stacked on top of the absolute abundance of the mutant strain (dark grey). (C) The relative abundance for the mutant strain corresponding to panel (B). We sketch the mean relative abundance *x* over time (green line) inferred from multiple replicate cocultures. An error band (light green) around the time series demonstrates the variation between replicates.

**FIG. S2.**
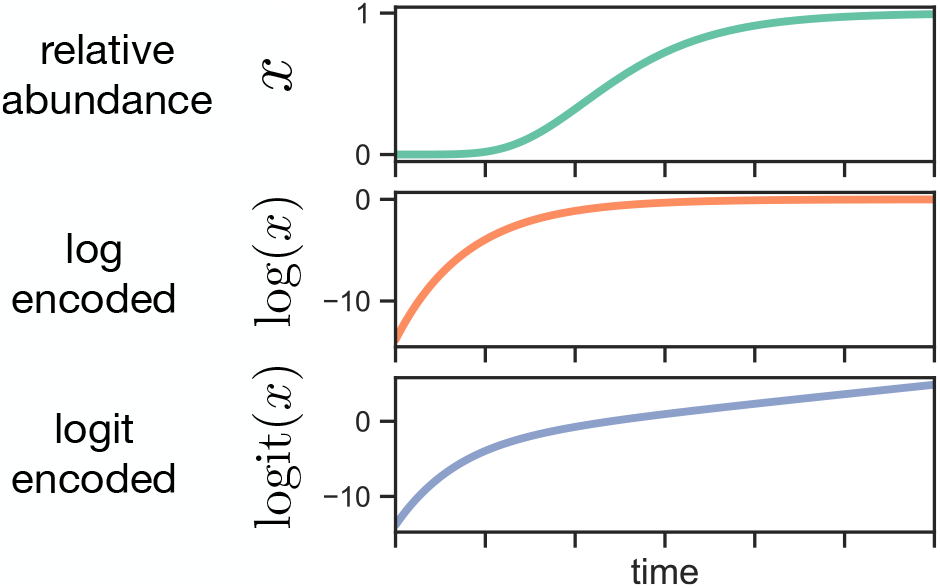
The effect of encodings on a non-logistic relative abundance trajectory. Example trajectory of relative abundance *x* (top panel) for a mutant invading and eventually replacing a wild-type population, simulated with the Gompertz equation *dx/dt* = *rx* log(1*/x*). Below, we show the Gompertz trajectory under the encodings log *x* and logit *x* = log(*x/*(1 − *x*)). Compare to Fig. 1A for a trajectory simulated with the logistic equation.

**FIG. S3.**
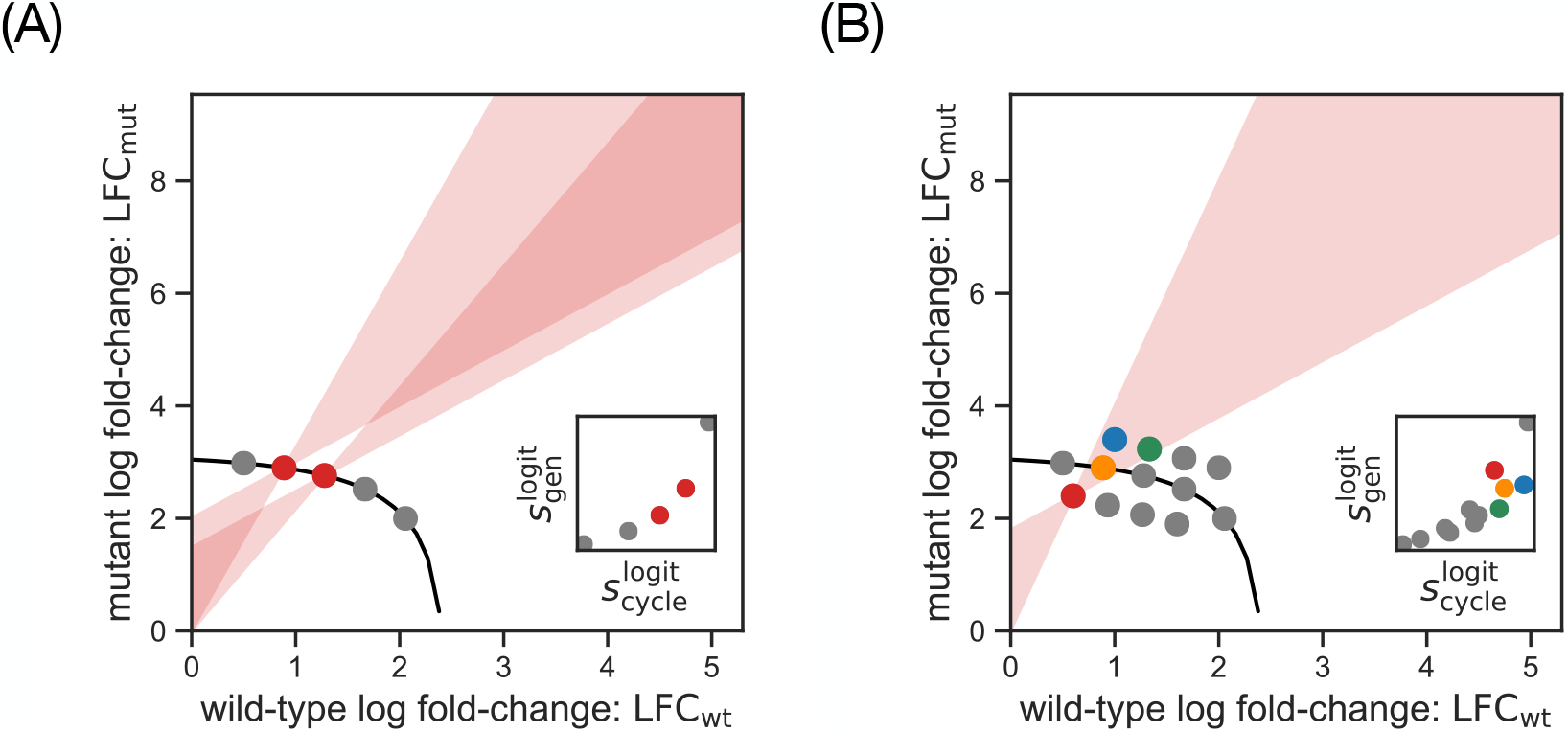
The variation of wild-type and mutant log fold-change under resource consumption constraints. (A) Schematic fold-change variation under a perfect resource consumption constraint between wild-type and mutant. Each dot corresponds to a mutant strain in a 1:1 competition growth cycle with the wild-type strain (Sec. S5). There is negative covariation between mutant and wild-type LFCs because the resource constraint forces one to decrease if the other increases. For our model of population dynamics, we can calculate this constraint exactly (black line: Eq. (S36) in Sec. S5). For two mutants (red points) we highlight the bow tie-shaped areas (red shading; compare Fig. 1D) where other competitions must lie to have a rank discrepancy between relative fitness per-cycle 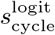 and per-generation 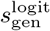 with those mutants, but they are inevitably empty due to the negative covariance across competitions from the resource constraint. Inset: Correlation of fitness per-cycle and per-generation for these same competitions, showing no rank discrepancy. (B) Same as (A) but schematically showing how a deviation from the resource consumption constraint can allow competitions to fall in each other’s red bow tie-shaped regions (e.g., blue, orange, and green competitions compared to the red one), and thus have a rank discrepancy between relative fitness per-cycle 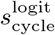 and per-generation 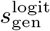 (inset).

**FIG. S4.**
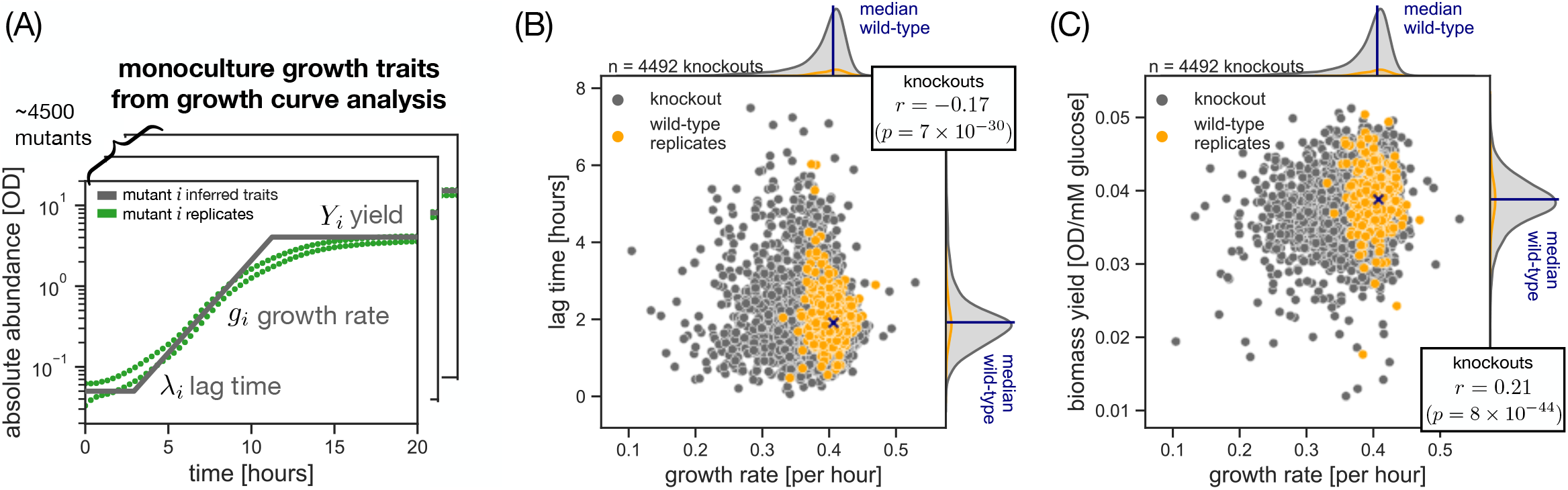
Empirical trait distribution for single gene-knockouts of yeast. (A) Overview of the growth curve data set and the estimated growth traits for the knockout library of *Saccharomyces cerevisiae* (Methods) [6]. (B) Covariation between estimated maximum growth rate *g* and lag time *λ* across all mutant strains (grey dots; Pearson correlation coefficient *r* = −0.17, *p* = 7 × 10^−30^) as well as wild-type replicates (orange dots; *r* = −0.16, *p* = 0.002). The reference wild-type strain for our pairwise coculture simulations is defined by the median trait values (black cross) of all wild-type replicates. (C) Covariation between measured maximum growth rate *g* and biomass yield *Y* across all mutant strains (grey dots; *r* = 0.21, *p* = 8 × 10^−44^) as well as wild-type replicates (orange dots; *r* = −0.06, *p* = 0.25). Histograms on the top and right sides of (B) and (C) are marginal distributions along each axis, with colors matching the scatter points and the vertical black line marking the median wild-type trait value.

**FIG. S5.**
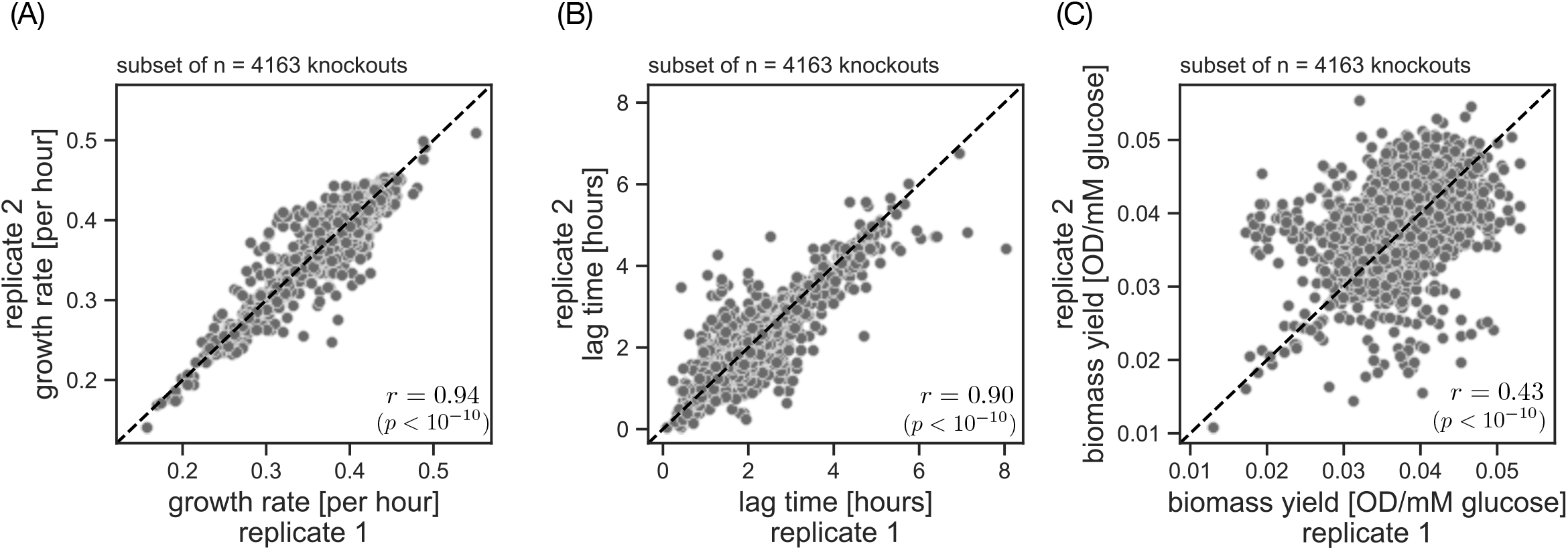
Replicate measurements for growth rate, lag time, and yield in our empirical trait data set. (A) Co-variation of growth rate between replicate measurements of the knockouts (grey dots; Pearson correlation coefficient *r* = 0.94, *p* = 0.00). Each dot represents a mutant genotype from the single-gene knockout collection in *Saccharomyces cerevisiae* [6]. For the vast majority of genotypes in our data set (4163 out of 4492 knockouts) we are able to fit two traits from two independent growth curve measurements (Fig. S4A; Methods). (B) Covariation of lag time between replicate measurements of the knockouts (grey dots; *r* = 0.90, *p* = 0.00) (C) Covariation of biomass yield between replicate measurements of the knockouts (grey dots; *r* = 0.43, *p* = 4.81 × 10^−188^).

**FIG. S6.**
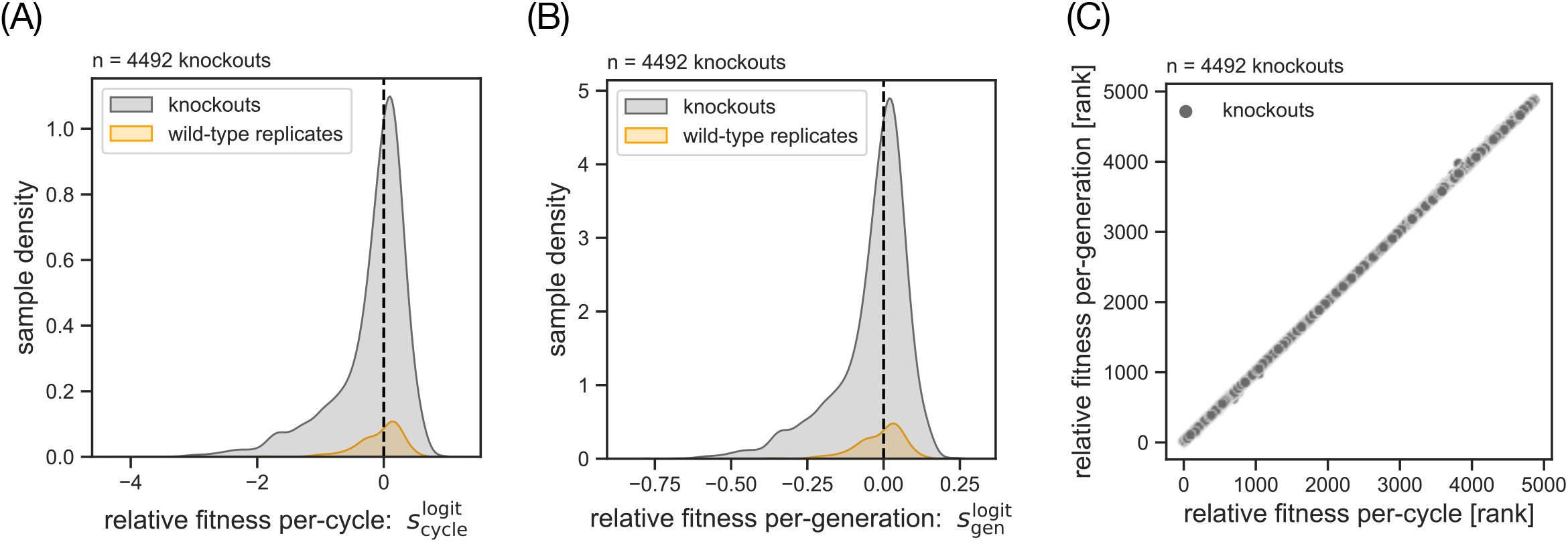
The distribution of fitness effects under relative fitness per-cycle and per-generation. (A) Distribution of relative fitness per-cycle 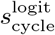 for the single-gene knockouts of yeast (grey) and wild-type replicates (orange) in our data set (Fig. S4A, Methods) [6], based on the fitness data from Fig. 2B. The vertical black line marks zero relative fitness (e.g., the median wild-type). (B) Same as (A) but with relative fitness per-generation 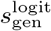. (C) Covariation in fitness ranks between relative fitness per-cycle 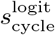 and per-generation 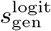. For each fitness statistic, we calculate the mutant ranking (higher rank means higher fitness and mutants with equal fitness are assigned the lowest rank in the group), based on the rank data from Fig. 2B.

**FIG. S7.**
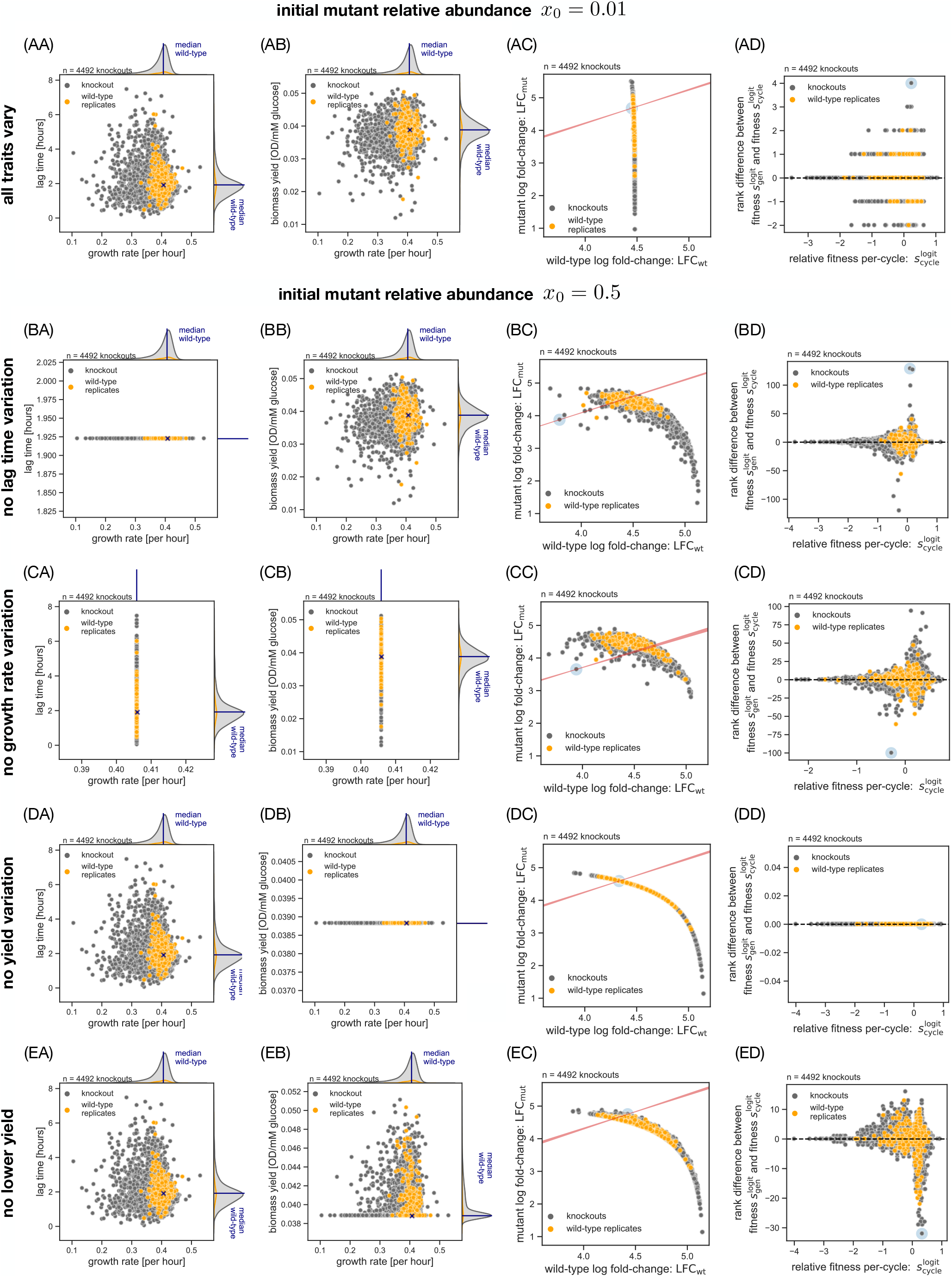
Exploring alternative conditions for rank discrepancy between fitness statistics in yeast gene-deletion data. Plots columns (A) and (B) have the same format as Fig. S4B,C, and columns (C) and (D) match Fig. 2B,C (in reverse order). Row (A): Low mutant frequency *x* = 0.01. Row (B): Standard mutant frequency *x* = 0.5, but where we artificially eliminate variation in lag time to test the effect on the fitness rank discrepancy. Row (C): Same as row (B) but no variation in growth rate. Row (D): Same as row (B) but no variation in biomass yield. Row (E): Same as row (B), but yields of mutants lower than the median wild-type yield are set to the wild-type value.

**FIG. S8.**
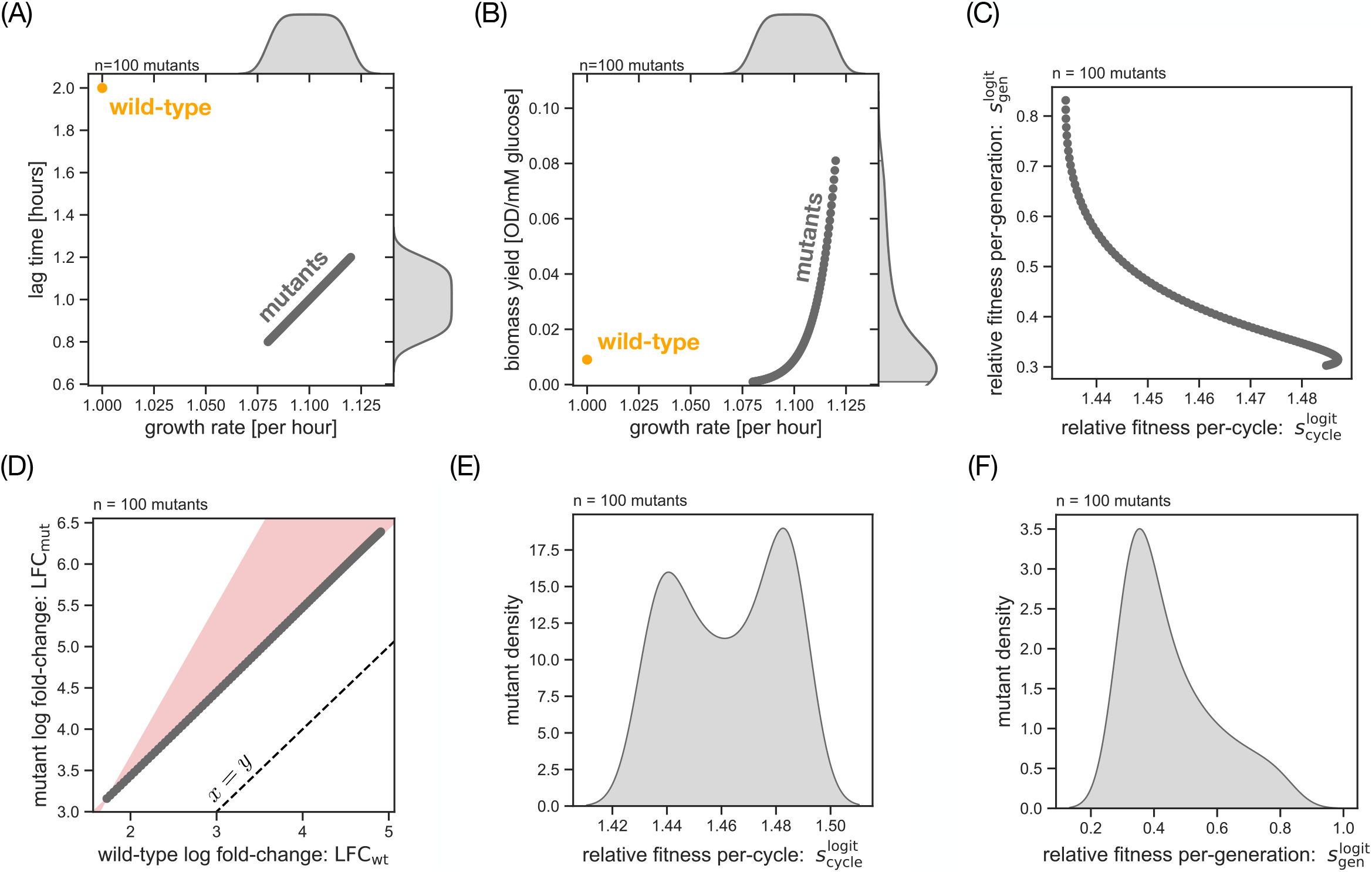
Hypothetical case where relative fitness per-cycle and per-generation are anticorrelated across mutants. (A) Covariation between growth rate and lag time for a synthetically generated set of mutants (grey dots) and a single wild-type (orange dots). The variation here may represent the standing variation in an evolved population with improved growth rate and lag time over the wild-type ancestor. The graphic overlap between points means they appear as a single line. (B) Covariation between growth rate and biomass yield for the mutants (grey dots) and wild-type (orange dot) in this synthetic data set. (C) Covariation between relative fitness per-cycle 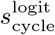 and per-generation 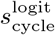. We simulate pairwise competitions for all mutants against the wild-type with the exact same settings as for the empirical data set (Fig. 2A; Methods). (D) Covariation between mutant and wild-type fold-change in the competitions. Based on the competition data in panel C (grey dots), we estimate the LFC values for all mutant-to-wild-type competitions (grey dots). We highlight the bow tie-shaped area of rank discrepancy for the mutant with the lowest overall fold-change (red shading; compare Fig. 1D). The dashed black line shows the isocline for zero relative fitness per-cycle (Eq. (9)). Distributions of relative fitness per-cycle 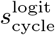 (E) and per-generation 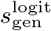 (F) for the mutants in panel (A,B). Based on the fitness data from panel (C).

**FIG. S9.**
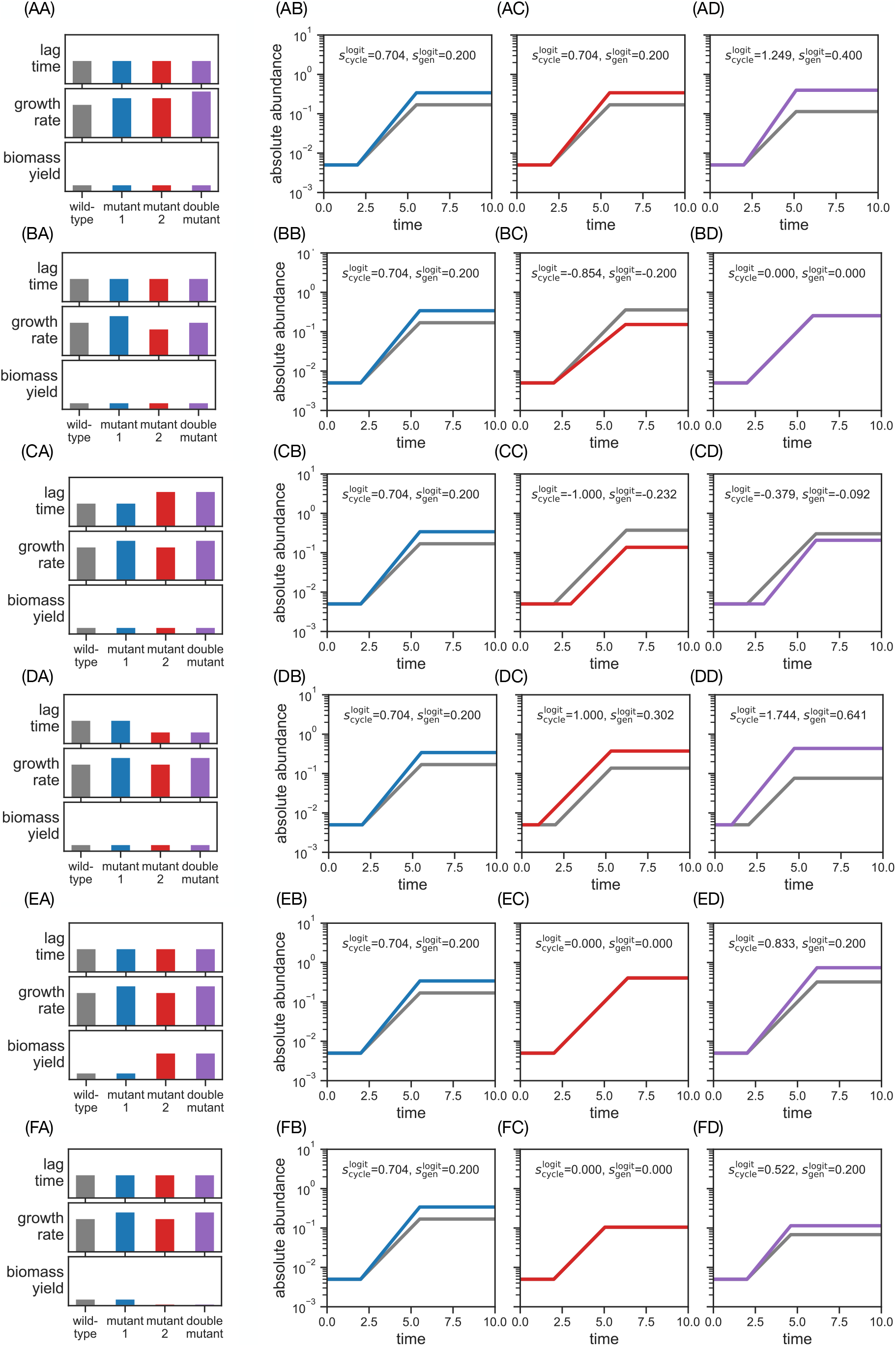
Competition growth cycles for double mutants — first column of Fig. 3C,D. Each row corresponds to a pair of mutations that perturb growth traits in the minimal model of population dynamics (Fig. 2A, Methods). This figure shows all combinations where the first mutant increases growth rate (first column of Fig. 3C,D). Column (A): Perturbations of growth traits for single and double mutants. Column (B): Growth curves of wild-type (grey) and mutant 1 (blue). Column (C): Growth curves of wild-type (grey) and mutant 2 (red). Column (D): Growth curves of wild-type (grey) and double mutant (purple).

**FIG. S10.**
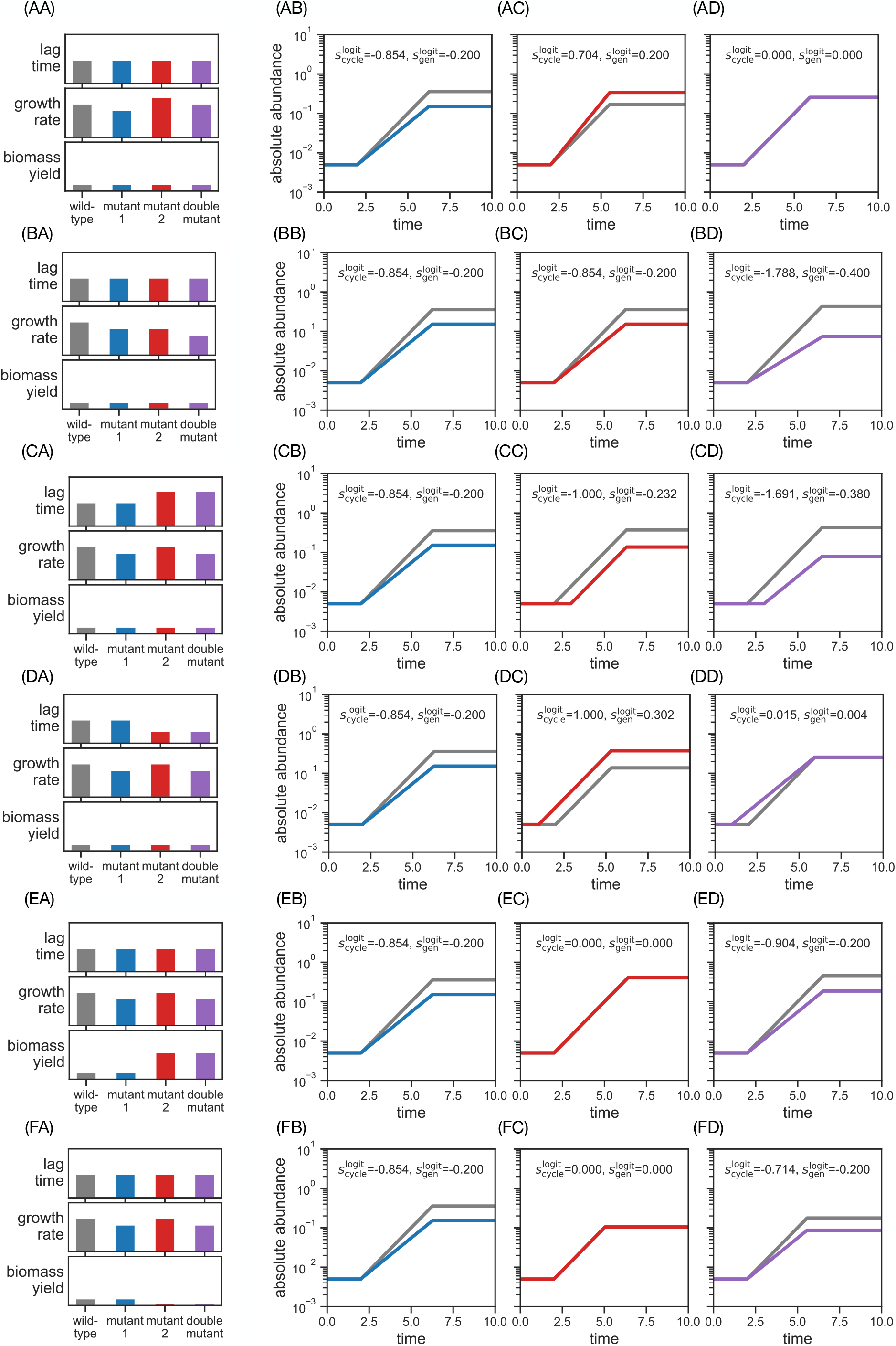
Competition growth cycles for double mutants — second column of Fig. 3C,D. Same as Fig.S9 but for mutant pairs where the first mutant decreases growth rate.

**FIG. S11.**
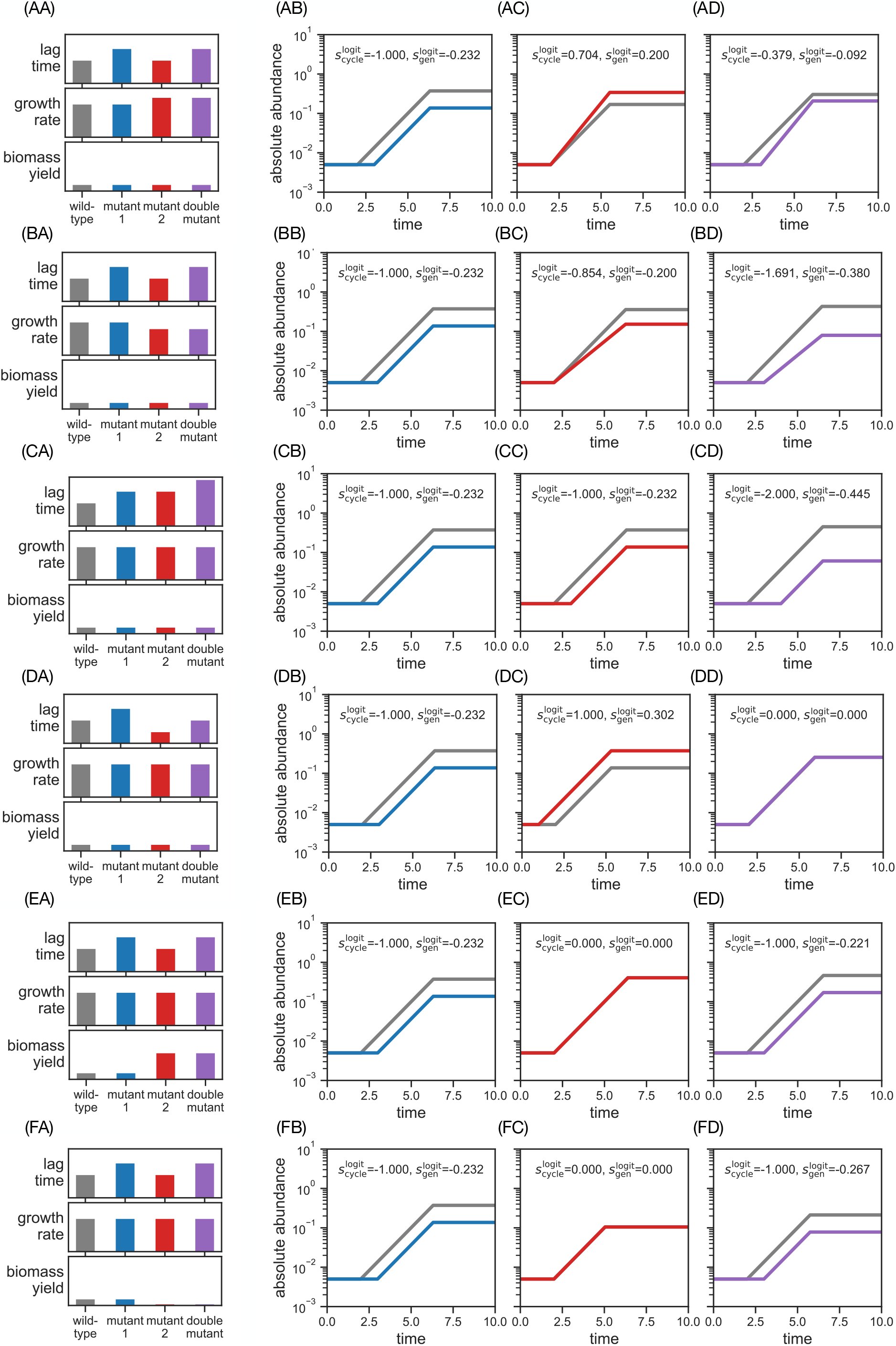
Competition growth cycles for double mutants — third column of Fig. 3C,D. Same as Fig.S9 but for mutant pairs where the first mutant increases lag time.

**FIG. S12.**
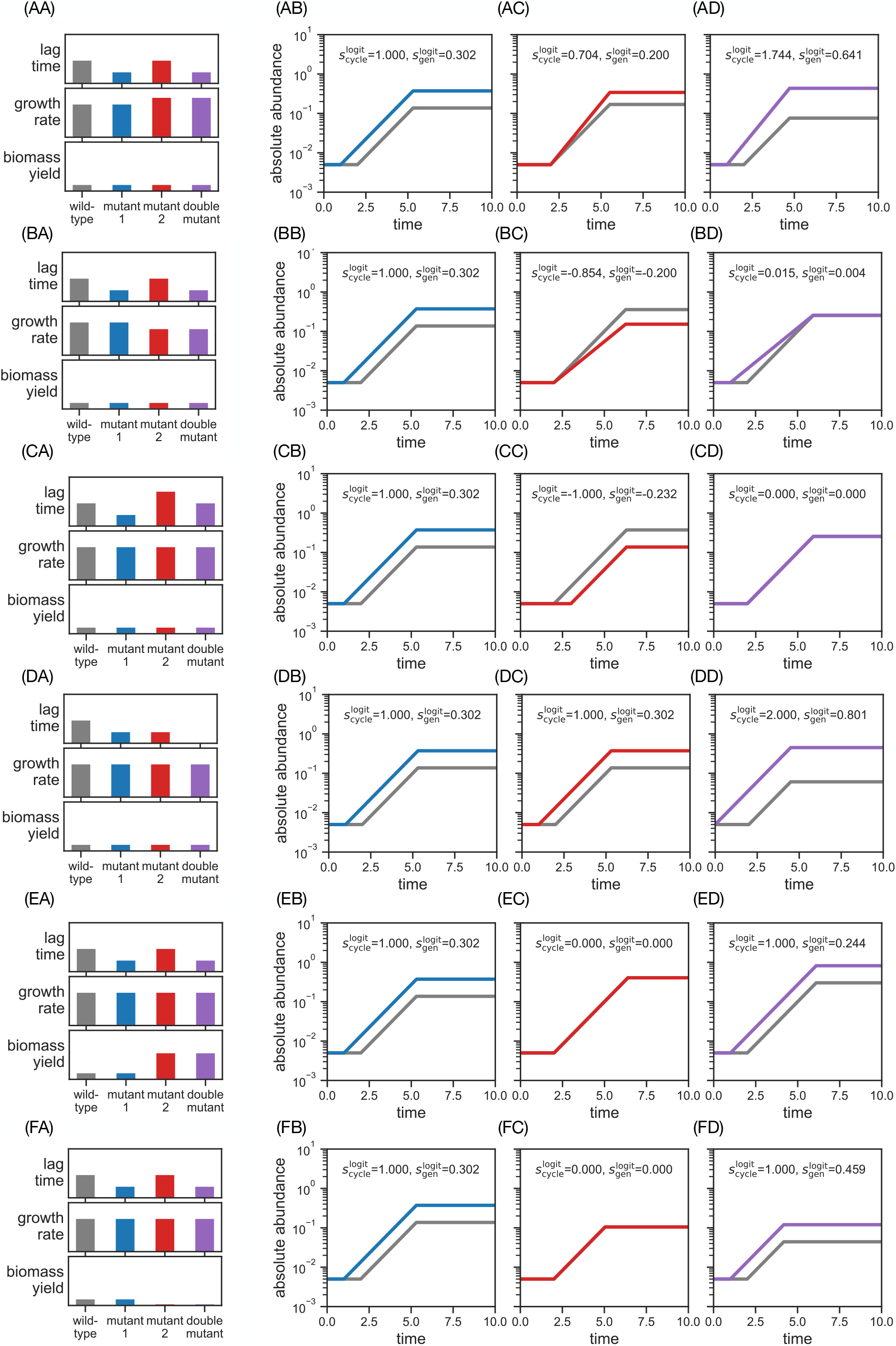
Competition growth cycles for double mutants — fourth column of Fig. 3C,D. Same as Fig. S9 but for mutant pairs where the first mutant decreases lag time.

**FIG. S13.**
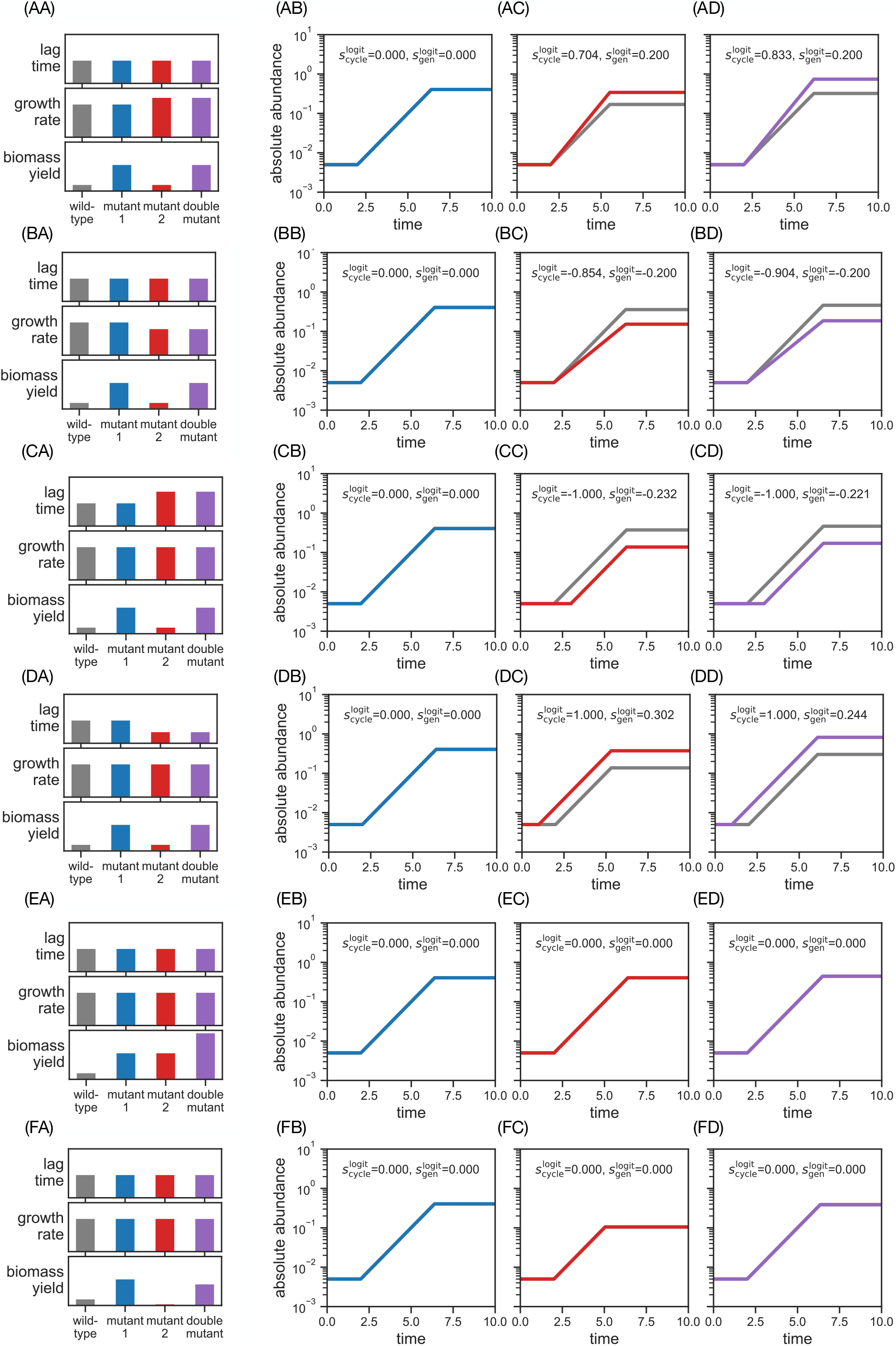
Competition growth cycles for double mutants — fifth column of Fig. 3C,D. Same as Fig. S9 but for mutant pairs where the first mutant increases biomass yield.

**FIG. S14.**
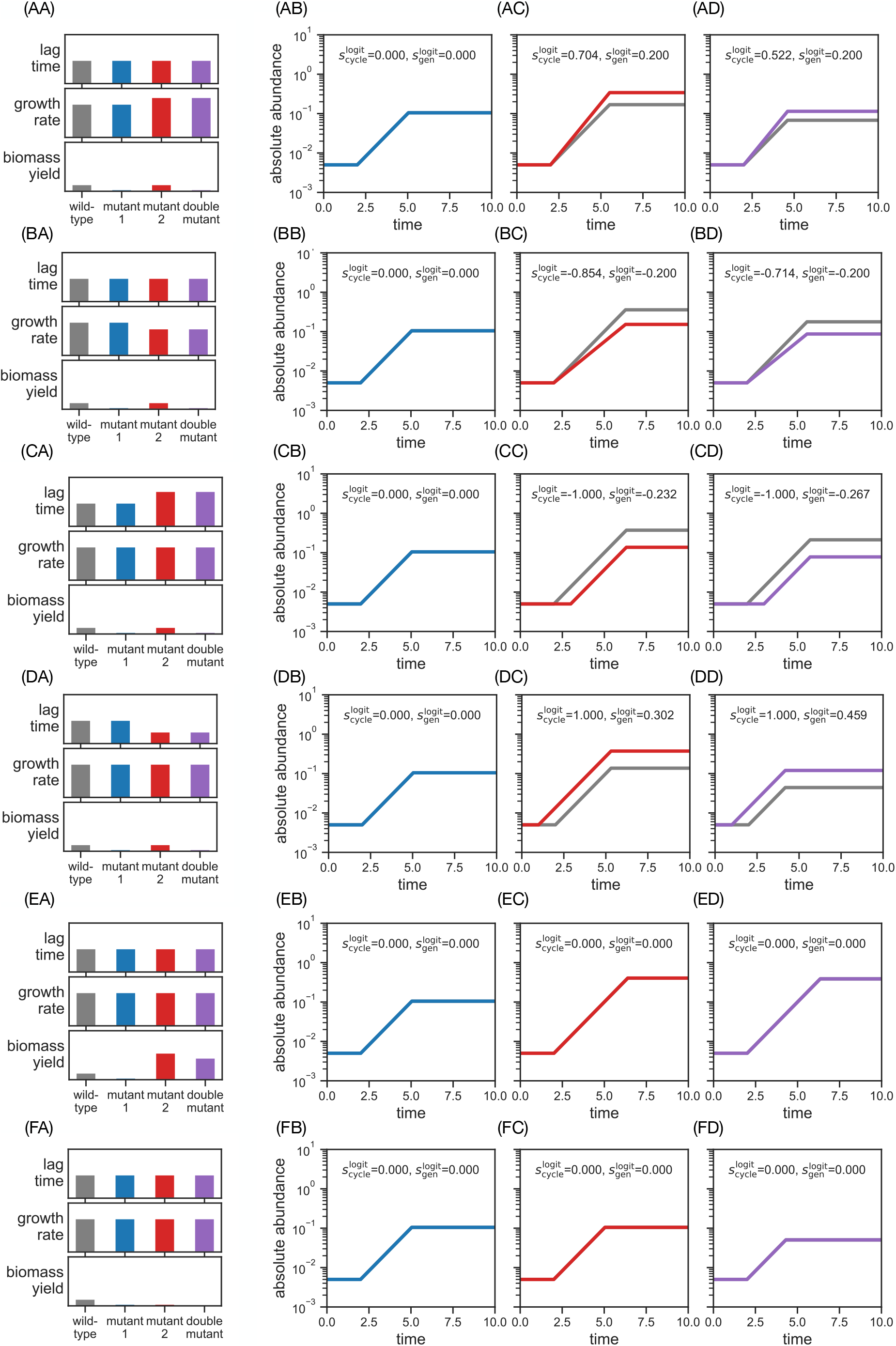
Competition growth cycles for double mutants — sixth column of Fig. 3C,D. Same as Fig. but for mutant pairs where the first mutant decreases biomass yield. For the double mutant where the second mutant also decreases yield, we cannot use the additive mutation effects (Methods) because that would result in a negative yield for the double mutant. Here we have defined the double mutant to have a yeield value of *Y*_mut_ = 0.1. This choice does not affect the fitness or epistasis, since biomass yield is a neutral trait with fitness zero in our model of population dynamics.

**FIG. S15.**
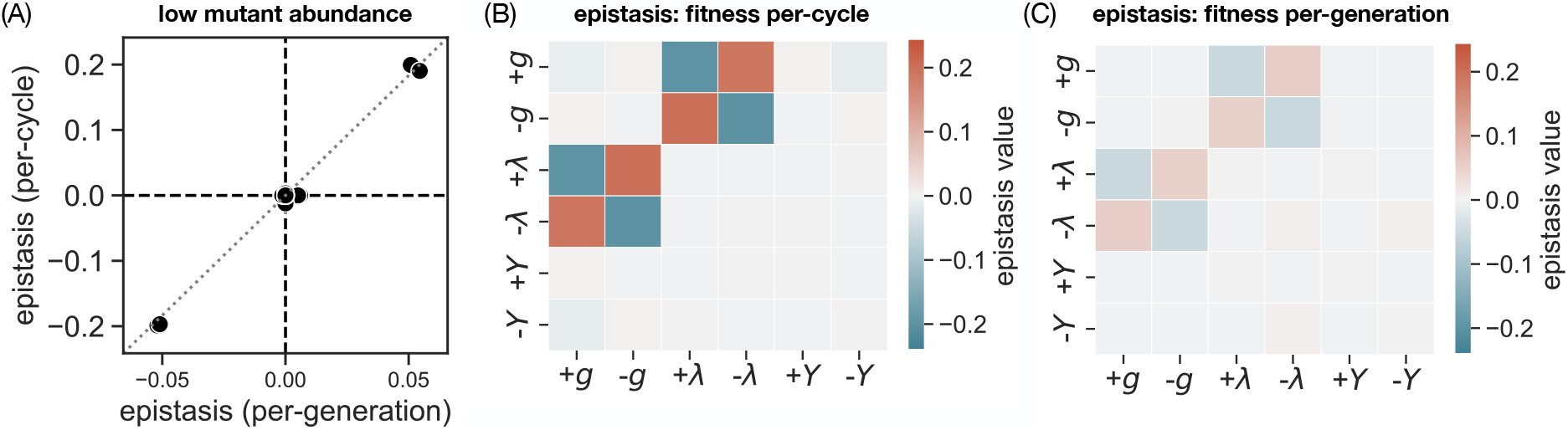
Epistasis patterns at low mutant relative abundance. Same as Fig. 3B–D but where the mutant starts each competition at low relative abundance (*x*_0_ = 0.01; Sec. S8), compared to the 1:1 initial conditions in Fig. (*x*_0_ = 0.5).

**FIG. S16.**
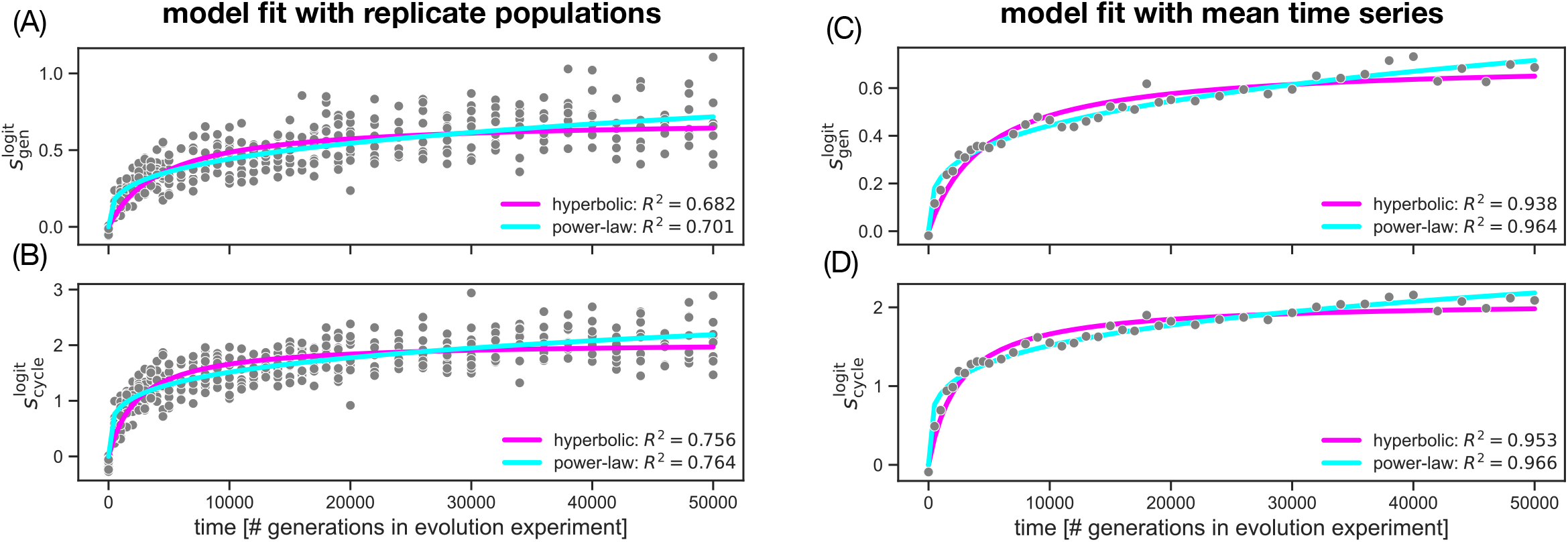
Long-term fitness trends in the LTEE under relative fitness per-cycle and per-generation. (A) Fit of a hyperbolic model (pink line; Eq. (S56) in Sec. S9) and a power-law model (cyan line; Eq. (S57) in Sec. S9) to a pooled time-series of relative fitness per-generation 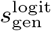. We pool the fitness per-generation 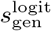 from all 12 lines of the LTEE into a single data set (grey dots) and repeat the fits of Wiser et al. [11] (Sec. S9). We compute the fraction of variance explained *R*^2^ as a measure for the quality of fit. (B) Same as panel (A) but using relative fitness per-cycle 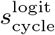. (C) Same as panel (A) but averaging fitness across all 12 lines at each time point. Note that the fraction of variance explained in is much higher compared to panel (A), because the averaged time-series has fewer points (Sec. S9). (D) Same as panel (C) but using relative fitness per-cycle 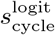. For a comparison between our results and the original fit by Wiser et al. [11], see Sec. S9.

**FIG. S17.**
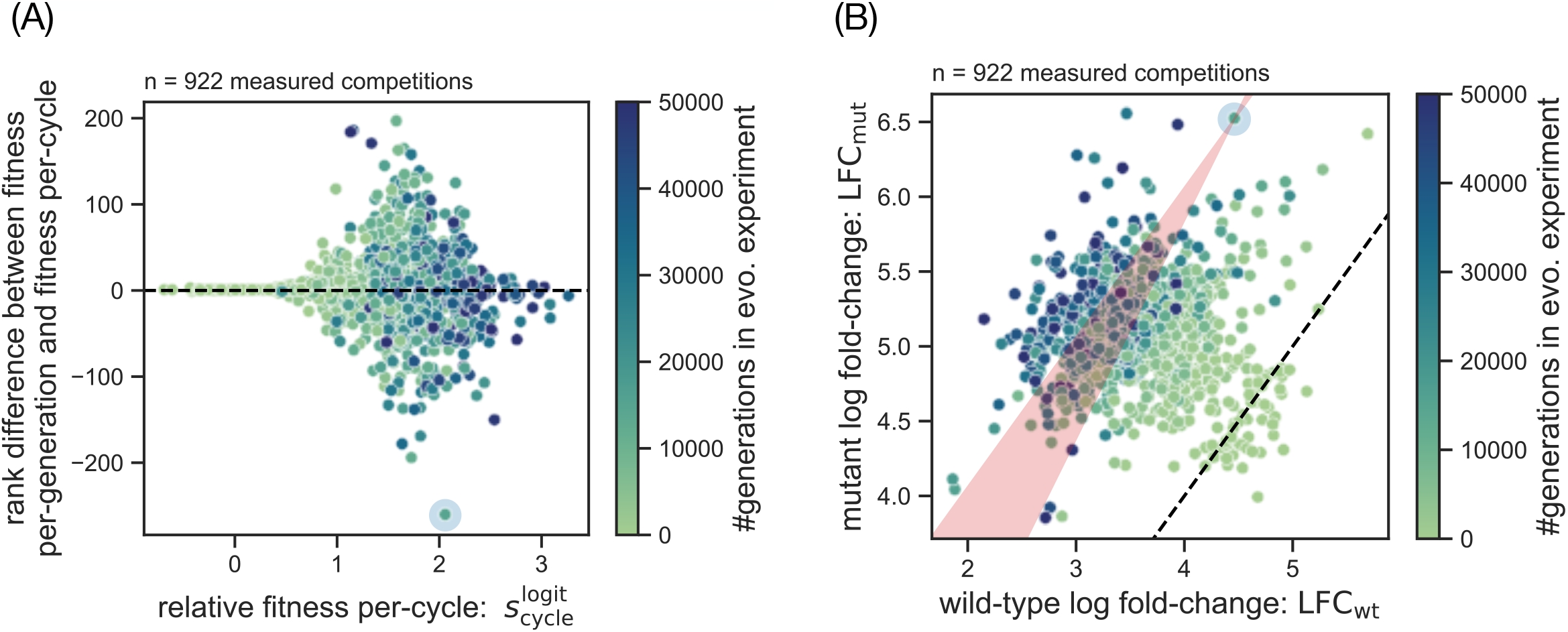
Comparison of relative fitness per-generation and per-cycle across the complete LTEE competition data set. (A) Rank discrepancy between relative fitness per-cycle 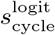 and per-generation 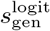 for evolved populations of the LTEE. For each fitness statistic, we calculate a ranking (higher rank means higher fitness and mutants with equal fitness are assigned the lowest rank in the group; compare Fig. 2B) across all 12 evolved lines and all time points. The rank difference is defined as the rank in 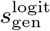 minus the rank in 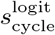. We highlight the evolved population with the greatest rank difference (blue halo). (B) Covariation between the wild-type and mutant fold-change for the competition data from the LTEE [11]. The term “mutant” (y-axis) refers to an evolved population at a given time point from one of the 12 lines of the LTEE. We highlight the evolved population with the greatest rank difference (blue halo; same as in panel (A)) and draw its corresponding bow tie-shaped area of rank discrepancy between 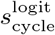 and 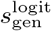 (red shading; compare Fig. 1D). The coloring in all panels refers to the time point at which the evolved population was sampled from the LTEE.

**FIG. S18.**
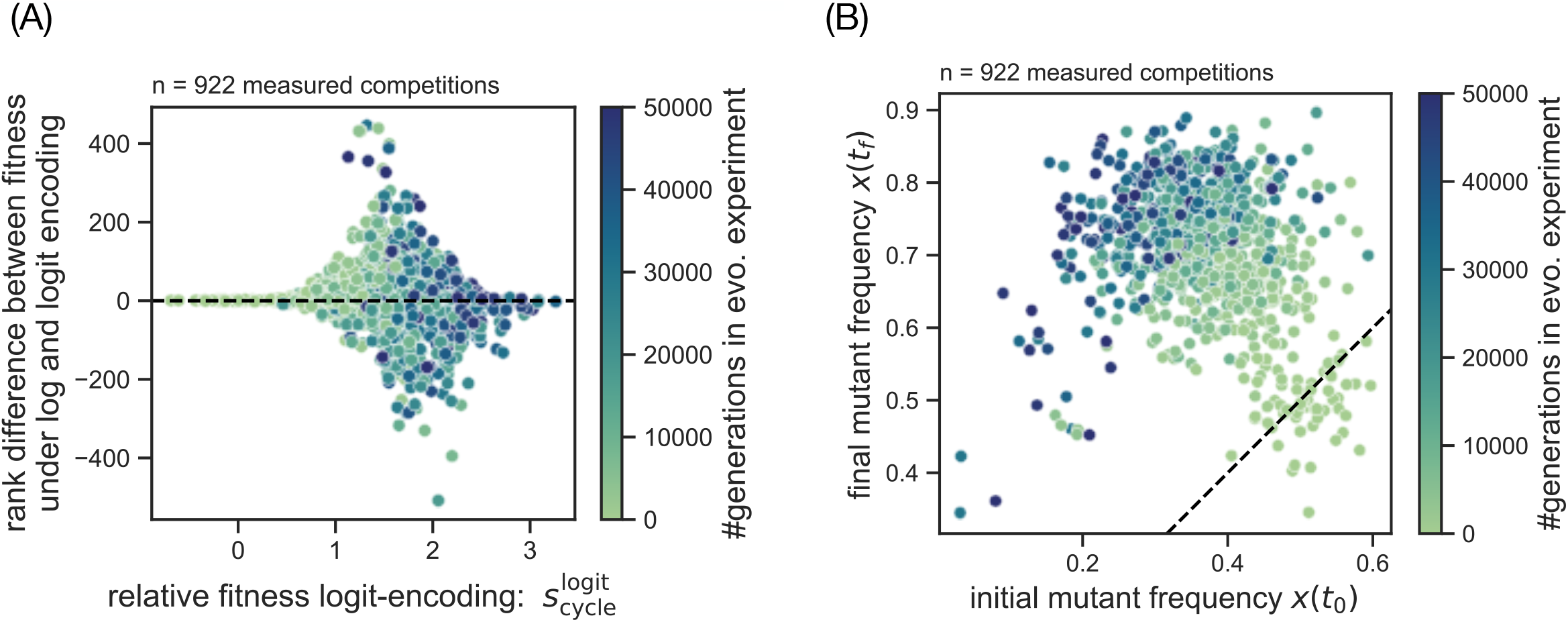
Comparison of log-encoding and logit-encoding for relative fitness per-cycle in the LTEE competition data set. (A) Same as Fig. S17A but for rank discrepancy between logit-encoded relative fitness per-cycle 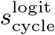 and log-encoded relative fitness per-cycle 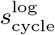 for evolved populations of the LTEE. (B) Covariation between the wild-type relative abundance and mutant relative abundance at the start of the competition growth cycle for the competition data from the LTEE [11]. The term “mutant” in the axis labels refers to an evolved population at a given time point from one of the 12 lines of the LTEE. The coloring in all panels refers to the time point at which the evolved population was sampled from the LTEE.

**FIG. S19.**
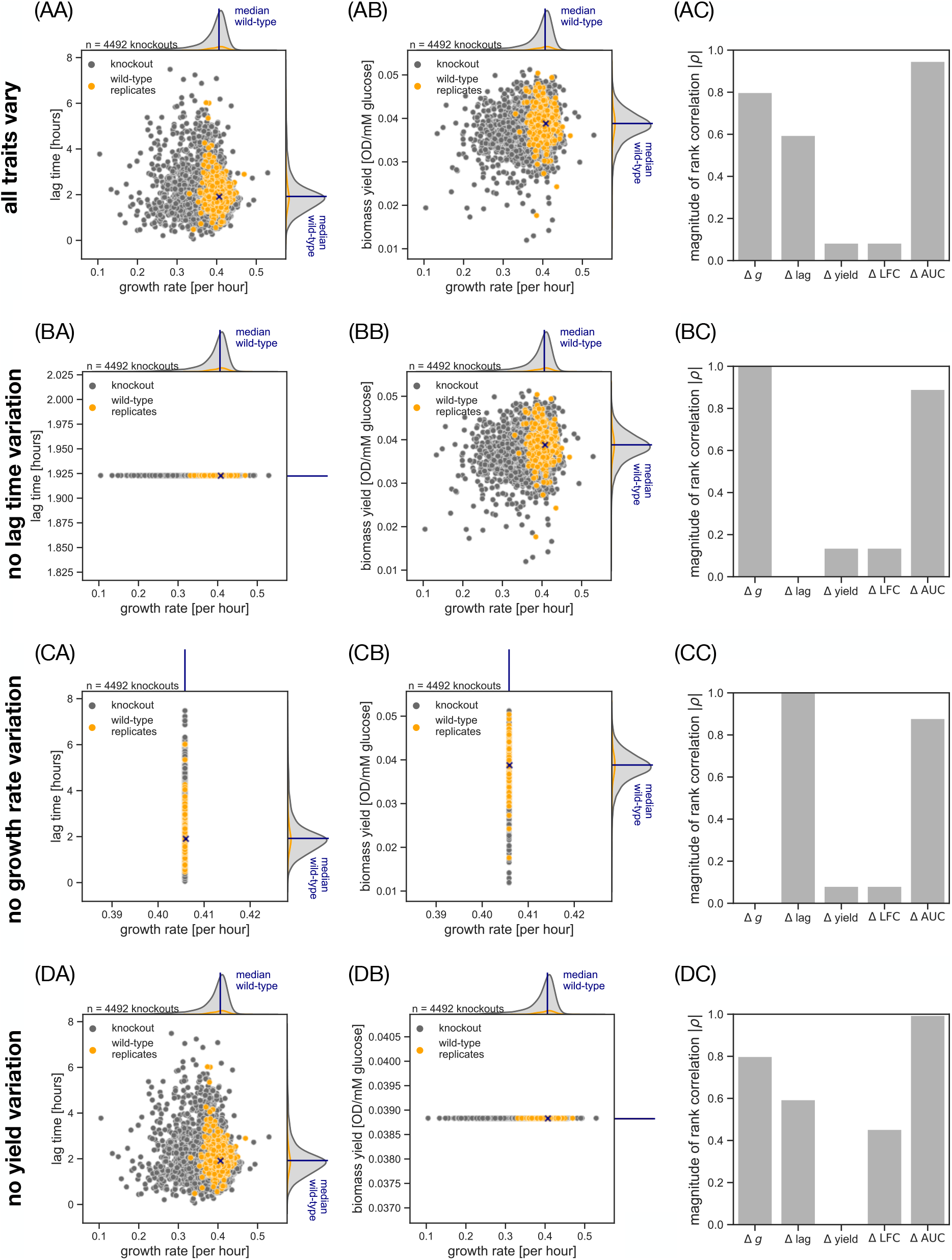
Predicting relative fitness with monoculture proxies under different scenarios of trait variation. Plots in columns (A) and (B) show growth trait variation across mutants, with the same format as Fig. S4B,C. Plots in column (C) show quality of prediction for different monoculture proxies under the trait distribution in columns (A) and (B). As the ground truth, we estimate the relative fitness per-cycle 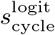 using a simulated 1:1 competition growth cycle (Fig. 2A; Methods). For each mutant *i*, we compute the growth rate difference Δ*g* = *g*_*i*_ − *g*_wt_, the lag time difference Δlag = *λ*_*i*_ − *λ*_wt_, the difference in biomass yield Δyield = *Y*_*i*_ − *Y*_wt_, the difference in log fold-change ΔLFC = LFC_*i*_ − LFC_wt_, and the difference in area under the growth curve ΔAUC = AUC_*i*_ − AUC_wt_ from simulated monoculture growth curves (Sec. S10). We quantify the agreement between the monoculture proxy and the relative fitness per-cycle 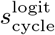 using the Spearman correlation coefficient *τ*. Row (A): Empirical growth trait variation from data on yeast single-gene knockout mutants (identical to Fig. S4B,C). Row (B): Same as row (A) but where we artificially eliminate variation in lag time to test the effect on the monoculture proxies. Row (C): Same as row (A) but with no variation in growth rate. Row (D): Same as row (A) but with no variation in biomass yield.

**FIG. S20.**
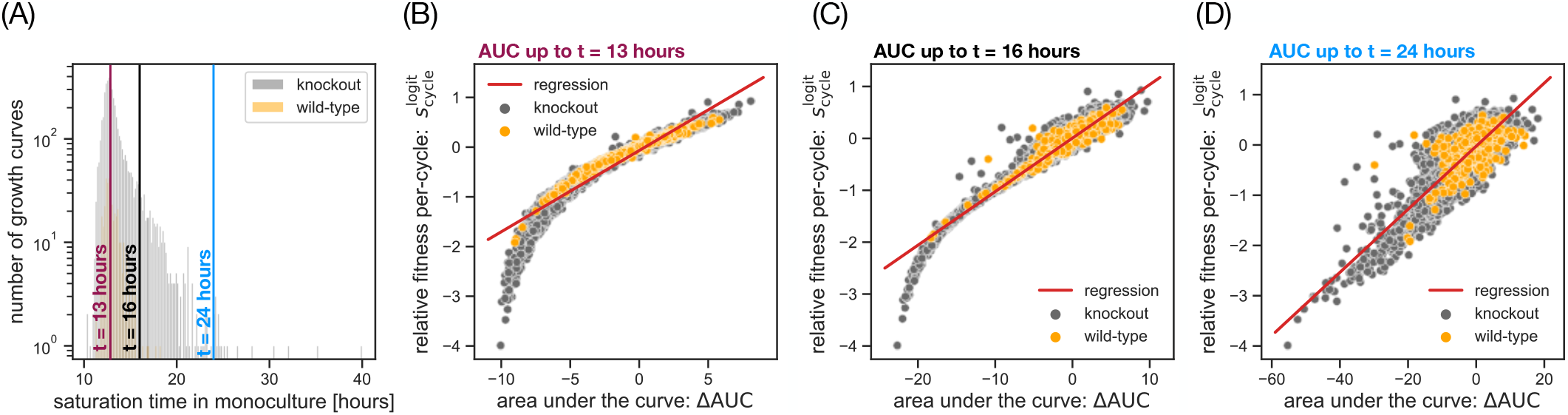
The choice of the cutoff time for evaluating the area under the curve (AUC). (A) Distribution of saturation times in monoculture for the knockouts (grey bars) and wild-type replicates (orange bars) in our empirical data set (Fig. S4). The saturation time *t*_sat_ is defined as the time when the limiting resource is depleted (*R*(*t*_sat_) = 0) and can be estimated numerically from the trait data (Methods). Three vertical lines indicate example choices for the cutoff time *t*_eval_ of the AUC: the most frequent saturation time (*t*_eval_ = 13 hours; red line), a saturation time halfway in the decay of the distribution (*t*_eval_ = 16 hours; black line), and an external time scale (*t*_eval_ = 24 hours; blue line). (B) Covariation between relative fitness per-cycle 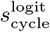 and AUC with the cutoff times marked in panel (A): 13 hours (B), 16 hours (C), and 24 hours (D). We compute the AUC from simulated monoculture knockouts (grey dots) and wild-type replicates (orange dots). As the ground truth, we take the relative fitness per-cycle 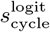 from a pairwise competition (Fig. 2A). The red lines show the best fit from a linear regression of 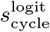 to the AUC.

**FIG. S21.**
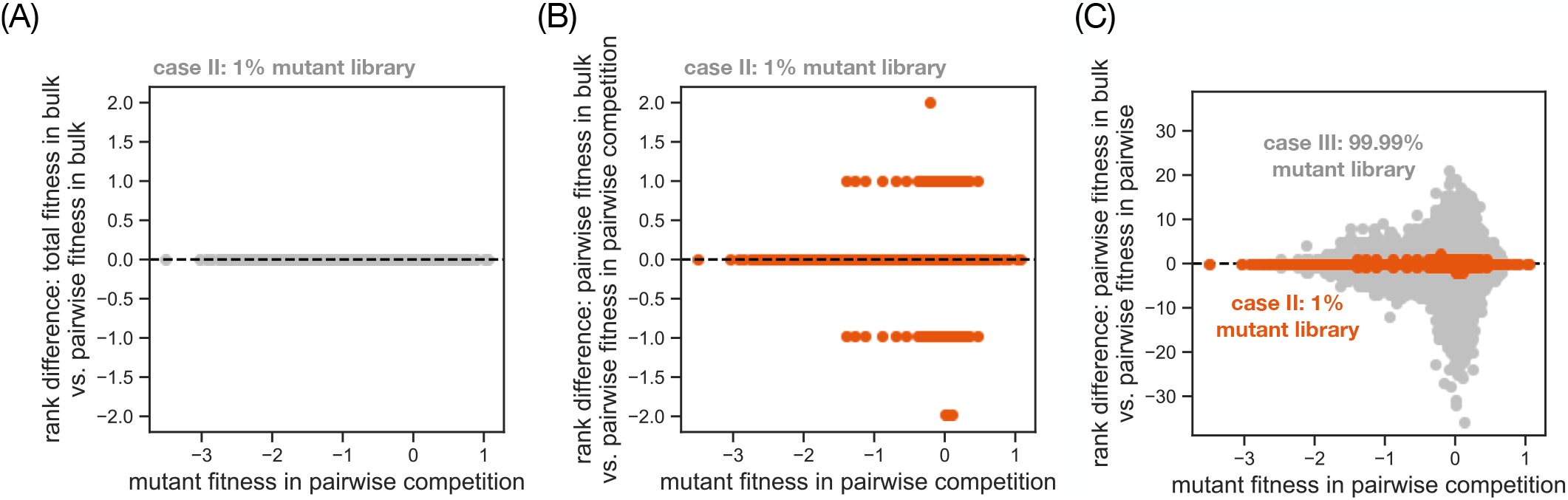
The error in mutant fitness rankings between bulk and pairwise competition experiments. (A) Rank difference between total relative fitness and pairwise relative fitness in a bulk competition growth cycle with low mutant library abundance (Fig. 5A, case II). Based on the fitness data in Fig. 5C, we calculate a mutant ranking for total relative fitness in bulk (Eq. (24)) and a ranking for pairwise relative fitness in bulk (Eq. (25)) (higher rank means higher fitness and mutants with equal fitness are assigned the lowest rank in the group). The rank difference is defined as the rank in total relative fitness 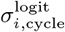 minus the rank in pairwise relative fitness 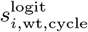. The ranks exactly match in this case because the mutants are sufficiently rate in case II that the population fitness is almost identical to the wild-type fitness. (B) Based on the pairwise relative fitness at low mutant library abundance in Fig. 5C (orange dots; case II), we calcuate a mutant ranking for pairwise relative fitness in bulk (Eq. (25)) and a mutant ranking for the relative fitness per-cycle in pairwise competition (Eq. (9). The rank difference (orange dots) is defined as the rank in bulk competition minus the rank in pairwise competition. The rank differences are small because the mutants are sufficient rare in case II that higher-order effects in the bulk competition are minor. (C) Based on the fitness data in the inset of Fig. 5C, we estimate a mutant ranking for pairwise relative fitness (Eq. (25)) in the case of a bulk competition growth cycle with high mutant library abundance (grey dots; compare Fig. 5A, case III). For each case, the rank difference is defined as the rank in the bulk competition minus the rank in pairwise competition. The rank difference for case II (organge dots) is identical to the data in panel (B). All fitness statistics in this figure are based on the logit-encoding, however, since the underlying relative abundances are small (*x* ≪ 1), it is approximately equivalent to the log-encoding (log *x* ≈ logit *x*).

**FIG. S22.**
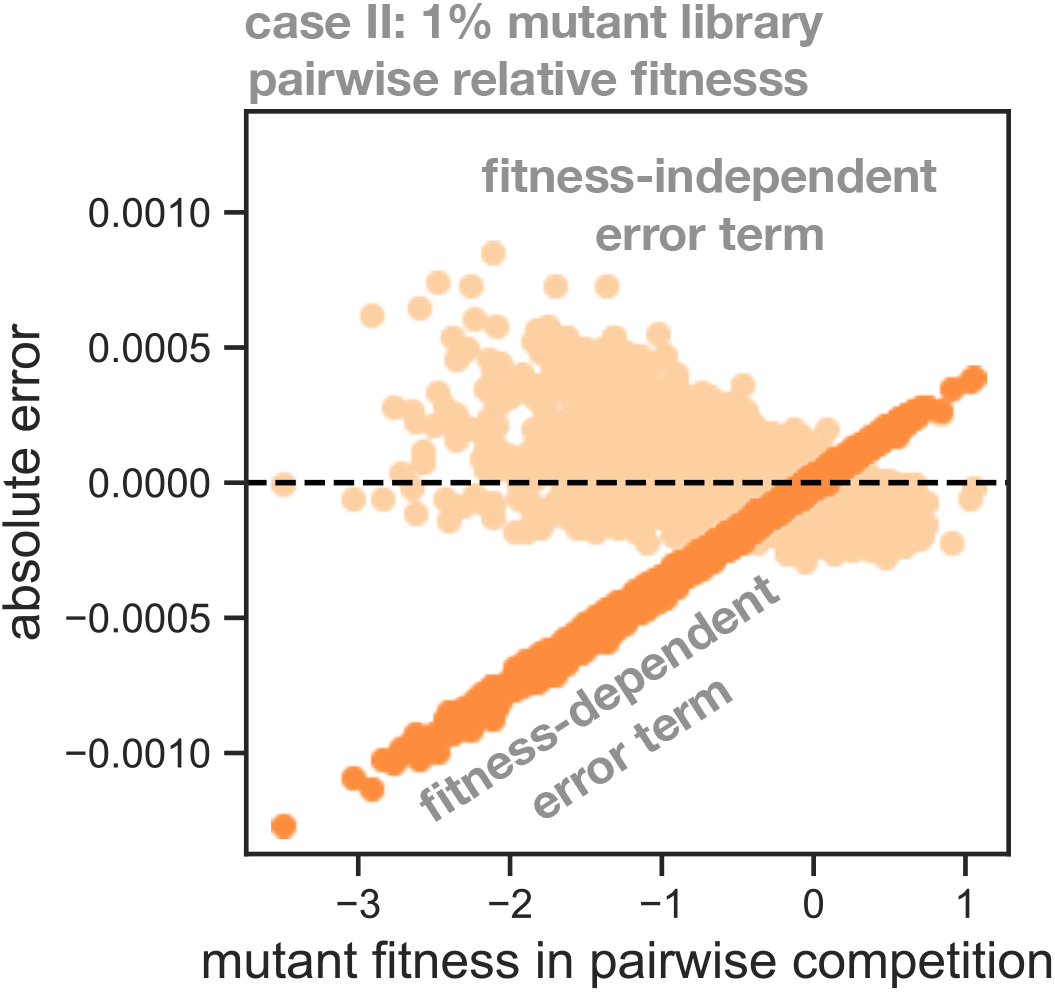
A decomposition for the error from higher-order interactions in bulk competition experiments. For a bulk competition growth cycle with low mutant library abundance (Fig. 5A, case II), we calculate the pairwise relative fitness (Eq. (25)) for the knockouts in our empirical data set (Fig. S4A) using a previously established approximation [10] (Sec. S14). The error from higher-order interactions is defined as the pairwise fitness in bulk minus the fitness in pairwise competition. Based on the approximation for our model of population dynamics (Sec. (S14)), we derive a decomposition that separates the absolute error into two terms that we call the *fitness-dependent error term* (dark orange dots) and the *fitness-independent error term* (light orange dots). For full details on the decomposition, see Sec. (S15).

**FIG. S23.**
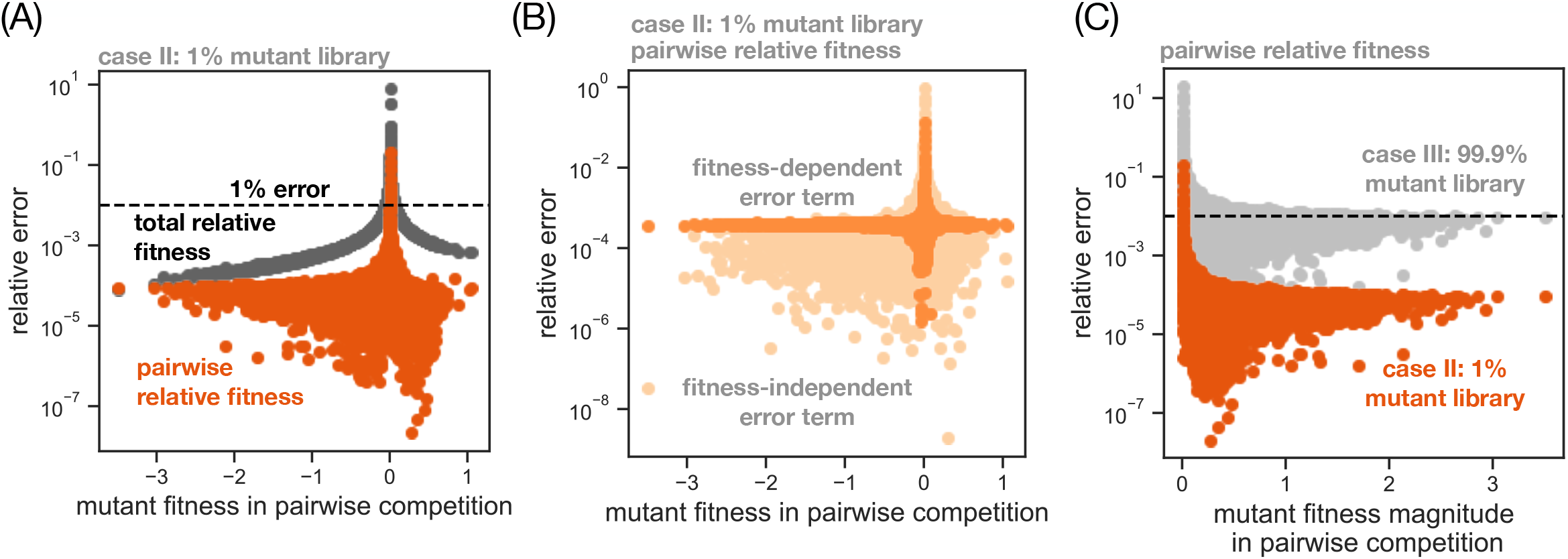
The relative error between bulk and pairwise competition experiments. (A) Same as Fig. 5C but showing relative rather than absolute error. The relative error of each bulk statistic is defined as the absolute error (Fig. 5C), divided by the fitness in pairwise competition (Eq. (S102) in Sec. S16). A dashed grey line indicates the threshold of 1% relative error. (B) Same as Fig. but showing relative rather than absolute error. (C) Same as the inset of Fig. 5C but showing relative rather than absolute error. On the x-axis, we plot the absolute value of relative fitness per-cycle in the pairwise competition.

**FIG. S24.**
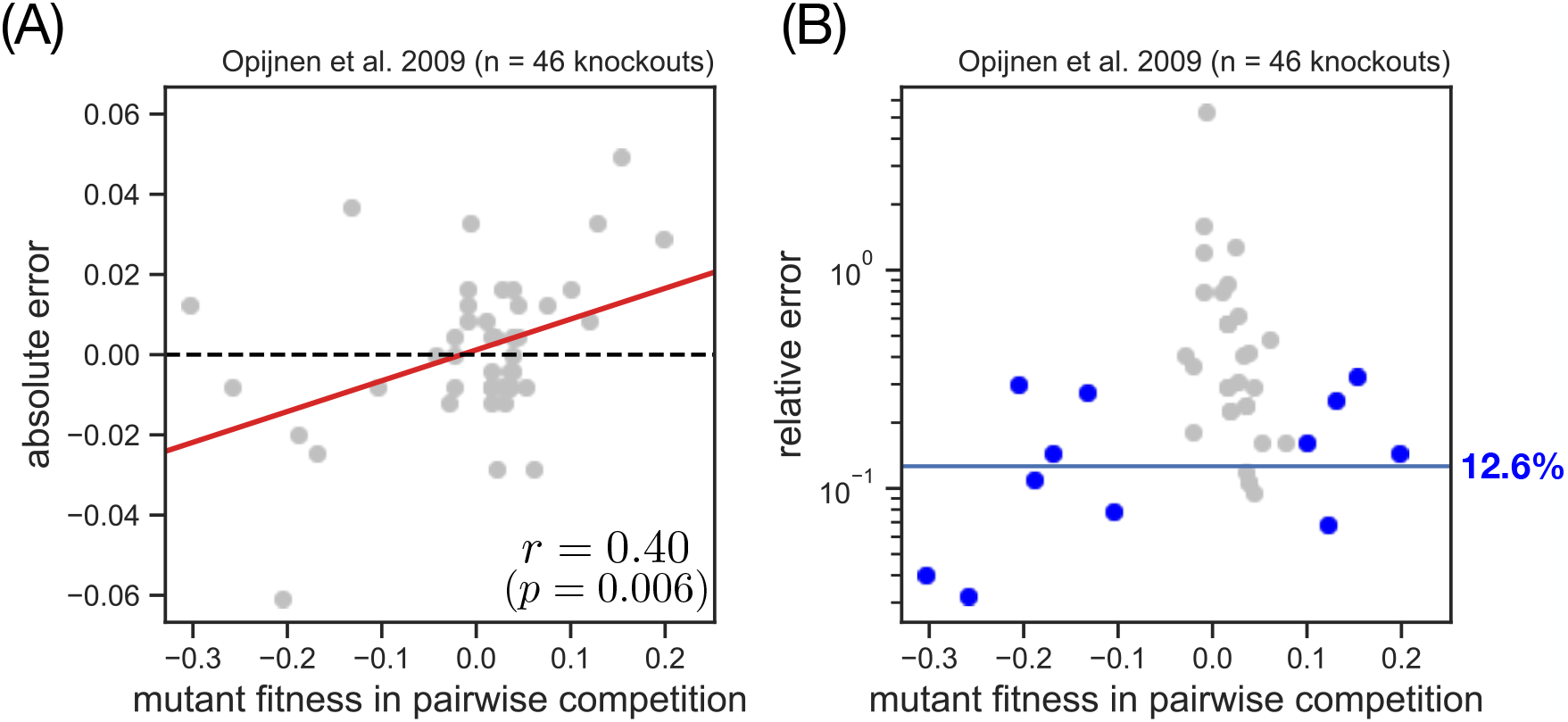
The bulk fitness error based on measurements by van Opijnen et al. [43]. (A) Absolute error 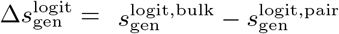 between the pairwise fitness measured in bulk competition 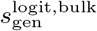 and the pairwise fitness measured in pairwise competition 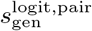 for the set of single-gene knockouts of *S. pneumoniae* shown in Fig. 3c in the original publication. We calculate the Pearson correlation coefficient *r* and the p-value for the null hypothesis of zero correlation, and indicate the linear regression between the two variables with a red line. (B) Same as (A) but showing relative rather than absolute error. We define a set of knockouts with non-neutral fitness (blue dots; 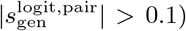 to calculate the geometric mean relative error (blue line). See Fig. 6A for fitness data and experiment conditions.

**FIG. S25.**
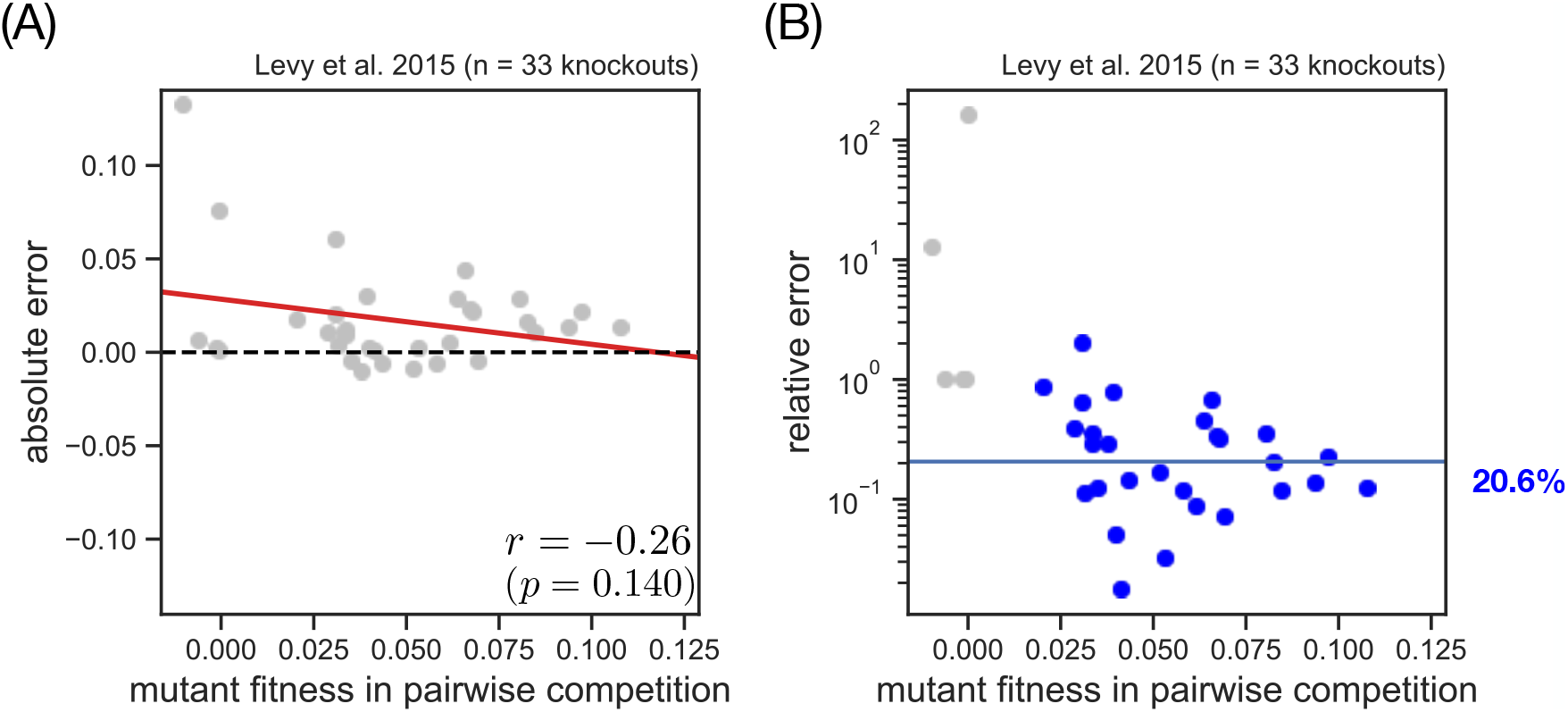
The bulk fitness error based on measurements by Levy et al. [32]. Same as Fig. but for the set of evolved genotypes of *S. cerevisiae* shown in Fig. 2d in the original publication. In (A), we calculate the Pearson correlation coefficient *r* and the p-value for the null hypothesis of zero correlation, and indicate the linear regression between the two variables with a red line. In (B) we define a set of knockouts with non-neutral fitness (blue dots; 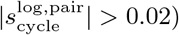 to calculate the geometric mean relative error (blue line). See Fig. 6A for fitness data and experiment conditions.

